# Excitatory neurons downregulate synaptic proteins before motor onset and glia reduce neuronal support in Huntington’s disease mice

**DOI:** 10.64898/2026.11.12.766120

**Authors:** Rodrigo Alvarez-Pardo, Sicheng Luo, Patricia Rodríguez-Rodríguez, Javier Lázaro Sánchez, Héctor Albert-Gascó, Cristina Olmedo-Salinas, Kristina Desch, Max Gubert-Olivé, Josep Muncunill, Miguel Díaz-Hernández, Eugenio F. Fornasiero, Marko Jovanovic, Ella Doron-Mandel, Beatriz Alvarez-Castelao

## Abstract

**Background:** Huntington’s disease burdens protein homeostasis and affects neurons and glia differently across brain regions, but how these two main cell populations respond as the disease progresses remains unresolved. Whole-tissue proteomes mix these cell types and hide how the proteome of each changes. We therefore measured the proteome of each cell type separately at different disease stages.

**Methods:** We metabolically labelled cell-type-specific proteomes in living R6/1 mice and wild-type littermates for 15 days at different disease stages. We examined cortical and hippocampal excitatory neurons at 1.5, 3 and 7 months, cerebellar inhibitory neurons at 3 and 7 months, and cortical, striatal and cerebellar glia at 7 months. We combined mass spectrometry with functional enrichment and competitive protein-set tests to understand the differential proteostatic response to mutant huntingtin across disease progression, brain regions and cell types. Differential abundance was tested with a peptide-centric linear mixed model (msqRob, Benjamini-Hochberg q < 0.1), and proteins whose levels varied together in cortical and hippocampal neurons were grouped into modules by weighted correlation network analysis.

**Results:** Hippocampal excitatory neurons showed broader and more coordinated regulation than cortical neurons already before motor onset, whereas the cortical response increased with age and no cortical protein module differed between genotypes before the most severe stage. All three neuronal populations upregulated ubiquitin–proteasome system (UPS) proteins as the disease advanced. Protein folding changed, with more subunits of the chaperonin CCT/TRiC but fewer J-domain proteins and, in excitatory neurons, fewer HSP70 chaperones.

At the most severe stage, glia of the three studied regions regulated mainly different proteins, but all increased UPS proteins and decreased key proteins of neuronal support, among them the main glial glutamate transporter GLT-1. Cholesterol-related proteins decreased in cortical and cerebellar glia, and respiratory-chain proteins in striatal glia.

**Conclusions:** Synaptic proteins decreased early in cortical and hippocampal neurons and were not restored at later ages. A later and broader maintenance response coexisted with their continued and increasing downregulation. In this first cell-type-resolved view of the labelled proteome in a Huntington’s disease mouse model, progression appears as a change in the coordination of cellular functions, which whole-tissue measurements would dilute.

## Background

Huntington’s disease (HD) is caused by an expanded CAG repeat in the HTT gene, but its molecular consequences differ not only with repeat length but also across cell populations and brain regions. Mutant huntingtin aggregates caused by the polyglutamine expansion place a burden on protein homeostasis. Proteostasis depends on coordinated protein synthesis, folding, assembly, trafficking, modification and removal, in which chaperones, autophagy and the ubiquitin–proteasome system (UPS) play a central role [1]. In HD mice, mutant huntingtin decreases UPS activity at synapses [2]. A subset of chaperone-network genes is expressed less in the brains of patients with Alzheimer’s, Huntington’s and Parkinson’s disease [3]. This evidence links proteostasis to the disease, but whether all these steps change together remains unresolved. Neuronal function depends on the continued maintenance of synaptic proteins, and on metabolic support from neighbouring glia. Cell identity also matters. Astrocytes adopt disease-related states and respond differently by region in HD [4, 5]. A recent time-resolved transcriptomic study in zQ175 mice separated neuronal programmes by disease stage [6]. These studies establish heterogeneity across cell identity and time, but leave open how the protein machinery changes within each population. Protein measurements carry information that transcript abundance does not [7]. Bulk protein measurements, however, mix neighbouring cell populations, so opposite changes in neurons and glia can offset each other, and a change confined to one population can be diluted by the surrounding tissue.

Bio-orthogonal noncanonical amino acid tagging (BONCAT) labels newly synthesised proteins so that they can be recovered from intact tissue. Cell-selective labelling in animals was established first in Drosophila and then in mice, with a modified methionyl-tRNA synthetase, MetRS, which incorporates azidonorleucine (ANL) into newly synthesised proteins [8, 9]. The optimised multicopy 3xMetRS* line labels more efficiently and can be used in heterozygous animals without methionine depletion [10]. We applied this approach, with cell-type-specific Cre-driver lines, to R6/1 mice, which express a human HTT exon-1 transgene with an expanded repeat [11]. Cortical and hippocampal excitatory neurons were examined before motor onset, at early symptoms and at the most severe stage studied, cerebellar inhibitory neurons at the last two stages, and glia from three regions at the most severe stage. We asked two questions. Do later responses reverse the early protein changes? And do the folding, proteasome and metabolic systems respond together across populations? By following individual proteins, pathways and protein sets, we could separate selective regulation of maintenance and catabolic machinery from persistent synaptic changes and from regional responses of glial support proteins.

## RESULTS

We crossed the 3xMetRS* line with neuronal and glial Cre-driver lines, administered ANL for 15 days, and enriched labelled proteins from intact tissue by click chemistry before identifying them by data-independent acquisition mass spectrometry (DIA-MS, Fig. 1a, b) [10]. We studied excitatory neurons in cortex and hippocampus at 1.5, 3 and 7 months, inhibitory neurons in cerebellum at 3 and 7 months, and glia in cortex, striatum and cerebellum at 7 months, comparing R6/1 mice with wild-type (WT) littermates (n = 4 to 6 mice per genotype, Supplementary Table 1, Supplementary File 1).

**Fig. 1.**
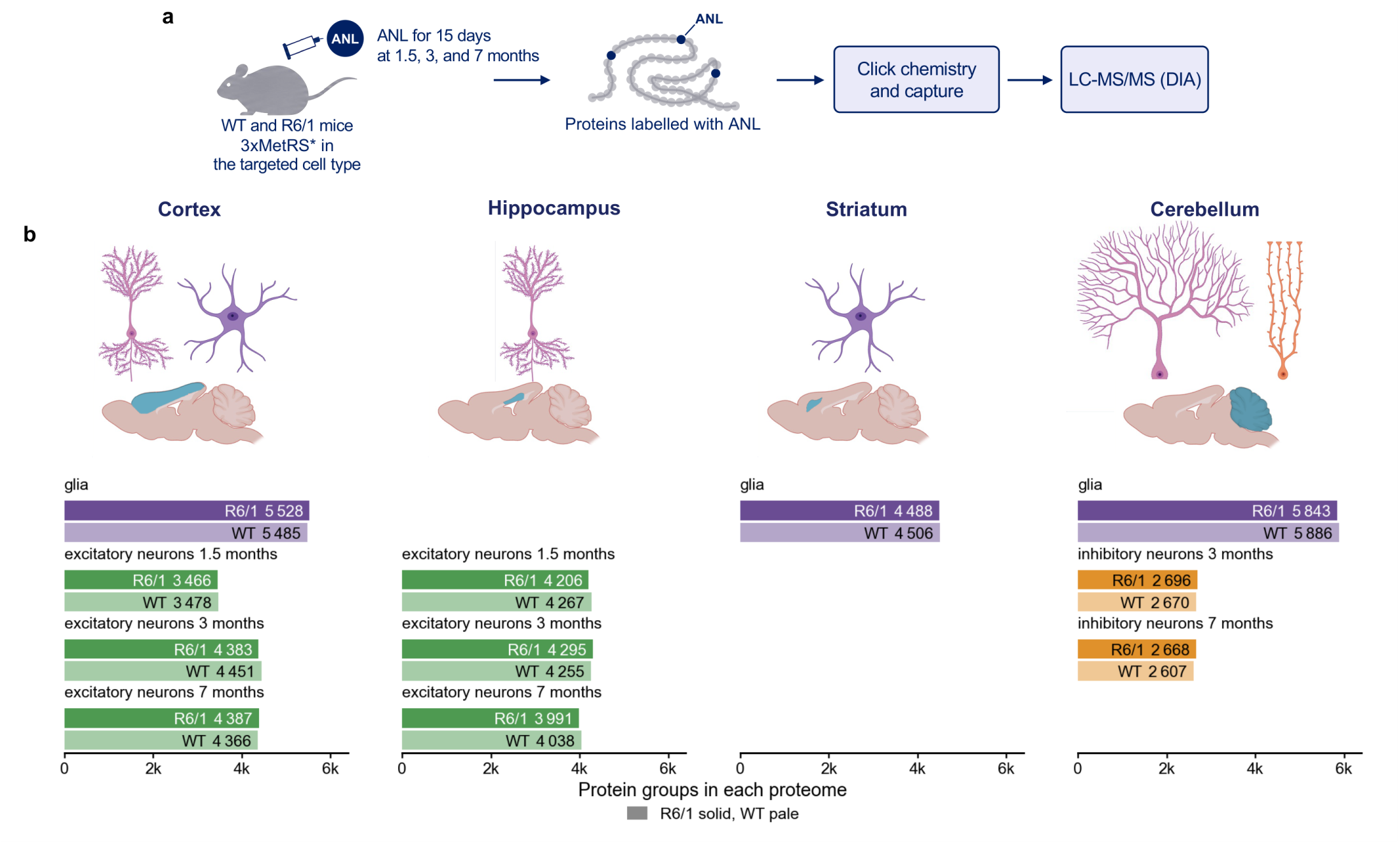
**Cell-type-specific labelling of proteins in R6/1 and WT mice**. **(a)** Experimental workflow. R6/1 mice and WT littermates expressing a mutant methionyl-tRNA synthetase (3xMetRS*) in one cell type (Cre-driven expression) received daily intraperitoneal injections of azidonorleucine (ANL) for 15 days. The proteins of the cell type of interest incorporate ANL, which allows their capture by click chemistry and their identification by liquid chromatography-tandem mass spectrometry in data-independent acquisition mode (LC-MS/MS, DIA). **(b)** Populations studied and number of protein groups identified in each. Excitatory neurons were labelled in cortex and hippocampus at 1.5, 3 and 7 months, inhibitory neurons in cerebellum at 3 and 7 months, and glia in cortex, striatum and cerebellum at 7 months. A protein group was counted in a genotype if it was quantified in both genotypes or found in that genotype only. n = 4 mice per genotype, except for cortical excitatory neurons (6 R6/1 and 6 WT at 1.5 months, 5 and 5 at 3 months, 5 R6/1 and 6 WT at 7 months).

Principal component analysis mostly separated WT and R6/1 samples (Supplementary Fig. 1). Hierarchical clustering grouped the 51 samples at 7 months first by cell type, then largely by region and, in most preparations, by genotype. As expected, replicates of the same preparation and genotype had the highest correlation (median Spearman ρ 0.96, Supplementary Fig. 2a, b, Supplementary Table 2). Because a loss of labelled cells in R6/1 mice could reduce the amount of labelled protein recovered, we compared the total MS signal before normalisation, which did not differ significantly between genotypes in any preparation (Supplementary Fig. 2c).

The amount and direction of differential regulation between WT and R6/1 differed between populations (Fig. 2, Supplementary Fig. 3a). In cortical excitatory neurons, regulation more than tripled with age (211, 390 and 694 regulated protein groups at 1.5, 3 and 7 months). Most proteins were downregulated at 1.5 and 3 months, but more were upregulated at 7 months. Hippocampal excitatory neurons regulated more proteins than cortical ones at every age, peaking at 3 months (665, 2,222 and 1,419 protein groups). Most proteins were downregulated at 1.5 and 7 months and upregulated at 3 months. Cerebellar inhibitory neurons also had broader regulation at 3 than at 7 months (1,458 and 788), with slightly more downregulated proteins at 3 months and nearly balanced directions at 7 months (Fig. 2a).

**Fig. 2.**
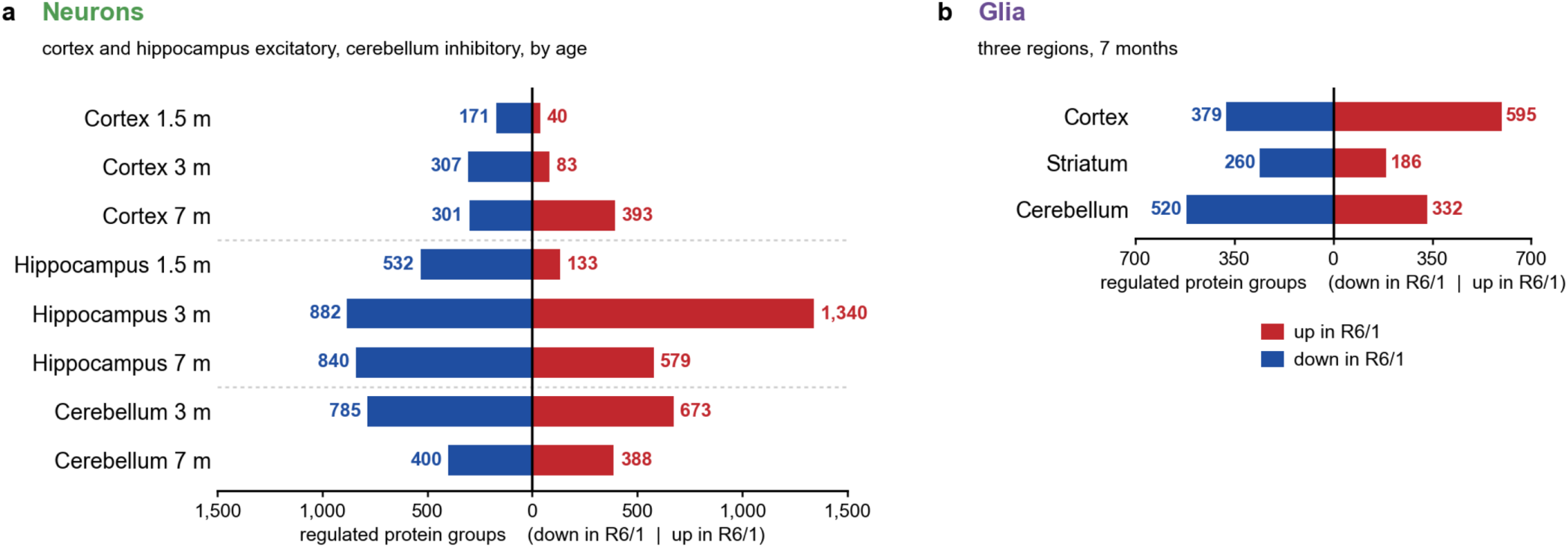
Proteins regulated in R6/1 mice by cell population, region and age. Number of protein groups higher or lower in R6/1 mice than in WT littermates of the same age. **(a)** Excitatory neurons of cortex and hippocampus at 1.5, 3 and 7 months, and inhibitory neurons of cerebellum at 3 and 7 months. **(b)** Glia of cortex, striatum and cerebellum at 7 months. A protein group was considered regulated if its abundance differed between genotypes (msqRob, Benjamini-Hochberg q < 0.10, no fold-change cut-off) or if it was detected in one genotype only (Methods). n = 4 mice per genotype, except for cortical excitatory neurons (6 R6/1 and 6 WT at 1.5 months, 5 and 5 at 3 months, 5 R6/1 and 6 WT at 7 months).

Among glia, when the three studied regions were compared, cortex had the broadest protein regulation, mainly by upregulation, whereas downregulation predominated in striatum and cerebellum (Fig. 2b). Because the number of detected protein groups differed between preparations, depending on how much of each labelled population the tissue contains, we repeated the counts on the groups of proteins quantified in all populations of the same cell type at the same age, and the differences between regions held.

R6/1 fold changes correlated between ages of the same population, most strongly in neurons of cerebellum and hippocampus (Spearman ρ 0.74 to 0.81, Supplementary Fig. 3b), and between glial regions (ρ 0.44 to 0.57, Supplementary Fig. 3c). Almost no regulated protein changed direction between ages (Supplementary Fig. 3d). For a comprehensive study of the data we used Ingenuity Pathway Analysis (IPA), the PANTHER overrepresentation test, weighted correlation network analysis (WGCNA) and the competitive protein-set test cameraPR (limma).

### 1 Cortical and hippocampal excitatory neurons downregulate synaptic proteins before motor onset

To follow the response over time, we compared the proteins regulated in the neurons of each region at each age using Venn diagrams (Supplementary Table 3), and analysed the whole response of each age with IPA, which predicts whether a pathway is up/downregulated (Supplementary Fig. 4, Supplementary Table 4). In the case of the Venn diagrams, each section contains the proteins regulated at only one age, at several ages or at all ages, split by direction. We used the PANTHER overrepresentation test on every section of the Venn diagrams to dissect the differential cellular response among conditions. PANTHER enrichment is reported only for sections that returned enriched terms. A functional annotation absent from the “one condition” sections could, however, still change through proteins of other sections. Additionally, IPA, whose input was all proteins regulated at each age, could detect functions that changed at that age through proteins also regulated at other ages, making the two approaches complementary.

#### 1.1 Cortex

In cortical neurons, most regulated proteins changed at a single age, fewer were shared between two ages, most of them between 3 and 7 months and downregulated, and 23 changed at every age (Fig. 3a).

**Fig. 3.**
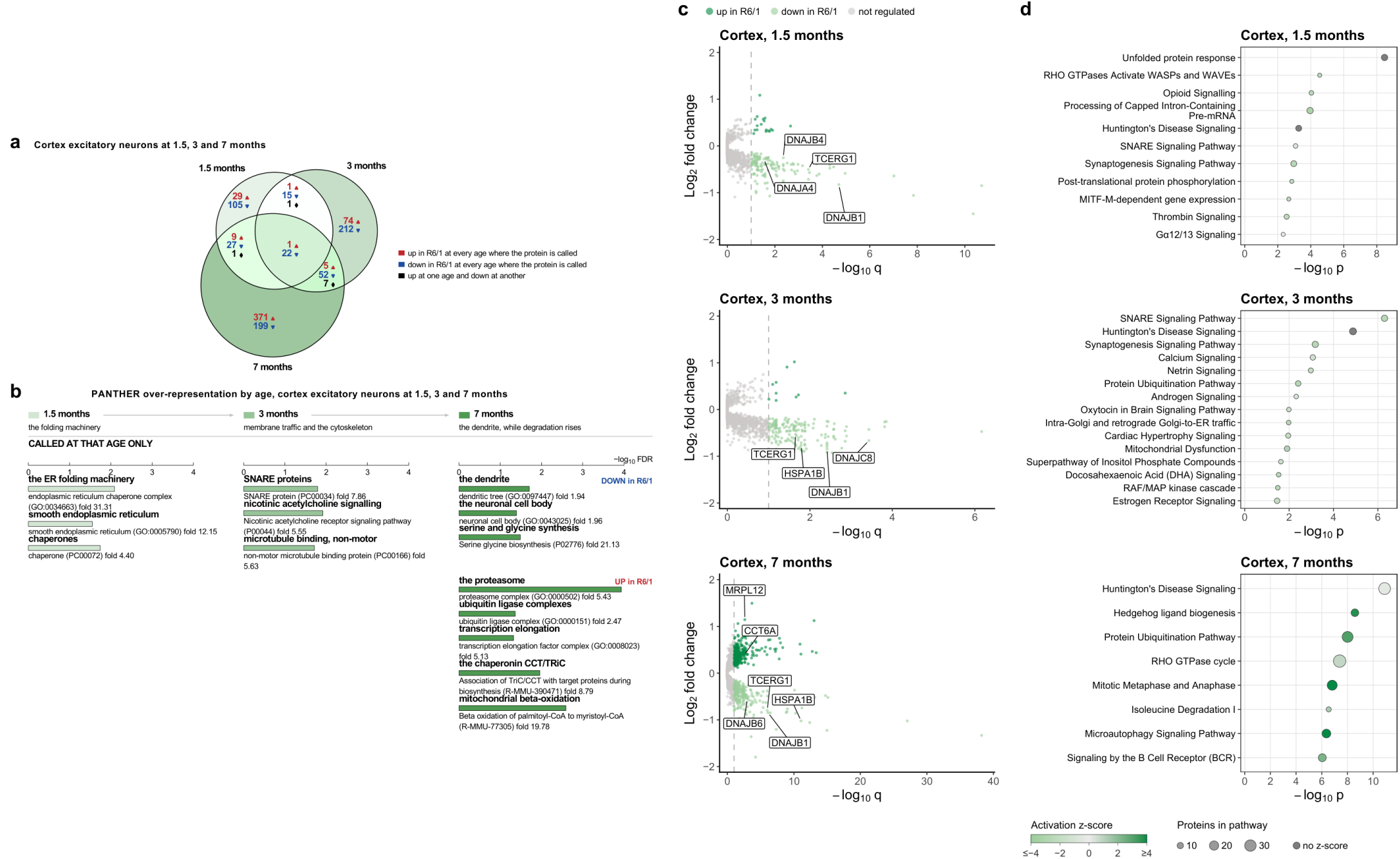
**Protein changes in cortical excitatory neurons of R6/1 mice at 1.5, 3 and 7 months**. **(a)** Venn diagram of the protein groups regulated at each age. In each section, the numbers give the protein groups that were upregulated (red) or downregulated (blue) in R6/1 at a given age, or upregulated at one age and downregulated at another (black). **(b)** Functions over-represented among the proteins regulated at one age only, by direction (PANTHER, Fisher’s exact test with Benjamini-Hochberg correction, FDR < 0.05). Each bar carries a plain name of the function, followed by the PANTHER term, its identifier and its fold enrichment. Bar length is −log10 FDR. The proteins upregulated at 1.5 or 3 months only returned no enriched term. **(c)** Volcano plots of the protein groups quantified in both genotypes at each age, log2 fold change (R6/1/WT) against −log10 q. Regulated proteins (q < 0.10) are coloured as in the key above the plots, the dashed line marks q = 0.10, and selected proteins are labelled. **(d)** IPA canonical representative pathways enriched among the regulated proteins at each age (q < 0.10). Dot size gives the number of regulated proteins in the pathway and colour the activation z-score, as in the key below the plots (dark grey, no z-score computed). n = 6 R6/1 and 6 WT mice at 1.5 months, 5 and 5 at 3 months, and 5 R6/1 and 6 WT at 7 months.

As early as 1.5 months, before motor onset, cortical neurons had fewer chaperones. These were enriched in the downregulated section and included J-domain proteins, which deliver client proteins to heat-shock protein 70 (HSP70) chaperones, and the endoplasmic reticulum (ER) chaperone BiP (Fig. 3a, b).

IPA confirmed the chaperone decrease, with the unfolded protein response as the most significantly changed pathway (9 of its 10 proteins downregulated). Additional downregulated pathways were synaptogenesis, actin pathways, calcium pathways involving calcium/calmodulin-dependent kinases and their target CREB1 (opioid signalling), and pre-mRNA processing including the huntingtin-binding proteins PRPF40A and TCERG1 (IPA, Fig. 3c, d) [12, 13].

At 3 months, at motor onset, cortical neurons had less membrane-fusion machinery, including synaptic, endosomal and Golgi soluble N-ethylmaleimide-sensitive factor attachment protein receptors (SNAREs), and microtubule-binding proteins, both enriched in the downregulated section (Fig. 3a, b). IPA placed SNARE signalling as the most significant downregulated pathway, and showed downregulation of proteins associated with synaptogenesis and the UPS. Synaptogenesis was carried mostly by the same proteins as SNARE signalling, and the protein ubiquitination pathway contained mainly chaperones at this stage (Fig. 3c, d).

At the most severe disease stage studied (7 months), the proteins upregulated only at this stage were enriched in UPS components, transcription elongation factors, subunits of the chaperonin containing TCP-1 (CCT/TRiC), which folds newly made proteins and also inhibits huntingtin aggregation, and mitochondrial beta-oxidation of saturated fatty acids [14]. The downregulated section was enriched in proteins of the dendrite and the neuronal cell body and in serine and glycine synthesis (Fig. 3a, b).

In IPA, the term Huntington’s disease signalling, which was mostly carried by proteasome subunits and chaperones, was the most significant pathway. The increase in UPS was reflected also in the protein ubiquitination pathway, which contains the three parts of the HSP70 disaggregase, HSP70 chaperones, J-domain proteins, and an HSP110 nucleotide-exchange factor [15]. The cytosolic HSP70 chaperones and their two J-domain proteins decreased, whereas the nucleotide-exchange factor HSPA4 increased (Fig. 3c, d, Supplementary Table 4).

The proteins regulated at every age formed a small persistent core (Fig. 3a, Supplementary Table 3). More than half of the downregulated core proteins belonged to synapses and neuronal projections, among them synaptopodin, required for the spine apparatus, and the spine scaffold IRSp53 [16]. The core also contained the J-domain chaperone DNAJB1. ARC, which removes AMPA receptors from the synapse, was less downregulated with age (log2 fold change −1.07, −0.93, −0.69), whereas NPTXR, which binds AMPA receptors and helps build synapses, was more downregulated (−0.85, −0.89, −1.33) (Supplementary Table 1) [17, 18]. Both trends are compatible with fewer AMPA receptors being kept at the synapse.

In summary, these neurons had fewer chaperones and proteins related to RNA processing and synaptic signalling from 1.5 months on, and proteins that fuse and transport membrane vesicles at 3 months. At 7 months, when the regulation was largest, proteasome and chaperonin subunits increased, while HSP70 chaperones and proteins of the dendrite and its actin machinery decreased further, and a small core of synaptic proteins remained low at every age.

#### 1.2 Hippocampus

In hippocampal neurons, unlike in cortical neurons, most proteins that responded as early as 1.5 months sustained their responses at later ages, with 273 proteins remaining regulated across all ages examined (Fig. 4a). In agreement with this observation, PANTHER enrichment analysis on the proteins regulated only at 1.5 months returned no enriched terms. However, the proteins also regulated at later ages yielded enriched terms, among them synaptic terms in those downregulated at 1.5 and 7 months (Supplementary Table 3). At 1.5 months, all members of the neurexin and neuroligin pathway, the most significant in IPA, were downregulated, including the postsynaptic scaffolds PSD-95 (DLG4), SHANK3 and HOMER1, which anchor receptors at the postsynaptic density, and the NMDA-receptor subunit GRIN2A.

**Fig. 4.**
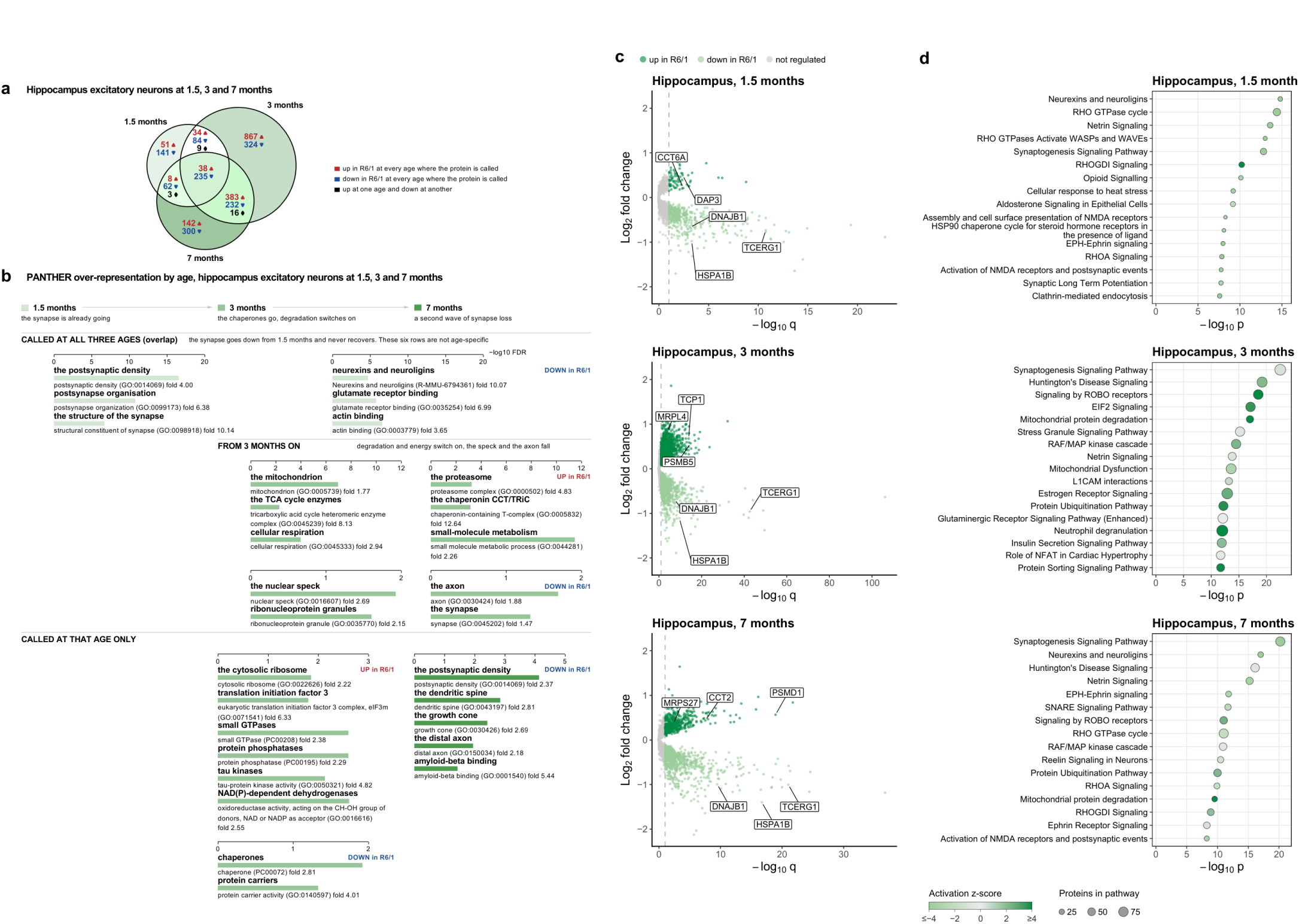
**Protein changes in hippocampal excitatory neurons of R6/1 mice at 1.5, 3 and 7 months**. **(a)** Venn diagram of the protein groups regulated at each age. In each section, the numbers give the protein groups that were upregulated (red) or downregulated (blue) in R6/1 at a given age, or upregulated at one age and downregulated at another (black). **(b)** Functions over-represented among the proteins regulated at all three ages, at 3 and 7 months, and at one age only, by direction (PANTHER, Fisher’s exact test with Benjamini-Hochberg correction, FDR < 0.05, against all proteins identified in the study). Each bar carries a plain name of the function, followed by the PANTHER term, its identifier and its fold enrichment. Bar length is −log10 FDR. The proteins regulated at 1.5 months only, and those upregulated at 7 months only or at all three ages returned no enriched term. **(c)** Volcano plots of the protein groups quantified in both genotypes at each age, log2 fold change (R6/1/WT) against −log10 q. Regulated proteins (q < 0.10) are coloured as in the key above the plots, the dashed line marks q = 0.10, and selected proteins are labelled. **(d)** IPA canonical representative pathways enriched among the regulated proteins at each age (q < 0.10). Dot size gives the number of regulated proteins in the pathway and colour the activation z-score, as in the key below the plots. n = 4 R6/1 and 4 WT mice at each age.

Proteins that promote actin branching in dendritic spines, including WAVE1, N-WASP and Arp2/3-complex subunits, were also downregulated, in the Rho GTPase pathways and in the ephrin and netrin guidance-cue pathways. The downregulated clathrin-mediated endocytosis pathway contained proteins that bend the membrane and recruit dynamin, and dynamin itself, which cuts the vesicle free from the membrane. Finally, three pathways (heat-stress response, HSP90 chaperone cycle and aldosterone signalling) were decreased, carried by downregulated ER and cytosolic chaperones, among them inducible HSP70 chaperones, HSP90 and its co-chaperones and J-domain proteins (Fig. 4c, d). As in cortex, hippocampal neurons had fewer chaperones, and their synaptic changes are compatible with excitatory synapses weakening before motor onset.

From 3 months on, the proteins upregulated at 3 and 7 months were enriched in mitochondrial metabolism, the tricarboxylic acid (TCA) cycle and respiration, in the proteasome, and in the chaperonin CCT/TRiC. Those downregulated at both ages were enriched in nuclear speckles and RNA-protein granules, related to RNA processing and storage, and in proteins tied to neuronal connections.

The upregulated 3-month section was enriched in the translation machinery, the cytosolic ribosome and the initiation factor eIF3, and in small GTPases, many of them RAB proteins of vesicle trafficking between the Golgi, endosomes and the cell surface. It was also enriched in NAD(P)-dependent dehydrogenases of energy metabolism (TCA cycle and lactate production) and redox defence (NADPH synthesis). The downregulated 3-month section was enriched in chaperones, among them mitochondrial chaperones and prefoldin, which transfers newly made proteins to CCT/TRiC and also limits huntingtin aggregation [19]. IPA placed synaptogenesis as the most significant pathway, with members changing in both directions (Fig. 4a-d).

At 7 months, the downregulated section was enriched in the postsynaptic density, dendritic spines, growth cones and distal axons, and in proteins annotated as binding amyloid-beta, such as APOE, clusterin and the prion protein (Fig. 4a, b). IPA again placed synaptogenesis as the most significant downregulated pathway, and SNARE signalling was also downregulated. In contrast, the UPS pathways and mitochondrial protein degradation were upregulated (Fig. 4c, d). Together, these changes are compatible with a second wave of synapse loss.

The persistent set, regulated at every age, was nearly twelve times larger than in cortex. Its downregulated proteins were enriched in the postsynaptic density and in actin binding (Fig. 4b). Over the three ages, the synaptic proteins that increased at 3 months differed from those downregulated at 1.5 months. So the 3-month increase did not reverse the early decrease, which included axonal and presynaptic release proteins at every age and was broadest at 7 months. Two proteasome subunits were already upregulated at 1.5 months and those peaked at 3 and 7 months, whereas respiratory-chain subunits increased only from 3 months on. The inducible HSP70 chaperones and DNAJB1 decreased at every age, most strongly at 7 months, whereas two CCT/TRiC subunits were already up at 1.5 months and all were up at 3 and 7 months.

### 2 Matched protein modules show hippocampal neurons ahead of cortical ones

We next asked which proteins changed together in each region across ages. For this we used WGCNA, which groups proteins whose levels correlate across samples into modules and summarises each module by its eigengene, a single value that represents the level of the whole module [20, 21]. One network was built for each region from every protein present at all three ages, 3,182 proteins from 33 mice for cortex and 3,564 from 24 mice for hippocampus (Fig. 5a). The modules whose eigengene differed between R6/1 and wild-type mice, the genotype modules, are numbered C1 to C5 in cortex and H1 to H6 in hippocampus. They held 855 of the cortical proteins (27%) but 2,617 of the hippocampal ones (73%). In hippocampus, the protein clustering tree divided into two large branches, one with H3 to H6, whose proteins were higher in R6/1 mice, and one with H1 and H2, whose proteins were lower, a split visible at 1.5 months and strongest at 3 and 7 months. The response of the hippocampal network was therefore broader and more coordinated than that of the cortex. As with the regulated proteins, the cortical modules reached significance later than the hippocampal ones. In hippocampus, H1 and H3 differed between genotypes from 1.5 months and the other four modules from 3 months, whereas in cortex C1, C2 and C4 reached significance only at 7 months, and C3 and C5 only with the ages pooled (Fig. 5b).

**Fig. 5.**
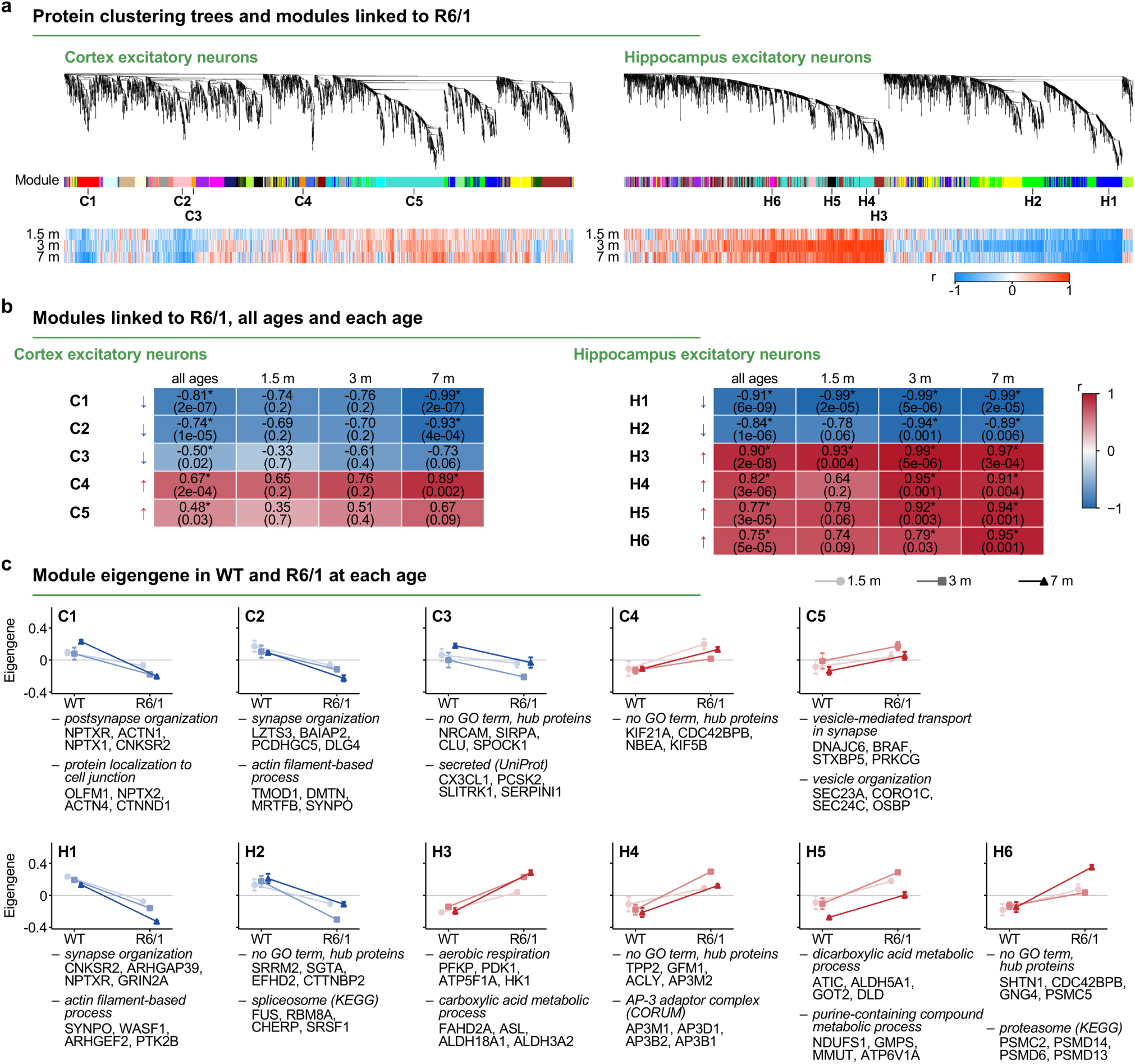
**Protein co-expression modules linked to R6/1 in cortical and hippocampal excitatory neurons**. One network (WGCNA) was built for each region with the three ages together. **(a)** Protein clustering trees. The colour band gives the module of each protein, and the modules linked to R6/1 are numbered. The three rows below show the correlation of each protein with the genotype at each age (blue lower and red higher in R6/1 than in WT). **(b)** Correlation of each module eigengene with genotype, with all ages pooled and at each age (Pearson correlation, Benjamini-Hochberg correction across the modules of the region). Each cell gives r, with q in brackets and an asterisk when q < 0.05. Arrows mark modules lower (blue) or higher (red) in R6/1. A module is linked to R6/1 when q < 0.05 with all ages pooled. **(c)** Module eigengene in WT and R6/1 mice at each age (mean ± s.e.m.), blue when lower and red when higher in R6/1. Under each graph are up to two over-represented GO biological process terms (clusterProfiler, Benjamini-Hochberg adjusted p < 0.05), each followed by four of its proteins not already listed above, those most strongly correlated with the module eigengene. C3, C4, H2, H4 and H6 returned no GO term and show their four hub proteins (highest module membership). For H2 and H6 the most significant of their enriched KEGG pathways is shown. For H4 the AP-3 adaptor complex is shown. For C3 the UniProt keyword secreted is shown. All three were tested with a hypergeometric test (adjusted p < 0.05). All terms and the selection rule are in Supplementary Table 5. n = 4 mice per genotype at each age, except in cortex (6 R6/1 and 6 WT at 1.5 months, 5 and 5 at 3 months, 5 R6/1 and 6 WT at 7 months).

Of the modules lower in R6/1 mice, in cortex C1 and C2 were enriched in synaptic terms, postsynaptic in C1 and also actin in C2 (Fig. 5c, Supplementary Fig. 5). In hippocampus, H1, the module most strongly linked to the R6/1 genotype, was enriched in most of the GO terms of C1 and C2 and held 150 of the 235 hippocampal proteins downregulated at every age (Fig. 4a, Supplementary Tables 3 and 5). H2 was enriched in the spliceosome pathway.

The modules higher in R6/1 mice contained proteins of vesicle traffic, metabolism and the proteasome. In cortex, C5 was enriched in vesicle-mediated transport in the synapse and in vesicle organisation (Fig. 5c, Supplementary Fig. 5). In hippocampus, H3, H5 and H6 matched the mitochondrial, metabolic and proteasome proteins upregulated at 3 and 7 months (Fig. 4b). H4 was the largest hippocampal genotype module. It had no enriched GO term and contained six of the seven subunits of the AP-3 adaptor complex, which sorts membrane proteins from endosomes to lysosomes and to synaptic vesicles.

These modules suggest that, as synaptic proteins decreased, hippocampal neurons had more proteins for energy production and protein degradation, compatible with a higher demand for energy and for the clearance of misfolded proteins. In cortex, the higher vesicle module C5 along with the lower postsynaptic modules is compatible with synapses changing rather than only decreasing in number.

The cortical module C4, which was higher in R6/1, had no enriched GO or KEGG term but appeared to be composed of proteins longer than average. To test this, we compared the C4 proteins with two proteomes of our earlier study, the whole hippocampal proteome and the ANL-labelled proteome of excitatory neurons [9]. The median length of the C4 proteins was about three times that of either proteome, the highest of the 11 modules, and this was not explained by chance (Supplementary Fig. 6a, b). This cortical group partly held in hippocampus, and the eigengene of its proteins was also higher in R6/1 at every age (Supplementary Fig. 6c). Because the C4 proteins changed together and share their length, this increase could reflect a change in how neurons handle some long proteins under the proteostatic stress produced by the expression of mutant huntingtin, rather than a specific functional programme. Many C4 proteins are related to membrane traffic and to the cytoskeleton that supports it.

Seven pairs of cortical and hippocampal modules shared more proteins than expected by chance, and in every pair both modules changed in the same direction in R6/1 (Supplementary Fig. 7a). Together, the networks of the two regions shared a decrease of synaptic and spine proteins and an increase of proteasome subunits and of the AP-3 and COPI vesicle coats (Supplementary Table 5). Only the hippocampal metabolic module H5 had no cortical counterpart, although metabolic enzymes were upregulated in both excitatory populations at 7 months (Supplementary Fig. 8b). All 11 genotype modules were preserved in the network of the other region (Supplementary Fig. 7b). Thus, the two regions changed the same groups of proteins, which also moved together in the other region, compatible with a common response of excitatory neurons that differed mainly in timing and extent.

### 3 From motor onset, cerebellar inhibitory neurons shift protein quality control from HSP70 co-chaperones to the proteasome and CCT/TRiC

In cerebellar inhibitory neurons, 1,037 protein groups were regulated at 3 months only, 367 at 7 months only and 421 at both ages (Fig. 6a).

**Fig. 6.**
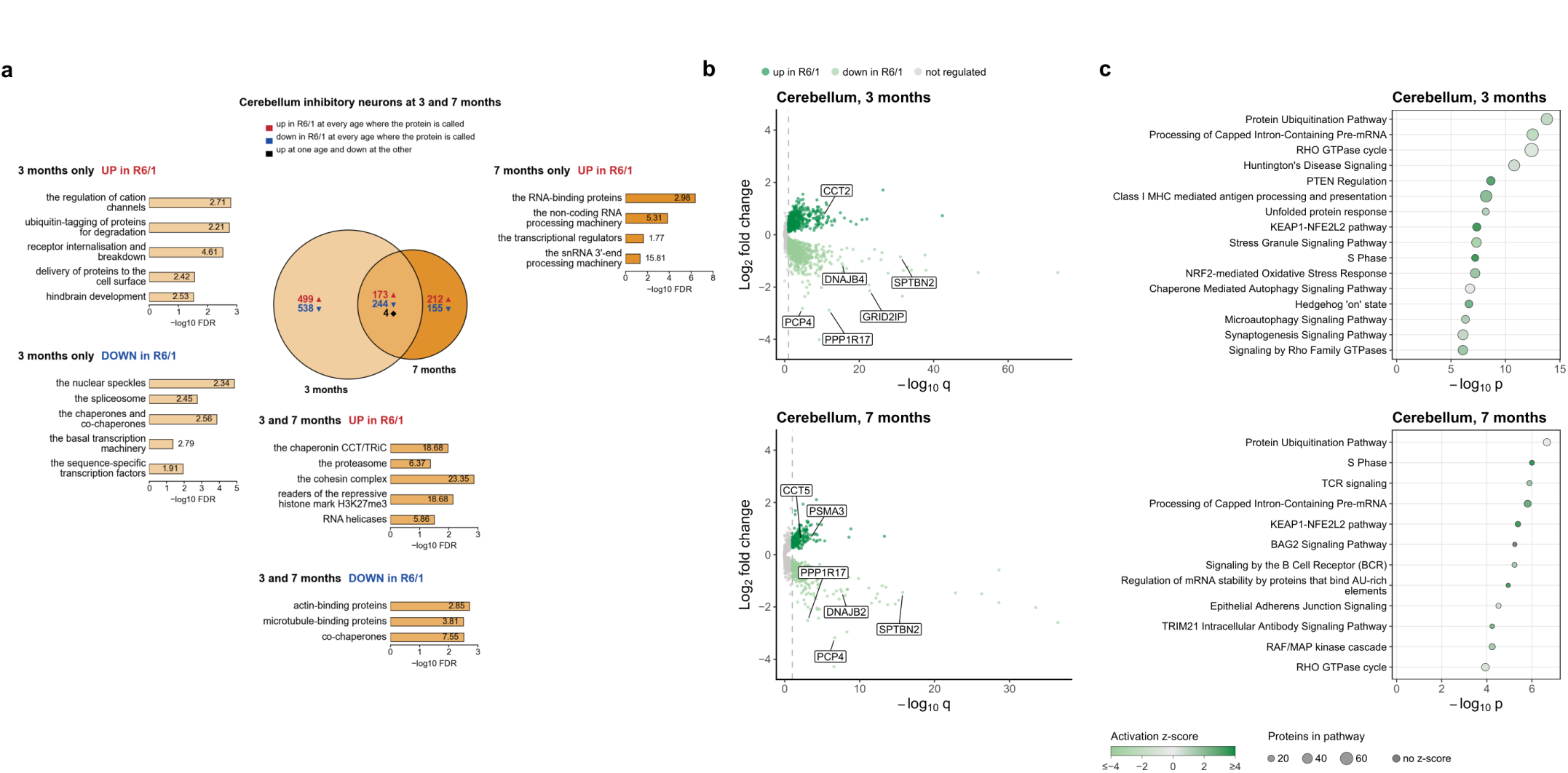
Protein changes in cerebellar inhibitory neurons of R6/1 mice at 3 and 7 months. **(a)** Venn diagram of the protein groups regulated at each age, surrounded by the functions over-represented in its sections. In each section of the Venn diagram, the numbers give the protein groups that were upregulated (red) or downregulated (blue) in R6/1 at a given age, or upregulated at one age and downregulated at the other (black). The functions are those over-represented among the proteins regulated at one age only or at both ages, by direction (PANTHER, Fisher’s exact test with Benjamini-Hochberg correction, FDR < 0.05). Each bar carries a plain name of the function and its fold enrichment. Bar length is −log10 FDR. The proteins downregulated at 7 months only returned no enriched term. **(b)** Volcano plots of the protein groups quantified in both genotypes at each age, log2 fold change (R6/1/WT) against −log10 q. Regulated proteins (q < 0.10) are coloured as in the key above the plots, the dashed line marks q = 0.10, and selected proteins are labelled. **(c)** Representative IPA canonical pathways enriched among the regulated proteins at each age (q < 0.10). Dot size gives the number of regulated proteins in the pathway and colour the activation z-score, as in the key below the plots (dark grey, no z-score computed). n = 4 of each group, R6/1 and WT mice at each age.

At 3 months, the upregulated section was enriched in proteins that control calcium and cation entry, among them L-type calcium channels and NMDA-receptor subunits, which allow calcium influx into the cell, and the ryanodine receptor, which amplifies this influx through calcium-induced calcium release from the endoplasmic reticulum. This section was also enriched in proteins that internalise and break down receptors, many of them endosomal and lysosomal, and in proteins that deliver membrane proteins to the cell surface. UPS proteins also increased, as did synaptic adhesion and axon-guidance molecules, grouped under the term hindbrain development. The downregulated proteins were enriched in RNA production and processing, and in protein folding carried mainly by the same chaperones that decreased in hippocampus, some of them also in cortex (Fig. 6a, Supplementary Table 1).

In IPA, the most significant regulated pathway was protein ubiquitination, with an increase in proteasome subunits and a decrease in chaperones. The downregulated pathways included pre-mRNA processing and stress granule signalling. The term synaptogenesis had members changing in both directions (Fig. 6b, c, Supplementary Table 4). Together, these changes are compatible with a greater turnover of the calcium channels, receptors and other membrane proteins at the surface of cerebellar inhibitory neurons.

At 7 months, the upregulated section was enriched in RNA-binding proteins, among them splicing regulators, in non-coding RNA processing, including the Integrator complex, which matures the small RNAs of the spliceosome, and in transcriptional regulators (Fig. 6a). IPA placed protein ubiquitination as the most significant pathway. Pre-mRNA processing increased, the opposite of 3 months, although no protein of this pathway changed direction. The reversal came from different splicing proteins. Most of those downregulated at 3 months were no longer regulated at 7 months, and others were upregulated instead. Stress granule signalling remained significant. Synaptogenesis, NMDA-receptor postsynaptic events and the neurexin and neuroligin pathway were also significant, mostly with downregulated members, among them PSD-95 (DLG4) (Fig. 6b, c, Supplementary Tables 1 and 4).

The proteins regulated at both ages formed the shared cerebellar response. The shared upregulated ones were enriched in protein folding and turnover, with CCT/TRiC and the proteasome, as in the hippocampal proteins upregulated at 3 and 7 months, and in chromatin organisation. The shared downregulated ones were enriched in cytoskeleton components, with actin- and microtubule-binding proteins, and in HSP70 co-chaperones, as actin-binding proteins and chaperones were in the hippocampal core (Fig. 4b, 6a, Supplementary Table 3).

### 4 Region-specific glial responses converge on downregulation of the glutamate transporter GLT-1

Glia were labelled with the Aldh1l1-Cre/ERT2 line, which in adult mice labels more than 90% of astrocytes in cortex, striatum and hippocampus and no neurons [22]. In this study, glia refers to these labelled cells. In glia most of the regulated proteins changed in a single region, 737 in cortex, 293 in striatum and 666 in cerebellum (Fig. 7a). The upregulated cortical section was enriched for the spliceosome, mRNA export and mRNA stabilisation, and also for the nuclear envelope and lamina and the proteasome regulatory particle (Fig. 7a). In IPA, pre-mRNA processing was the most significant pathway with all of its members upregulated. Actin pathways had members changing in both directions (Fig. 7b, c, Supplementary Table 4).

**Fig. 7.**
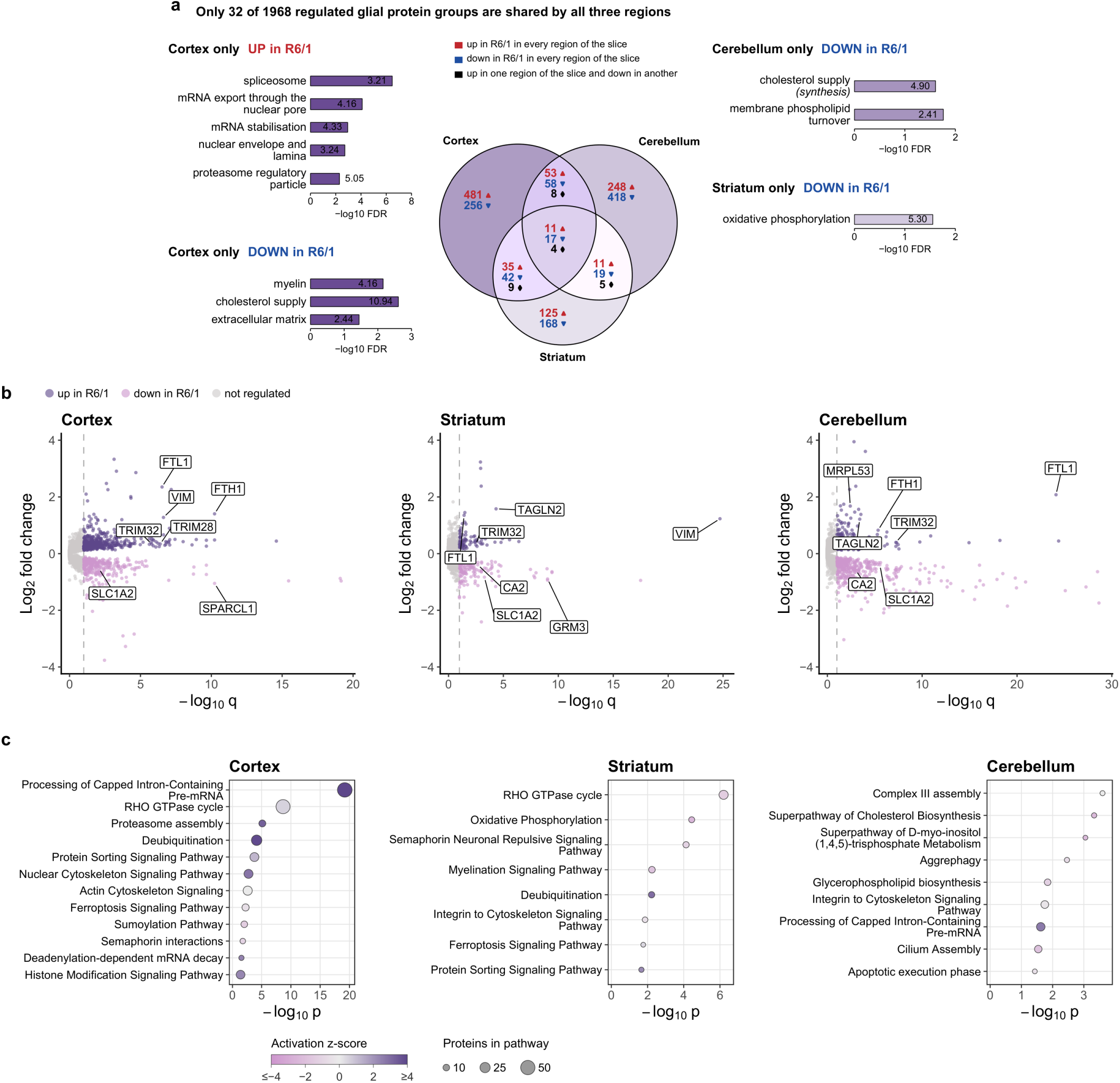
**Protein changes in cortical, striatal and cerebellar glia of R6/1 mice at 7 months**. **(a)** Venn diagram of the protein groups regulated in glial cells of each region, surrounded by the functions over-represented in its sections. In each section of the Venn diagram, the numbers give the protein groups that were upregulated (red) or downregulated (blue) in R6/1 in every region of the section, or upregulated in one region and downregulated in another (black). The functions are those over-represented among the proteins regulated in one region only, by direction (PANTHER, Fisher’s exact test with Benjamini-Hochberg correction, FDR < 0.05). Each bar carries a plain name of the function and its fold enrichment. Proteins upregulated in cerebellum only or in striatum only returned no enriched term. **(b)** Volcano plots of the protein groups quantified in both genotypes in each region, log2 fold change (R6/1/WT) against−log10 q. Proteins upregulated in R6/1 are purple and those downregulated are pink (msqRob, Benjamini-Hochberg q < 0.10, no fold-change cut-off). Grey, not regulated. Representative proteins are labelled. **(c)** IPA canonical representative pathways enriched among the regulated proteins of each region (q < 0.10, proteins quantified in both genotypes). The x-axis is −log10 p (IPA, uncorrected, p < 0.05). Dot size gives the number of regulated proteins in the pathway and colour the activation z-score, as in the key below the plots. n = 4 R6/1 and 4 WT mice per region.

The downregulated cortical section was enriched for cholesterol supply and the extracellular matrix (Fig. 7a). The cholesterol-supply group contained the apolipoproteins APOE and clusterin, which normally carry cholesterol from glia to neurons. The rate-limiting enzyme of cholesterol synthesis was also downregulated (Supplementary Table 1).

In striatal glia, the downregulated section was enriched for oxidative phosphorylation, represented by subunits of respiratory complexes I, III, IV and V (Fig. 7a). In IPA, the RHO GTPase cycle was the most significant pathway. IPA showed upregulation of the UPS and of endosomal protein sorting. Ferroptosis signalling, the iron-dependent form of cell death, was also enriched, and the proteins that supply cysteine for glutathione were downregulated too (Fig. 7c). Together, these changes are compatible with striatal glia producing less energy and being less protected against oxidative damage.

In cerebellar glia, which include the Bergmann glia that wrap Purkinje-cell synapses, the downregulated section was enriched for enzymes of cholesterol synthesis and for membrane phospholipid turnover (Fig. 7a). IPA showed an increase in pre-mRNA processing. Most of the other pathways had members changing in both directions, which points to selective regulation rather than a uniform decrease (Fig. 7c, Supplementary Table 4). Kir4.1 and the gap-junction protein connexin-43, with which Bergmann glia take up the potassium released at Purkinje-cell synapses and spread it through the glial network, were also downregulated (Supplementary Table 1).

We then compared the three glial regions for markers of astrocyte reactivity, whose hallmark is an increase in the intermediate filaments GFAP and vimentin [23]. GFAP did not change in any region, whereas vimentin increased in cortical and striatal glia. Other intermediate filaments also changed by region. Synemin, which is expressed by reactive astrocytes, increased in striatal glia, and nestin, which is reported to respond to injury, increased in cerebellar glia [24, 25]. Most other reactive-astrocyte markers detected did not change either, so the glial response was a partial reactivity, specific to each region, rather than full reactive gliosis.

Finally, 32 proteins were regulated in all three glial regions, 11 upregulated, 17 downregulated and 4 that changed direction between regions, among them vimentin (Fig. 7a, Supplementary Table 3). The upregulated shared proteins included some UPS components and ferritin light chain, which increased strongly in all three regions.

Glia of all three regions downregulated GLT-1, their main transporter for removing the glutamate released by neurons at synapses. The metabotropic glutamate receptor mGluR3, the main type in adult astrocytes, was downregulated as well, and so were subunits of the Na+/K+-ATPase, which takes up potassium, in all three regions (Supplementary Table 1) [26].

Overall, the glial response was largely region-specific. The three regions shared only a small core of proteins, most pathways changed in one or two regions, and even those that changed in every region did so through different proteins or to a different extent. Yet all three converged on more ferritin and UPS proteins and fewer glutamate-uptake and potassium-uptake proteins.

### 5 Beyond region-specific responses, neurons of all three regions upregulate the proteasome and downregulate synaptic calcium signalling

We next compared the three neuronal populations at 7 months, the most severe disease stage studied in all three regions. Most regulated proteins changed in a single region, 392 in cortex, 1,025 in hippocampus, and 567 in cerebellum, whereas 39 changed in all three. Cortical and hippocampal excitatory neurons and cerebellar inhibitory neurons thus shared a small core of proteins related to protein maintenance and synaptic signalling. Proteasome subunits and the chaperonin subunit CCT5 increased, together with proteasome-dependent degradation pathways, whereas DNAJB1, PSD-95 and three kinases that carry calcium signals from the synapse to the nucleus decreased (Supplementary Fig. 8). The proteins that changed in a single region differed in function between regions. In hippocampus, the downregulated proteins were enriched in synaptic transmission, synaptic plasticity and the postsynaptic density, and in transcription regulators, whereas the upregulated proteins were enriched in mitochondrial metabolism, with the TCA cycle, respiration and fatty-acid and sugar breakdown. These changes are compatible with impaired synaptic function and a shift of energy metabolism towards mitochondria. In cerebellum, the upregulated proteins were enriched in transcription initiation, chromatin organisation and mRNA metabolism, in line with altered gene expression. In cortex, only transcription elongation was enriched among the upregulated proteins, suggesting a more diffuse response.

### 6 Neurons and glia respond differently, sometimes in opposite directions

In cortex, 820 proteins were regulated only in glia, and 546 only in excitatory neurons (Fig. 8a). Proteins upregulated exclusively in glia were enriched in RNA processing, in chromatin and nuclear organisation and in protein K63-linked ubiquitination. Glia-specific downregulation pointed to the structures with which glia surround and support neurons, such as the extracellular matrix (Fig. 8a, Supplementary Table 3). Proteins upregulated only in excitatory neurons were enriched in the broad term catalytic complex, including subunits of pyruvate and succinate dehydrogenase from the mitochondria. Those exclusively downregulated in excitatory neurons were enriched in extracellular region, intermediate filament organisation and dendritic and postsynaptic terms, including the neurofilament chains, the spine proteins ARC, HOMER1, DLGAP3, and secreted synaptic organisers. Together with the decrease of matrix proteins in glia, this is compatible with a weaker extracellular environment of synapses, built by both cell types.

**Fig. 8.**
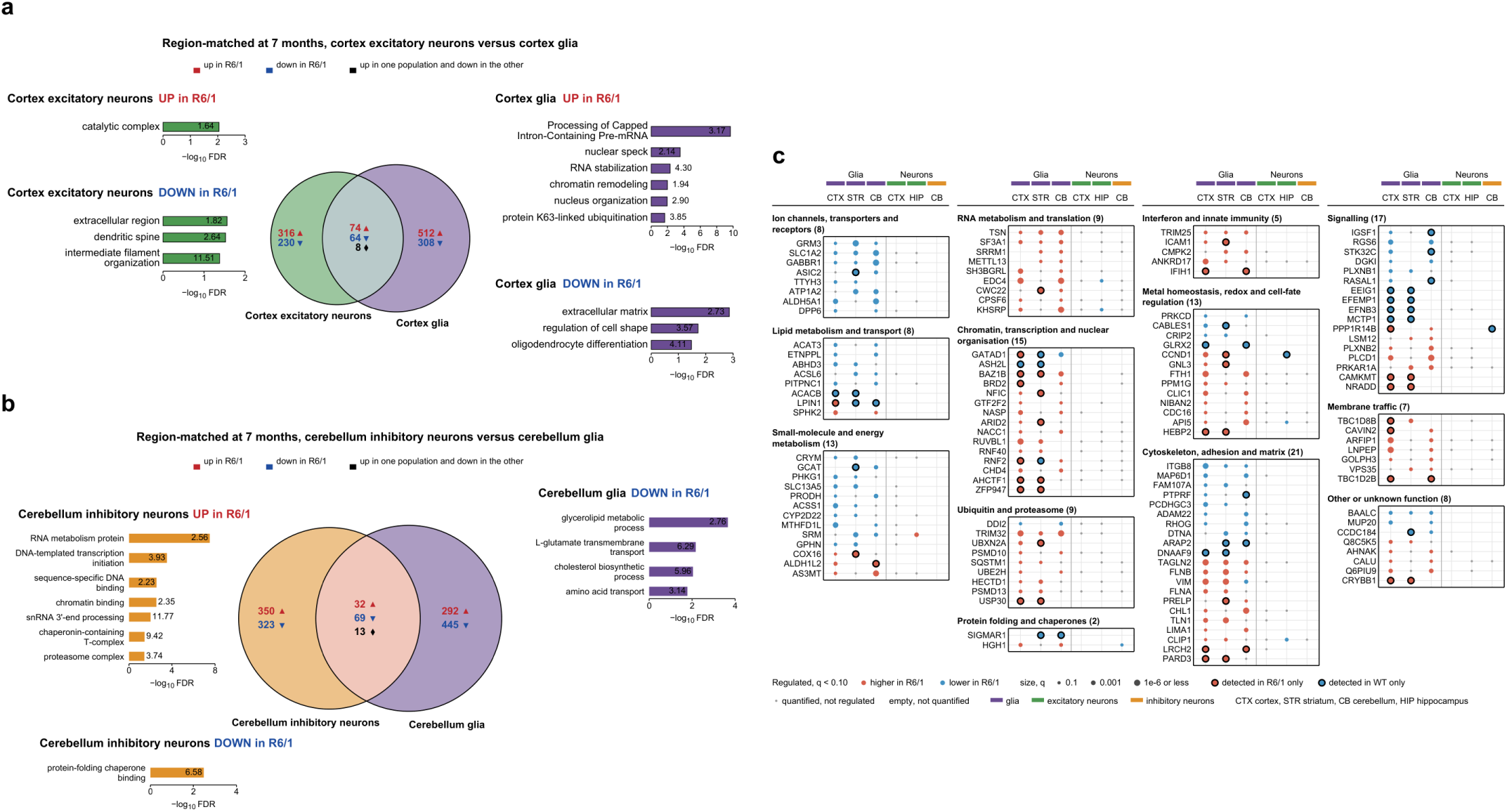
Protein changes in glia compared with neurons of R6/1 mice at 7 months. **(a)** Venn diagram of the protein groups regulated in cortical excitatory neurons and cortical glia, surrounded by the functions over-represented in its sections. In each section, the numbers give the protein groups that were upregulated (red) or downregulated (blue) in R6/1 in every population of the section, or upregulated in one population and downregulated in the other (black). The shown functions are the ones over-represented among the proteins regulated only in a given direction in one population (PANTHER, Fisher’s exact test with Benjamini-Hochberg correction, FDR < 0.05). Each bar carries the PANTHER term and its fold enrichment. **(b)** The same for cerebellar inhibitory neurons and cerebellar glia. The bar protein-folding chaperone binding is a PANTHER GO-Slim molecular function term. Proteins upregulated in glia only returned no enriched term. **(c)** Glial proteins regulated in the same direction in two or three brain regions and not in the same direction in any neuronal population at 7 months. Columns are glia of cortex (CTX), striatum (STR) and cerebellum (CB), excitatory neurons of cortex and hippocampus (HIP) and inhibitory neurons of cerebellum. Proteins are grouped by UniProt function. Circles are red when higher and blue when lower in R6/1, and their area increases with −log10 q. Black ring, detected in one genotype only. Grey dot, quantified and not regulated. Empty, not quantified. A circle of the other colour in a neuronal column marks a change in the opposite direction. n = 4 mice per genotype, except for cortical excitatory neurons (5 R6/1 and 6 WT).

In cerebellum, 737 proteins were regulated only in glia, and 673 only in inhibitory neurons (Fig. 8b, Supplementary Table 3). Proteins downregulated only in glia pointed to functions with which glia support neurons, such as cholesterol synthesis and glycerolipid metabolism, and included the glutamate transporter GLT-1 and the potassium channel Kir4.1. Proteins upregulated specifically in inhibitory neurons were enriched in the chaperonin CCT/TRiC and the proteasome, whereas those downregulated only in inhibitory neurons were enriched in protein-folding chaperone binding, mostly HSP70 co-chaperones of the DNAJ and BAG families. Together, this is compatible with inhibitory neurons mainly adjusting their own protein quality control, whereas glia reduce part of this support.

We then compared the proteins regulated at 7 months in all neuronal and all glial populations together. More than four in five regulated proteins changed in only one cell type, 1,779 in neurons and 1,318 in glia, whereas 638 changed in both (Supplementary Fig. 9). Proteins upregulated in both excitatory neurons and glia were enriched in UPS proteins, although only partly the same ones, whereas spliceosome proteins moved in opposite directions, up in glia and down in hippocampal excitatory neurons (Supplementary Fig. 10).

In the two excitatory populations, 143 proteins changed in the same direction and in no glia, including upregulated V-ATPase proton-pump subunits (Supplementary Table 6). To understand how glia of the different regions change together at this stage while neurons do not, we analysed the proteins regulated in the same direction in at least two glial regions and not in the same direction in any neuronal population at the same age. Most of the decreased proteins were related to cytoskeleton, adhesion and matrix proteins, ion channels, transporters and receptors, and proteins of energy and lipid metabolism, whereas the increased ones included chromatin, transcription, RNA-processing, ubiquitin–proteasome and interferon and innate-immunity proteins (Fig. 8c). Cortical and cerebellar glia had more ferritin heavy chain (Fig. 8c) and cortical glia less transferrin receptor 1, whereas hippocampal neurons had less ferritin heavy chain at 1.5 and 3 months (Supplementary Table 1).

To ask which changes are unique to glia across the disease course, we focused on proteins regulated in the same direction in all three glial regions and in no neuronal population at any age. These 9 proteins comprised GLT-1, mGluR3, the adhesion receptor integrin β8 and BAALC, a poorly characterised protein, which were downregulated, and the actin-binding proteins filamin B and transgelin-2, the chromatin remodelling protein BAZ1B, the RNA-binding protein translin and the antiviral ubiquitin ligase TRIM25, which were upregulated (Supplementary Table 6).

### 7 Glia show partial activation, while only neurons remodel their protein-folding machinery

We next asked, in a competitive set analysis (cameraPR in limma), whether selected functional protein sets (based mainly on Gene Ontology terms, GO) shifted together relative to the remaining quantified proteins in each comparison (Fig. 9, Supplementary File 2 and Supplementary Table 7) [27, 28]. CameraPR has no threshold on single proteins. It can therefore detect small, coordinated changes that single-protein calls miss. Because interferon and antiviral proteins were upregulated in glia across regions (Fig. 8c), we first asked whether glia mount an inflammatory response, compatible with a reaction to damaged neurons. A GO set of proteins that restrict viral replication was upregulated in glia and not in neurons. Additionally, we tested genes enriched in interferon-responsive astrocytes, a reactive state defined in mouse cortex after an inflammatory stimulus [29]. As expected for an astrocyte set, few of its members were quantified in neurons. In glia it was upregulated in striatum and cerebellum (Fig. 9a, Supplementary File 2a).

**Fig. 9.**
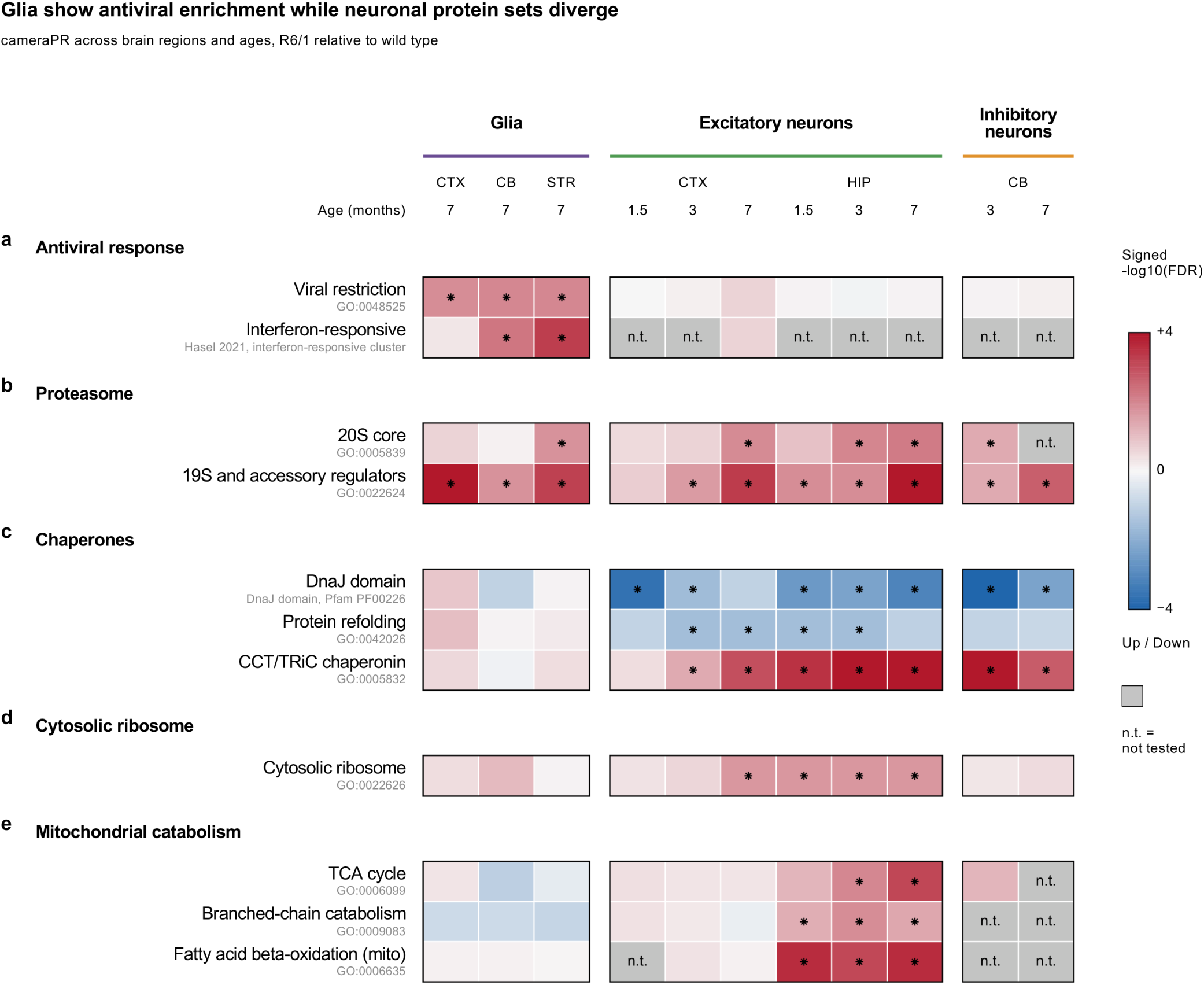
Protein-set tests in glia and neurons of R6/1 mice. Each cell tests whether the proteins of one set, as a group, were upregulated or downregulated in R6/1 compared with all other quantified proteins for a given condition (competitive set test, cameraPR). Each tested set comes from an annotated Gene Ontology term (mouse UniProt), except the DnaJ set (Pfam PF00226), and the beta-oxidation set was limited to mitochondrial proteins. For the interferon-responsive set, we used an astrocyte cluster responsive to IFN identified by Hasel et al. from single-cell brain data [29]. The source of each set is written in grey under its row, and all members are in Supplementary Table 7. **(a)** Antiviral response. **(b)** Proteasome. The 19S and accessory regulators include the PA28 activators. **(c)** Chaperones. **(d)** Cytosolic ribosome. **(e)** Mitochondrial catabolism. The cameraPR p-value of each cell was corrected with Benjamini-Hochberg false discovery rate (FDR). Colour gives the signed −log10 FDR, red when the set is upregulated in R6/1 and blue when it is downregulated, as in the scale bar. Asterisks mark FDR < 0.05. Grey cells (n.t.) had fewer than five set members in the test and were not tested. The proteins of each set are shown in Supplementary File 2. n = 4 mice per genotype, except for cortical excitatory neurons (6 R6/1 and 6 WT at 1.5 months, 5 and 5 at 3 months, 5 R6/1 and 6 WT at 7 months).

Based on the previous results, we analysed the proteasome distinguishing the 20S core and the 19S and accessory regulators (Fig. 9b, Supplementary File 2b). Overall, the regulation was similar, and both sets followed disease severity. However, the regulatory particle increased earlier and more widely than the core. The 19S and accessory set was significant in hippocampal neurons at every age, in cortical and cerebellar neurons from 3 months and in glia of all three regions, whereas among glia the 20S core was significant only in striatum.

We then tested three chaperone sets (Fig. 9c, Supplementary File 2c). Among them were the J-domain proteins, which deliver client proteins to HSP70, and the protein-refolding set, which consists mostly of HSP70-family chaperones and small heat-shock proteins. Overall, in neurons the J-domain set decreased and the CCT/TRiC set increased, the protein-refolding set decreased only in excitatory neurons, and none of the three sets reached significance in glia. Unlike the proteasome, these decreases did not consistently progress with disease severity.

Because the cytosolic ribosome was enriched among proteins upregulated in hippocampal neurons at 3 months (Fig. 4b), we then tested this set in all populations (Fig. 9d, Supplementary File 2d). Overall, the set increased modestly in hippocampal neurons at every age and in cortical neurons only at 7 months, and did not reach significance in cerebellar neurons or in glia.

We then tested three linked sets of mitochondrial fuel use (Fig. 9e, Supplementary File 2e). Overall, the three sets increased only in hippocampal neurons, and none reached significance in glia or in cortical neurons. Branched-chain amino acid catabolism and beta-oxidation were upregulated at every hippocampal age, whereas the TCA-cycle set was upregulated at 3 and 7 months and its significance followed disease severity, so the two pathways that supply the TCA cycle increased before the cycle itself.

## DISCUSSION

To our knowledge, we report the first metabolically labelled proteomes of specific cell types in an HD mouse model, following neurons from before motor onset to advanced disease. In R6/1 mice, cortical and hippocampal excitatory neurons downregulated synaptic proteins at the earliest stage studied, and these proteins remained low while later ages showed additional changes in other proteins. The proteasome increased in all populations, whereas the folding and metabolic systems did not respond in a concerted manner. Most proteins changed in only one cell type, but some changed in opposite directions. Only cell-type resolution reveals this, because a whole-tissue measurement would dilute most of these changes and cancel part of them.

### Synaptic proteins and calcium signalling

Synaptic proteins were already downregulated in hippocampal and cortical excitatory neurons before motor onset, when R6/1 hippocampus lacks inclusions and before the onset of the cognitive and motor deficits that appear at about 3 months, so their decrease may be one of the first steps of the disease in these neurons [30, 31]. The labelled proteins came from the excitatory neurons themselves, and their total MS signal did not differ significantly between genotypes (Supplementary Fig. 2c), so this early decrease is unlikely to reflect fewer labelled neurons or a contribution of other cell types. Synaptic genes, proteins and synapses were also reduced in the cortex and striatum of patients, suggesting that these changes are relevant to human disease [32–34].

Hippocampal neurons had less PSD-95 and GluR1, whereas synaptophysin and gephyrin did not decrease (Supplementary Table 1), the same pattern reported before in R6/1 tissue, where memory was impaired before motor symptoms [35]. The kinase Pyk2, which binds PSD-95 and is needed to cluster it at synapses and to keep spine density, was low in these neurons at every age (Supplementary Table 1). Pyk2 is also lower in the hippocampus of late-stage patients, and increasing it in R6/1 hippocampus at 4–5 months restored memory and partly restored spines, so this early decrease may contribute to memory deficits [36]. PSD-95, in contrast, was higher in post-mortem patient hippocampus homogenates [33].

On the presynaptic side, cortical neurons had lower levels of SNAP-25 and rabphilin 3A but not of syntaxin 1A, as HD frontal cortex had lower levels of SNAP-25 and rabphilin 3A but unchanged synaptobrevin II, syntaxin 1 and synaptophysin, which pointed to a defect in transmitter release rather than a loss of synapses [37]. Hippocampal neurons downregulated more proteins related to vesicle release, among them complexin II, although it was higher in the hippocampus of patients [38]. Our neurons already had less rabphilin 3A and bassoon at 1.5 months (Supplementary Table 1), well before bassoon is drawn into huntingtin inclusions from 16 weeks, and their mRNAs were lower in R6/1 cortex [39, 40].

Kinases that carry calcium signals to the nucleus were also decreased. CaMKK2 and the two kinases it activates, CaMKIV and CaMKIδ, were downregulated in neurons of all three regions at 7 months (Supplementary Fig. 8c). In cerebellar inhibitory neurons, these kinases and STIM1, which keeps calcium in the endoplasmic reticulum of Purkinje cells, were already lower at 3 months, while proteins that let calcium into the cytosol, among them RyR2, increased (Fig. 6a, Supplementary Table 1), a disruption of calcium signalling of the kind proposed to start Purkinje-cell dysfunction in spinocerebellar ataxias [41, 42]. These neurons also showed less of the Purkinje-cell proteins PCP2 and PPP1R17 at both ages and of phospholipase Cβ3 at 7 months (Supplementary Table 1), as Pcp2 and Plcb3 mRNAs were lower in laser-captured Purkinje cells of R6/2 and SCA7 mice [43].

### Glial support of synapses

Neurons downregulated proteins involved in synaptic communication, while glia had fewer proteins that support it. This extends to cell-type-specific proteins the evidence, mostly from transcripts, that astrocytes lose normal functions in HD [4, 44]. Glia of all three regions decreased the glutamate transporter GLT-1 and the glutamate receptor mGluR3 (Fig. 8c). GLT-1 also decreases in the striatum of HD patients as the disease advances [45]. Glia of all three regions also downregulated potassium-uptake proteins, and cerebellar glia Kir4.1 and connexin-43 (Supplementary Table 1). In R6/2 striatal astrocytes, restoring Kir4.1 normalised extracellular potassium and prolonged survival, although Kir4.1 did not change in R6/1 striatal glia [46]. In cerebellar glia, lower tubulins and vimentin with more nestin may point to a change in the shape of Bergmann glia (Supplementary Table 1). Cerebellar glia decreased cholesterol-synthesis enzymes, and cortical glia the rate-limiting enzyme HMG-CoA reductase (Fig. 7a, 8b, Supplementary Table 1), in line with lower cholesterol-synthesis genes in astrocytes of HD patient cingulate cortex [4]. Neurons needed glial cholesterol to form many mature synapses, and astrocytes with an expanded CAG repeat secreted less of it and failed to support the synaptic activity of mutant neurons, so less glial cholesterol could contribute to the neuronal decrease in synaptic proteins [47, 48].

In cortex, glia downregulated several matrix proteoglycans and secreted proteins that control the formation of excitatory synapses, while excitatory neurons had fewer secreted synaptic organisers (Fig. 8a, Supplementary Table 1) [49]. Both cell types may thus weaken the extracellular environment of synapses, in line with astrocytes in the striatum of HD patients, which upregulated a signal for matrix degradation [50].

Glia of all three regions, but no neuronal population, also downregulated integrin β8 and BAALC (Fig. 8c, Supplementary Table 1). In adult rodent brain, integrin β8 was found at synapses and in glial processes [51]. Downregulating glial integrins reduced nervous-system dysfunction in a Drosophila model of HD, so the decrease of integrin β8 found here could be protective [52]. Together, these changes take the glial interpretation beyond inflammatory markers to proteins that support synapses and brain metabolism.

### Protein maintenance in neurons and glia

In neurons, the protein-maintenance machinery did not change as one unit. Neurons increased CCT/TRiC and decreased J-domain proteins and, in cortex and hippocampus, the protein-refolding set (Fig. 9c). Our pattern agrees with R6/2 mice, which also express HTT exon 1 and had more CCT/TRiC and proteasome accessory proteins in the soluble proteome of whole brain regions [53]. In R6/2 brain, DNAJB1 protein decreased while its transcript did not, which pointed to sequestration or faster turnover [54]. Faster turnover could also explain why DNAJB1 was lower in all our neuronal comparisons, because we measured proteins labelled and kept over 15 days.

This imbalance matters for aggregation, because the J-domain proteins that decreased are needed by the machines that act on huntingtin aggregates. In vitro and in yeast, CCT/TRiC cooperated with the HSP70 system to turn polyglutamine-expanded huntingtin fragments into soluble oligomers [55]. DNAJB1 was the rate-limiting partner of Hsc70 and HSPA4 in suppressing and reversing huntingtin exon 1 aggregation in vitro, and the 19S particle with a DNAJB6-HSP70-HSP110 module fragmented aggregates for autophagy and delayed huntingtin inclusions [56, 57]. We found downregulation of DNAJB1 and DNAJB6 but upregulation of HSPA4 (Supplementary Table 1) and the 19S proteasome (Fig. 9b), so both machines may have lacked their J-domain partner. A second limit may lie in the 20S proteasome core. In vitro, oligomers of huntingtin and other disease-associated proteins impaired its gate and stopped the 19S from injecting substrates, so more 19S could face a core that admits less [58]. Cortical and hippocampal neurons upregulated the 19S before the 20S (Fig. 9b), a sequence compatible with this hypothesis.

In HD patients, proteasome function is reduced in cortex, cerebellum, substantia nigra and caudate-putamen from early stages, and patient skin fibroblasts show this reduction despite an increased expression of proteasome beta subunits [59]. More proteasome subunits therefore do not necessarily imply more degradation and could instead represent a compensatory attempt to build more proteasomes in response to impaired degradation. Module C4, higher in R6/1 neurons, was composed of unusually long proteins (Supplementary Fig. 6), which may point to a regulation that depends on protein length. Long proteins could be particularly sensitive to proteostatic stress, as amyloid-like aggregates in human cells sequestered relatively large proteins rich in unstructured regions [60].

Glia differed from neurons in their chaperones rather than in their proteasome subunits. Both upregulated the regulatory particle of the proteasome, but only neurons downregulated the co-chaperones that deliver misfolded proteins to HSP70 and the proteins that refold them (Fig. 9b, c), and hippocampal and cerebellar neurons also had less of the ubiquitin ligase that sends chaperone-bound proteins to the proteasome (Supplementary Table 1). This could reflect a larger capacity of glia to refold or degrade misfolded proteins, including mutant huntingtin. Cultured astrocytes clear mutant huntingtin faster than neurons, with higher proteasome activity, more and more inducible Hsp70, and a more active form of that ubiquitin ligase, and astrocytes, oligodendrocytes and microglia of HD mice form fewer and smaller nuclear inclusions than neurons [61–64]. In our study glia have sparse inclusions even at the more severe disease stage (data not shown). Thus, part of the glial response may come from mutant huntingtin taken up from neurons, as glia engulf neuronal huntingtin aggregates in the fly brain [65]. Vimentin increased in cortical and striatal glia. Because vimentin forms a cage around aggregated proteins when the proteasome is overloaded, this increase could help glia cope with mutant huntingtin [66].

Glia did not show a full inflammatory reaction. They increased interferon and antiviral proteins across regions, as single proteins (Fig. 8c) and as sets (Fig. 9a), in line with the innate immune signalling reported in glial cells of HD patients [67]. Innate immune signalling was also reported in striatal projection neurons, whereas in our data the viral-restriction set was upregulated only in glia [68].

### Ribosomal proteins and RNA splicing

The cytosolic-ribosome set increased modestly in hippocampal neurons at every stage studied and in cortical neurons at the most severe stage (Fig. 9d). This matches R6/1 mice, in which ribosomal proteins were synthesised at a higher rate in the whole brain and were more abundant in the striatum, and contrasts with the repression of protein synthesis by mutant huntingtin, which stalled ribosomes in a mouse striatal cell line [69, 70]. More ribosomal protein expression does not mean more functional ribosomes, nor more translation, however.

Generally, splicing proteins changed in both cell types, but differently. In neurons, pre-mRNA processing was downregulated in all three regions by 3 months (Supplementary Fig. 4) and increased again in cerebellar inhibitory neurons at 7 months, whereas in cortical and cerebellar glia, splicing proteins were upregulated at 7 months (Fig. 7a, c). The striatum of HD patients and of early symptomatic R6/1 mice shares mis-spliced genes and lower levels of splicing factors [71]. The transcription coactivator TCERG1, which binds huntingtin and links transcription to mRNA processing, decreased in every neuronal comparison and in cerebellar glia (Supplementary Table 1) [13, 72]. Longer repeats in its gene brought forward motor onset in HD patients, which makes it a candidate link between transcript processing and disease progression [73].

### Energy and redox metabolism

Hippocampal neurons changed their metabolism most consistently, upregulating the pathways that fuel mitochondria. In these neurons, branched-chain amino acid catabolism and fatty-acid beta-oxidation, which feed the TCA cycle, were first upregulated, and the cycle itself and the respiratory proteins followed (Fig. 4b, 9e), so the response extended as the disease progressed. Cortical neurons at 7 months increased SDHA, the catalytic subunit of succinate dehydrogenase, the enzyme shared by the TCA cycle and the respiratory chain (Supplementary Table 1). These increases are not what an energy failure would predict, and they do not contradict the respiratory defects of patients, which were restricted to the basal ganglia and absent from cortex and cerebellum [74]. Succinate dehydrogenase activity was also higher in the cortex of HD patients, and low plasma branched-chain amino acids in patients were proposed to reflect a need of the brain for TCA-cycle fuels, a proposal that fits the increased abundance of proteins of this pathway in our data from hippocampal neurons [75, 76]. Neurons were long thought to use little fatty-acid beta-oxidation, but nerve terminals can indeed oxidise fatty acids from lipid droplets to make ATP [77, 78]. Glia also had more ferritin, the iron-storage protein (Fig. 8c), and cortical glia less transferrin receptor, which takes up iron (Supplementary Table 1). Less uptake and more storage is the expected response to excess free iron, as in the brains of HD patients, where microglia accumulate ferritin and iron [79]. Striatal glia downregulated respiratory-chain subunits and the proteins that supply cysteine for glutathione (Fig. 7a, c), which may leave them less protected against oxidative damage. Complexes III and IV showed lower activity in the caudate of HD patients, and blocking cystine uptake leads to ferroptosis, whose signalling was enriched in these glia [80, 81].

### Progression in cortical and hippocampal neurons

Cortical and hippocampal excitatory neurons shared one network response that started earlier and was broader in hippocampus (Fig. 5, Supplementary Fig. 7). The broader hippocampal and later cortical response is consistent with heterozygous Q175 mice, whose hippocampal tissue had more altered microRNAs than cortical tissue at 6 and 10 months and kept more of them [82]. In the same knock-in line, single-nucleus RNA sequencing of motor cortex showed that excitatory neurons changed little at the early symptomatic stage and more at the late one [6]. Our data from hippocampal neurons already showed many regulated proteins at 1.5 months, whereas R6/1 hippocampal tissue at 8 weeks, before motor onset, showed far fewer phosphorylation changes than striatum [83]. Mees and colleagues proposed that signals from the many cell types of the hippocampus might cancel each other out.

### Outlook

The early and persistent decrease in synaptic proteins and actin regulators in R6/1 excitatory neurons has a parallel in patients with Alzheimer’s disease, whose brains show lower levels of both protein groups from early stages through advanced disease [84, 85]. Our results raise a broader question for neurodegenerative diseases, whether more maintenance machinery preserves the functions affected first or coexists with their persistent loss. In a TDP-43 mouse model, protein-folding factors increased only transiently before disease onset, whereas the chaperonin and 19S increases of R6/1 neurons persisted while the synaptic proteins downregulated first remained low along disease progression [86]. The HSP70 disaggregase, whose J-domain partner decreased in R6/1 neurons, also dismantles α-synuclein fibrils in vitro, so its decrease could matter in other diseases with protein aggregates [87]. These cell-type-specific proteomes also provide a reference for other mouse models of HD, for human tissue and for transcript studies.

### Conclusions

Following the labelled proteomes of identified neuronal populations across disease stages, and of glia at the most severe stage, showed that the populations of R6/1 mice differed in their responses, and that hippocampal neurons responded earlier and more broadly than cortical neurons. In excitatory neurons, synaptic proteins were downregulated before motor onset and were not restored at later stages, even as the chaperonin CCT/TRiC, the proteasome regulatory particle, ribosomal proteins and mitochondrial catabolic enzymes increased in selected populations. Increases in parts of the maintenance and catabolic machinery thus coexisted with the early synaptic deficit but did not reverse it. Glia of all three regions upregulated proteasome proteins and had fewer proteins of glutamate and potassium uptake, and cortical and cerebellar glia also those of cholesterol synthesis, pointing to a decrease in neuronal support rather than full reactive gliosis. Identifying which link between maintenance and function becomes limiting in each cell type may point to therapeutic targets in HD and in other neurodegenerative diseases with protein aggregates.

## METHODS

### Animals and ethics

Experiments used R6/1 mice and age-matched wild-type littermates, all carrying the same Cre-dependent 3xMetRS* labelling alleles [11]. The labelling line was generated as described previously [9, 10]. Briefly, a STOPFLOX-3x(MetRS*-2A)-GFP cassette under the CAG actin-derived promoter was knocked into the mouse ROSA26 locus by recombination-mediated cassette exchange in embryonic stem cells, custom made for us by Taconic GmbH (available on request from BA-C, JAX #040661, C57BL/6 background). Cortical and hippocampal excitatory neurons were labelled by crossing homozygous floxed-Stop 3xMetRS* animals to homozygous mice of a Cre driver line specific for excitatory neurons (Nex-Cre, kindly provided by K. A. Nave) [88]. Cerebellar inhibitory neurons were labelled with PV-Cre mice (JAX #008069) and glia with Aldh1l1-Cre/ERT2 mice (JAX #031008) [22, 89]. In this study, glia refers to the cells labelled by this line. Cre expression in glia was induced with tamoxifen, dissolved in corn oil at 20 mg/ml and injected intraperitoneally at 100 µl per mouse (about 75 mg/kg) once a day for 5 consecutive days, at 4 months of age to avoid toxicity in older R6/1 mice [22]. Colonies were maintained as double homozygotes for 3xMetRS* and Cre to reduce animal numbers, and were checked regularly for germline activation and off-target expression by FUNCAT and GFP staining [10]. Both male and female mice were used. Group sizes were 4 to 6 animals per studied condition. The 3xMetRS* and wild-type alleles were genotyped as in [10].

### Azidonorleucine labelling and administration

ANL was administered as described previously [10]. Briefly, azidonorleucine (ANL, Iris Biotech, HAA1625) was prepared at 300 mM, pH 7.5, and injected intraperitoneally at 10 µl per g of body weight, once a day for 15 days. Animals were sacrificed 6 h after the last injection. Age-matched animals that received no ANL were processed in parallel as no-ANL controls [10]. At the end of the labelling period mice were euthanised with CO2 in accordance with local law. Brains were removed, dissected into cortex, hippocampus, striatum and cerebellum, and frozen on dry ice.

### Tissue lysis, BONCAT and click enrichment of labelled proteins

BONCAT in tissue was performed as described previously [10]. Briefly, dissected tissue was homogenised and lysed in PBS supplemented with 1% (w/v) Triton X-100 and 1% (w/v) SDS, with protease inhibitor (1:750 dilution of protease inhibitor cocktail 3 without EDTA, Calbiochem) and benzonase (Sigma, 1:1,000), for 15 min at 75 °C. Lysates were cleared by centrifugation and stored at −80 °C. Cleared extracts were alkylated twice with 20 mM iodoacetamide for 3 h at room temperature in the dark, then cleaned by two passages over PD-SpinTrap G-25 columns (GE Healthcare). Click chemistry was carried out in 120 µl PBS (pH 7.8) supplemented with 0.08% (w/v) Triton X-100 and 0.2% (w/v) SDS, using 90 µg protein, 300 µM triazole ligand (Sigma, 678937), 50 µM biotin-alkyne tag (Thermo Fisher Scientific) and 83 µg/ml CuBr freshly diluted from a 10 mg/ml stock in dimethyl sulfoxide. Reactions were incubated overnight at 4 °C in the dark. Excess biotin-alkyne was removed by trichloroacetic acid precipitation. Tagged proteins were affinity-purified overnight at 4 °C on high-capacity NeutrAvidin agarose beads (Thermo Fisher Scientific) in PBS containing 1% Triton X-100, 0.15% SDS and protease inhibitor (binding buffer). Beads were washed extensively at room temperature, first in binding buffer, then in 0.4% SDS and finally in PBS and in 50 mM ammonium bicarbonate, both with protease inhibitor. Bound proteins were eluted in two consecutive steps with 5% β-mercaptoethanol and 0.03% SDS with protease inhibitor for 30 min at room temperature.

### Peptide preparation and DIA mass spectrometry

Eluted proteins were precipitated and desalted using the SP3 method, as described previously [90]. Briefly, 30–35 µl of protein eluate containing β-mercaptoethanol (βME) were processed. Samples were incubated for 30 min at 37 °C to allow reduction of disulfide bonds, followed by alkylation with iodoacetamide (IAA) for 30 min at 37 °C. IAA concentrations were adjusted to the initial βME concentration. With 1% βME, estimated as 143 mM before dilution, IAA was set to exceed it, at a final 174 mM. For glia samples from cortex and striatum, where the βME concentration was 5%, IAA was adjusted to a final 300 mM. Samples were subsequently diluted with water to approximately 1 ml and proteins were precipitated onto SP3 magnetic carboxylated beads (1:1 mixture of hydrophobic and hydrophilic SpeedBeads) at a final bead concentration of approximately 1 mg/ml, by addition of 100% ethanol at a 1:1 (vol:vol) sample:ethanol ratio. Following 15 min incubation with agitation, protein-bound beads were immobilised on a magnetic rack and washed three times with 80% ethanol. Residual ethanol was removed, and beads were resuspended in 500 µl of 50 mM ammonium bicarbonate. Proteins were digested on-bead with 0.8 µg sequencing-grade trypsin (1:50 enzyme-to-substrate ratio) overnight at 37 °C with agitation (approximately 17.5–18.5 h). Following digestion, beads were removed by magnetic separation, and the resulting tryptic peptides were evaporated to dryness in a vacuum concentrator. Dried peptides were reconstituted in 15 µl of 3% acetonitrile (ACN)/0.2% formic acid (FA), sonicated for 3 min, and transferred to autosampler vials for LC-MS/MS analysis. About 1 µg of total peptides was analysed on a Waters M-Class UPLC using an IonOpticks Aurora Ultimate column (1.6 µm, 75 µm × 25 cm) coupled to a benchtop Thermo Fisher Scientific Orbitrap Q Exactive HF mass spectrometer. Peptides were separated at a flow rate of 400 nl/min with a 150 min gradient, including sample loading and column equilibration times, using solvents A (0.1% formic acid in water) and B (0.1% formic acid in acetonitrile). The gradient of solvent B started at 2% B for 1 min, increased linearly to 10% B over 39 min, to 22% B over 70 min, to 30% B over 7 min, to 60% B over 7 min and to 90% B over 1 min, where it was held for 5 min, then decreased linearly to 50% B over 1 min, was held for 9 min, decreased linearly to 2% B over 1 min and was kept at 2% B for re-equilibration for the rest of the acquisition period. Data were acquired in data-independent mode using Xcalibur software (4.5.474.0). MS1 spectra were measured with a resolution of 120,000, an AGC target of 3e6 and a scan range from 350 to 1600 m/z. MS2 spectra were measured in 35 segment windows per MS1, each with an isolation window width of 36 m/z (0.5 m/z overlap with the neighbouring window), a resolution of 30,000, an AGC target of 3e6 and a stepped collision energy of 22.5, 25 and 27.5.

### Database search and protein quantification

DIA raw files were processed as described previously [10]. Briefly, files were searched using DIA-NN (versions 1.9.1 for the glia dataset and 2.0.2 for all others) against a previously generated predicted spectral library based on the UniProtKB mouse reference proteome (UP000000589), with deep-learning-based spectrum and retention-time prediction. The spectral library was generated from an in silico tryptic digest (Trypsin/P), allowing up to two missed cleavages. Peptide length was set to 7–35 amino acids and precursor charge states to 1–5. N-terminal methionine excision, methionine oxidation and protein N-terminal acetylation were enabled, with the latter two specified as variable modifications (maximum of one variable modification per peptide) and cysteine carbamidomethylation as a fixed modification. MS1 and MS2 mass accuracies and scan windows were automatically optimised per search. Peptide and protein identifications were filtered at a 1% false discovery rate (FDR). Peptidoform scoring and retention-time profiling were enabled. Protein inference was performed using proteotypic peptides, and quantification and cross-run normalisation were performed using DIA-NN default settings (retention-time-based normalisation). DIA-NN reports (Parquet format) and the corresponding FASTA database were subsequently imported into R and processed using the MS-DAP package (version 1.2.3) for protein-level inference and label-free quantification, requiring a minimum of one peptide per protein.

#### Cell-type-specific enrichment and differential abundance

Cell-type-specific enrichment was assessed by comparing ANL-labelled samples with no-ANL controls, as described previously [10]. Two complementary criteria were used. The exclusivity criterion kept proteins quantified in ≤1 of the controls (≤2 of 6 in cortex). It also required quantification in at least 2 of 4 labelled samples of a genotype for glia, 3 of 4 for hippocampal and cerebellar neurons and 4 of 6 for cortical neurons (3 of 5 when the group had five). The enrichment criterion was based on relative enrichment over control. Differential abundance was then modelled with a peptide-centric linear mixed-effect model as implemented in the msqRob algorithm within the MS-DAP pipeline, and p values were corrected for multiple testing by the Benjamini–Hochberg method [91].

We compared R6/1 mice with age-matched wild-type littermates, all carrying the Cre-dependent 3xMetRS* labelling system. Cortical and hippocampal excitatory neurons were sampled at 1.5, 3 and 7 months, cerebellar inhibitory neurons at 3 and 7 months, and glia from cortex, striatum and cerebellum at 7 months. Because labelled proteins were enriched from the ANL-labelled pool, each sample contains the proteins made during the 15-day labelling window that persisted until harvest. Hippocampal glia and no-ANL controls were excluded from the main biological comparisons.

The comparison between R6/1 mice and age-matched wild-type littermates was carried out by the msqRob algorithm within the MS-DAP pipeline, and no built-in normalisation was applied at this stage. Instead, to correct for systematic between-sample loading differences while preserving relative differences across conditions, a customised median-centring normalisation method was applied at peptide intensity level. For each type of neurons in each tissue, a global reference value was computed as the median peptide intensity across samples from all time points. For each individual sample, the per-sample median intensity was calculated, and each peptide intensity was normalised as normalised intensity = (raw intensity / sample median) × global median.

Protein-group identities, log2 abundance ratios (R6/1 over WT), raw p values and Benjamini–Hochberg (BH) q values computed within each contrast carry the output of the differential-abundance model described above. Positive values indicate higher abundance in R6/1. Protein groups with q < 0.1 were called quantitatively regulated, with no fold-change cut-off. Quantitative calls also required quantification in at least 2 of 4 samples of each genotype (3 in cortex). Genotype-enriched qualitative detection calls required quantification in at least 3 of 4 samples of one genotype (4 of 6, or 3 of 5, in cortex) and in at most 1 sample of the other. They carry no fold change or p value and were added to the regulated lists. All reanalyses used these protein-group statistics as they were. The sample-level differential-abundance model was not refitted.

Total MS signal before normalisation was compared between genotypes within each preparation using two-sided Mann–Whitney U tests. The signal was the DIA-NN Total.Quantity of each run, from the DIA-NN run summary. The quality-control figures keep the run exclusions of the primary analysis and report unadjusted p values. Sample similarity used Spearman correlations, and the per-sample median log2 intensities in Supplementary File 1 were compared with two-sided Welch t-tests. A non-significant difference was not treated as evidence of equivalence.

### Data analysis

We used PANTHER for functions enriched in the regulated-protein lists, split by direction and by the age or regional overlap that defined each list, and IPA for canonical pathways across all quantitatively regulated proteins in each comparison. The first describes what is specifically regulated in a particular overlap, the second describes the whole cellular response. PANTHER was run with nine annotation sets, the Gene Ontology complete and GO-Slim sets for biological process, molecular function and cellular component, PANTHER Pathways, PANTHER Protein Class and Reactome. Over-representation was tested with Fisher’s exact test and BH correction, and terms with FDR < 0.05 were called over-represented. All over-represented terms are listed in Supplementary Table 3, and the figures show selected terms from any of these sets. In the Venn diagram of all neurons against all glia (Supplementary Fig. 9), a protein that changed in opposite directions in two populations of the same cell type, for example up in cortical and down in hippocampal neurons, has no single direction for that cell type. It was counted in the total of that cell type but placed in no section of the diagram, so the sections add up to slightly less than the total. Supplementary Table 3 lists these proteins in three separate columns. Enrichment p values describe the representation of an annotation, whereas IPA activation z-scores are predictions based on the mapped members and their direction. Pathway labels that shared proteins were not counted as independent biological responses. The preparations are enriched and not pure, so proteins specific to one population were occasionally also found in the preparations of another. In the glial preparations we flagged the proteins most likely made by neurons. For each protein, we compared its highest level among glial cell types with its highest level among neuronal types in three datasets: the proteomes of cultured mouse brain cells, single-nucleus RNA of human brain (Human Protein Atlas) and single-cell RNA of mouse cortex and hippocampus [92–94]. A protein was flagged when at least two datasets measured it and glia had less than half of the neuronal level in all of them. A protein annotated to the presynapse, the postsynapse, the axon or synaptic signalling (Gene Ontology, MSigDB 2026.1) was flagged unless glia had at least half of the neuronal level in at least two datasets. This rule was not applied to proteins at least 2-fold more abundant in isolated astrocytes, microglia or oligodendrocytes than in isolated neurons [92]. This flagged 216 of the 1,931 proteins of the glial lists. This classification was quite strict: for 102 of the 216 proteins, glia reached at least a quarter of the neuronal level in at least one dataset. Only 114 of the 1,931 proteins (5.9%) were clearly enriched in neurons (glia had less than a quarter of the neuronal level in every dataset that measured them). A glial PANTHER term or IPA pathway was described only when more than 70% of its non-proteasome members were not flagged, but the full PANTHER results can be found in the tables. Flagged proteins were left out only of the glial cameraPR tests. With or without them, cameraPR gave the same calls in all 107 tests except one, the cytosolic ribosome in cerebellar glia (q = 0.044 with them and 0.084 without).

Pathway analysis was performed with Ingenuity Pathway Analysis (IPA, QIAGEN, version 1.24.1) on the proteins with q < 0.1, using the nervous system reference as biological filter.

### Competitive protein-set tests

Competitive set tests used cameraPR in limma 3.62.2 (R 4.4.1) [27, 28]. In each comparison, the test asked whether the protein groups of a set changed more than all other protein groups of that comparison. The interferon-responsive set held the 66 genes enriched in interferon-responsive astrocytes (cluster 8, adjusted p < 0.05 and positive average log fold change) [29]. Its genes were matched to protein groups by gene symbol or UniProt gene name, not by accession. For each contrast we kept protein groups with a finite log2 ratio and raw p value, removing a row only when p equalled 1 and the absolute log2 ratio was below 1 × 10−9. Proteins were ranked by a signed normal score derived from the two-sided raw p value and the sign of the log2 ratio, with p floored at 1 × 10−15. Tests used use.ranks = FALSE and an assumed inter-protein correlation of 0.01, and a set was tested only when at least five of its protein groups qualified. A protein group was assigned to a set once if any of its UniProt accessions belonged to the set, and all other qualifying protein groups in that contrast formed the background. Qualitative calls were not included for this test. BH correction was applied across the tested contrasts within each set, and sets with q < 0.05 were called significant.

Protein groups were matched across comparisons by exact UniProt group. Checks of response breadth on shared proteins used only protein groups with a finite log2 ratio and p value in every comparison involved. q values were examined at q < 0.1, and p values were not pooled across ages. Missing values were left missing, and near-zero log2 ratios were not read as recovery.

### Weighted gene co-expression network analysis

Weighted gene co-expression network analysis (WGCNA) was performed by the Genomics and Bioinformatics Platform of IdISBa with the WGCNA R package 1.74 in R 4.5.2 [20, 21]. One network was built for cortical and one for hippocampal excitatory neurons, each from all R6/1 and wild-type mice at 1.5, 3 and 7 months, using the proteins present at all three ages. Log2 intensities were median-centred per mouse. Signed hybrid networks were built using biweight midcorrelations (maxPOutliers = 0.05), with soft-thresholding powers of 6 (cortex) and 8 (hippocampus), and they were converted into a topological overlap matrix (TOM). Proteins were clustered by average linkage on 1-TOM and modules of proteins were identified using a dynamic tree-cutting algorithm (deepSplit = 2, minClusterSize = 30). Modules whose eigengenes had a dissimilarity below 0.15 were merged. Each module was summarised by its eigengene, the first principal component of the levels of its proteins. The eigengene of each module was correlated with genotype (Pearson correlation, Student’s t-test), with all ages pooled and within each age, and the p values were adjusted with the Benjamini-Hochberg method across the modules of each region. Modules with q < 0.05 with all ages pooled were called genotype modules. The GO biological process terms and KEGG pathways of each genotype module were tested for over-representation with clusterProfiler 4.18.4 and org.Mm.eg.db, using a hypergeometric test with Benjamini-Hochberg correction (adjusted p < 0.05), with the proteins of both regions as background [95, 96]. The genotype modules of the two regions were compared on the proteins present in both networks, by a hypergeometric test of their overlap, with BH correction across the 30 module pairs (FDR < 0.05), and with the modulePreservation function of WGCNA, using the same network settings and 100 permutations [97]. Modules with Zsummary > 10 were considered strongly preserved, those with 2 to 10 moderately preserved, and those with < 2 not preserved.

### Protein length of the network modules

Module C4, which had no enriched GO or KEGG term, was first noticed in an exploratory screen of protein properties made by BA-C with an AI assistant (see Declarations). Its length was then tested against our earlier proteomes of whole hippocampus (4,947 protein groups) and of labelled excitatory neurons (2,384 protein groups), using the sequence lengths given in those tables [9]. Module proteins were matched to the references by UniProt accession, and 32 of the 38 C4 proteins were found in the whole hippocampus proteome. For each of the 11 modules, lengths were compared with the reference by a two-sided Mann-Whitney test. The number of proteins longer than the 75th percentile of the reference (780 amino acids) was compared by an upper-tail hypergeometric test, with Benjamini-Hochberg correction across the modules (SciPy 1.13.1) [98]. The same tests were repeated against the labelled proteome (75th percentile 819.5 amino acids). The C4 eigengene in hippocampus (Supplementary Fig. 6c) was computed with the same code as the eigengenes of Fig. 5.

## Supporting information

MS-DAP Files

ProteinHeatMAPS

Graphical_Abstract

## Acknowledgements

We are especially grateful to M. Rahmatian for her great help and excellent technical assistance throughout this work. We thank C. Lozano, D. F. Rincón Cubillos, and I. Monge Rico for excellent technical assistance, and the animal facility of the Faculty of Medicine of the Complutense University of Madrid (CAI Animalario) for their excellent support. We thank the master’s student H. Arevalo Blanco, the bachelor students M. Sánchez-Escalonilla Martín-Caro, G. R. de Vicente Martinez and S. Vallejo Garlito for their help with some of the experiments.

## Abbreviations

AMPA: α-amino-3-hydroxy-5-methyl-4-isoxazolepropionic acid.
ANL: azidonorleucine.
ARC: activity-regulated cytoskeleton-associated protein.
BH: Benjamini-Hochberg.
BONCAT: bio-orthogonal noncanonical amino acid tagging.
CAG: cytosine-adenine-guanine trinucleotide repeat.
CCT/TRiC: chaperonin containing TCP-1 / TCP-1 ring complex.
DIA-MS: data-independent acquisition mass spectrometry.
DIA-NN: data-independent acquisition neural network.
DLG4: discs large MAGUK scaffold protein 4 (PSD-95).
DNAJB1: DnaJ heat shock protein family (Hsp40) member B1.
ER: endoplasmic reticulum
FDR: false discovery rate.
GLT-1: glutamate transporter 1 (SLC1A2).
GO: Gene Ontology.
HD: Huntington’s disease.
HSP70: heat-shock protein 70.
HTT: huntingtin.
IPA: Ingenuity Pathway Analysis.
KEGG: Kyoto Encyclopedia of Genes and Genomes.
Kir4.1: inwardly rectifying potassium channel 4.1 (KCNJ10).
LC-MS: liquid chromatography-mass spectrometry.
MetRS*: modified methionyl-tRNA synthetase.
MS: mass spectrometry.
NPTXR: neuronal pentraxin receptor.
PANTHER: Protein Analysis Through Evolutionary Relationships.
SNARE: soluble N-ethylmaleimide-sensitive factor attachment protein receptor.
TCA: tricarboxylic acid.
TCERG1: transcription elongation regulator 1.
UPS: ubiquitin-proteasome system.
WGCNA: weighted gene co-expression network analysis.
WT: wild type.

## Authors’ contributions

RA-P and ED-M performed experiments and analysed the data. SL analysed the data. PR-R analysed the data and edited the manuscript. JLS performed experiments. HA-G analysed the data and edited the manuscript. CO-S performed experiments. KD and MD-H edited the manuscript. EFF analysed the data and edited the manuscript. MJ designed experiments and analysed the data and edited the manuscript.

MG-O and JM performed the WGCNA analysis. BA-C designed and performed experiments, analysed the data and wrote the manuscript. All authors read and approved the final manuscript.

## Availability of data and materials

The mass spectrometry proteomics data have been deposited to the ProteomeXchange Consortium via the PRIDE partner repository. The dataset identifier will be provided in the published version of the article, and until then the data can be requested from the corresponding authors. Protein-group identities, log2 abundance ratios (R6/1 over wild type), raw p values and Benjamini-Hochberg q values for every contrast are given in Supplementary Tables 1 and 4. All code used for analysis is available on request from the corresponding authors.

## Ethics approval and consent to participate

All experiments involving animals were performed with permission from the local government offices in Spain (Committee of Animal Experiments at UCM and Environmental Counselling of the Comunidad de Madrid, protocol number PROEX 005.0/21). These experiments comply with Spanish regulations and follow European Union (EU) guidelines for animal welfare. Mice were housed in accordance with EU Directive 2010/63/EU Annex III and euthanised using methods in accordance with EU Directive 2010/63/EU Annex IV. This study did not involve human participants, human data or human tissue.

## Consent for publication

Not applicable.

## Use of artificial intelligence

BA-C used a large language model assistant (Claude, Anthropic) under her direction and checked every output. It was used for language editing and for checking grammar, style, and numbering. It wrote scripts that ran published tools on the data, such as cameraPR in limma, and assembled the figure files from these outputs under her revision. An exploratory, unsupervised screen made with the assistant suggested that the proteins of module C4 were long. They were then tested as described in the Methods (Supplementary Fig. 6). All scripts are available on request.

