## SupplementaryFigures for "Excitatory neurons downregulate synaptic proteins before motor onset and glia reduce neuronal support in Huntington’s disease mice"

#### Supplementary Figure 1

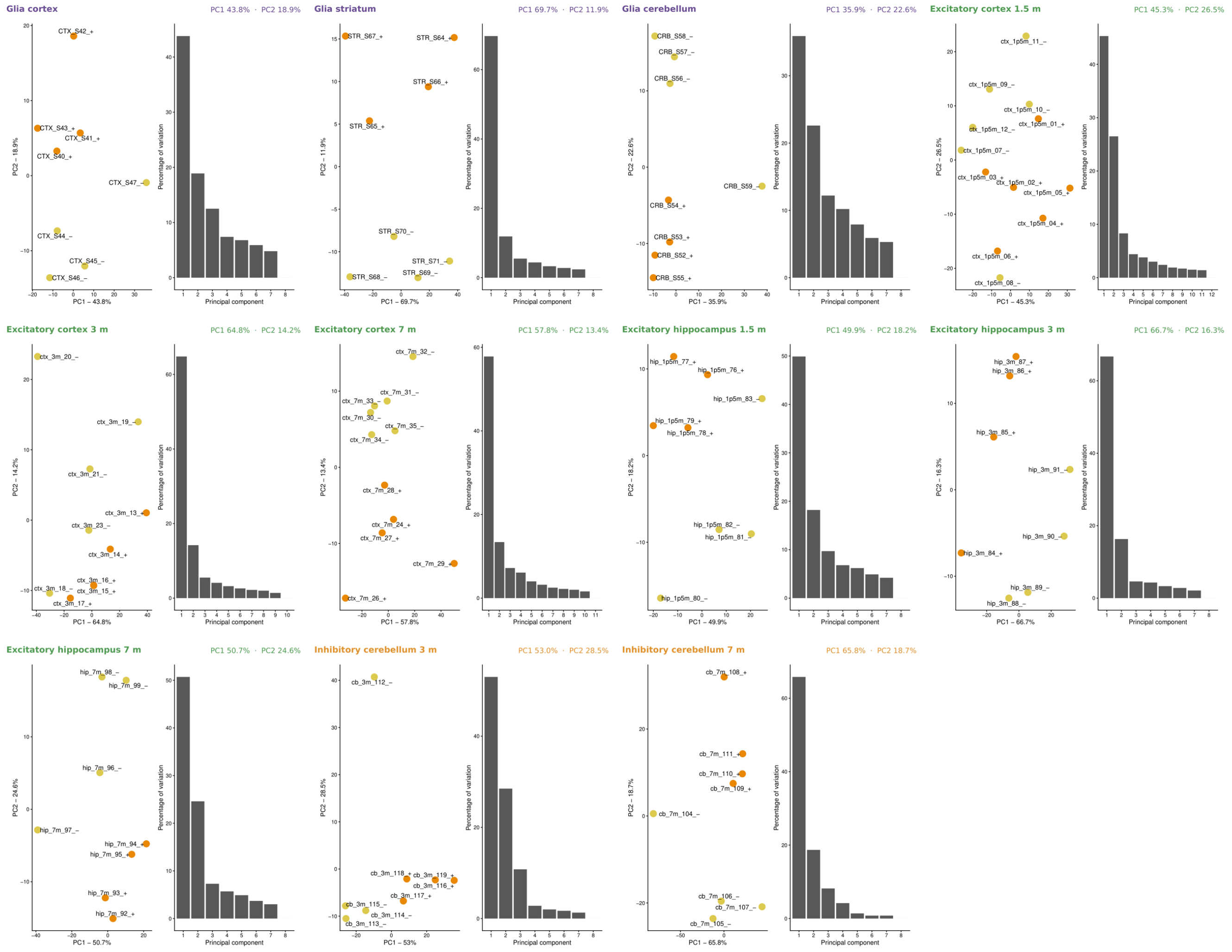


**Supplementary Fig. 1 Principal component analysis of each population.** Each pair of plots shows one population, glia of cortex, striatum and cerebellum at 7 months, excitatory neurons of cortex and hippocampus at 1.5, 3 and 7 months, and inhibitory neurons of cerebellum at 3 and 7 months. The left plot shows the samples on the first two principal components, R6/1 in orange and WT in yellow. Sample labels end in + for R6/1 and – for WT. The right plot shows the percentage of variation carried by each component. Component 1 carried 35.9 to 69.7% of the variation and component 2 a further 11.9 to 28.5%. n = 4 mice per genotype, except for cortical excitatory neurons (6 R6/1 and 6 WT at 1.5 months, 5 and 5 at 3 months, 5 R6/1 and 6 WT at 7 months).

#### Supplementary Figure 2

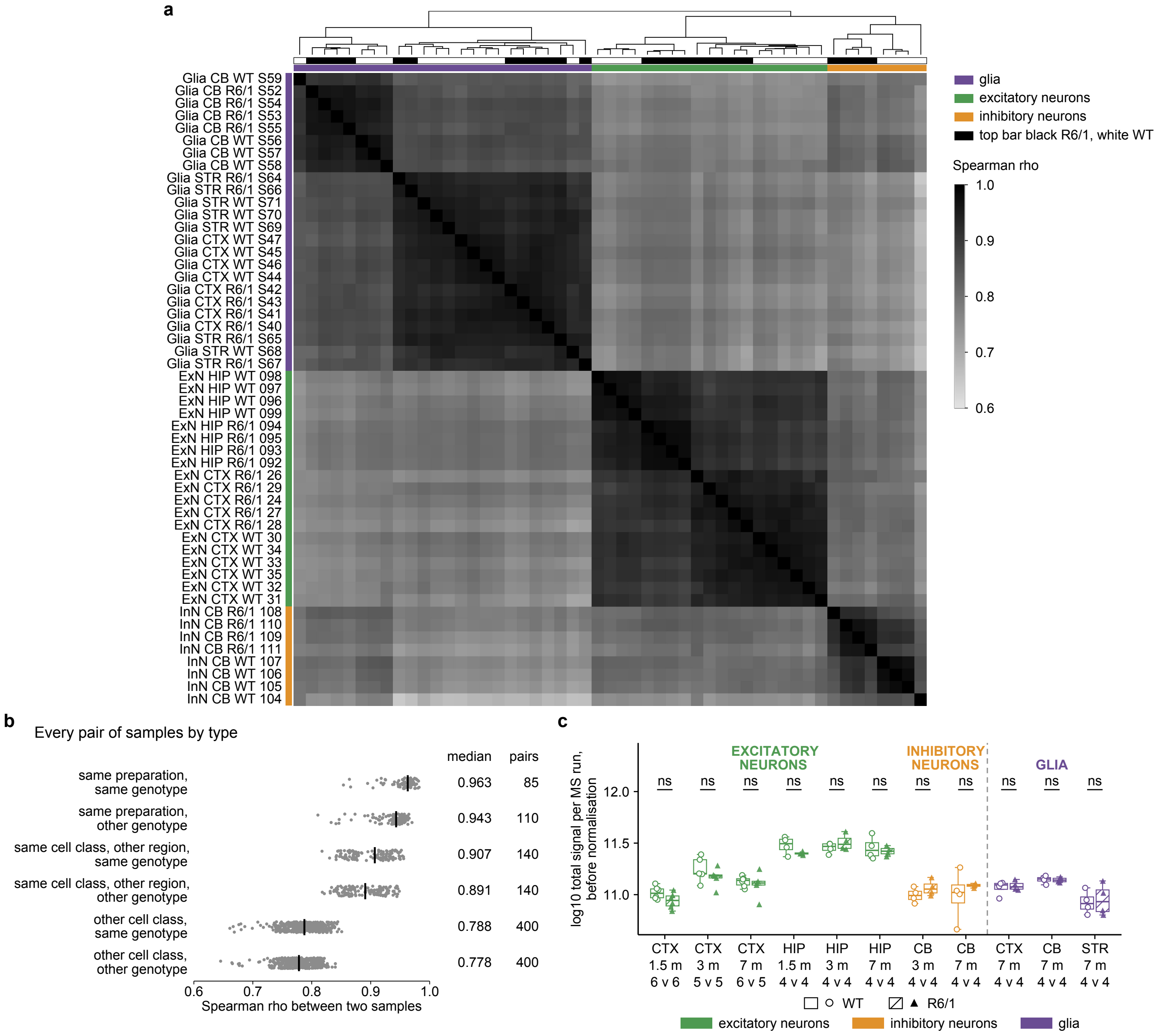


**Supplementary Fig. 2 Sample similarity and total precursor intensity of R6/1 and WT samples**. **(a)** Spearman correlation (rho) between every pair of the 51 samples at 7 months (26 WT and 25 R6/1). Each correlation used the proteins quantified in both samples of the pair, ranked by intensity within each sample. Ranks were compared instead of intensities because the populations were measured in four acquisitions and normalised separately. Samples are ordered by average-linkage clustering on 1 − rho. In the upper bars, purple marks glia, green excitatory neurons and orange inhibitory neurons, and black marks R6/1 and white WT. **(b)** Every pair of samples shown in panel (a) grouped by condition, with the median rho (black line) and the number of pairs. **(c)** Total precursor intensity of each run before normalisation, from the DIA-NN run summary, for the 97 runs of the 11 populations (49 WT and 48 R6/1), as log10. Boxes give the median and quartiles. WT runs are plain boxes with round points and R6/1 runs hatched boxes with square points. The numbers under each pair of boxes are the WT and R6/1 run counts. Genotypes were compared within each population (two-sided Mann-Whitney U test), and no comparison was significant (ns, $p \geq 0.05$). All values are in Supplementary Table 2. ExN, excitatory neurons. InN, inhibitory neurons. CTX, cortex. HIP, hippocampus. CB, cerebellum. STR, striatum. m, months. n = 4 mice per genotype, except for cortical excitatory neurons (6 R6/1 and 6 WT at 1.5 months, 5 and 5 at 3 months, 5 R6/1 and 6 WT at 7 months).

#### Supplementary Figure 3

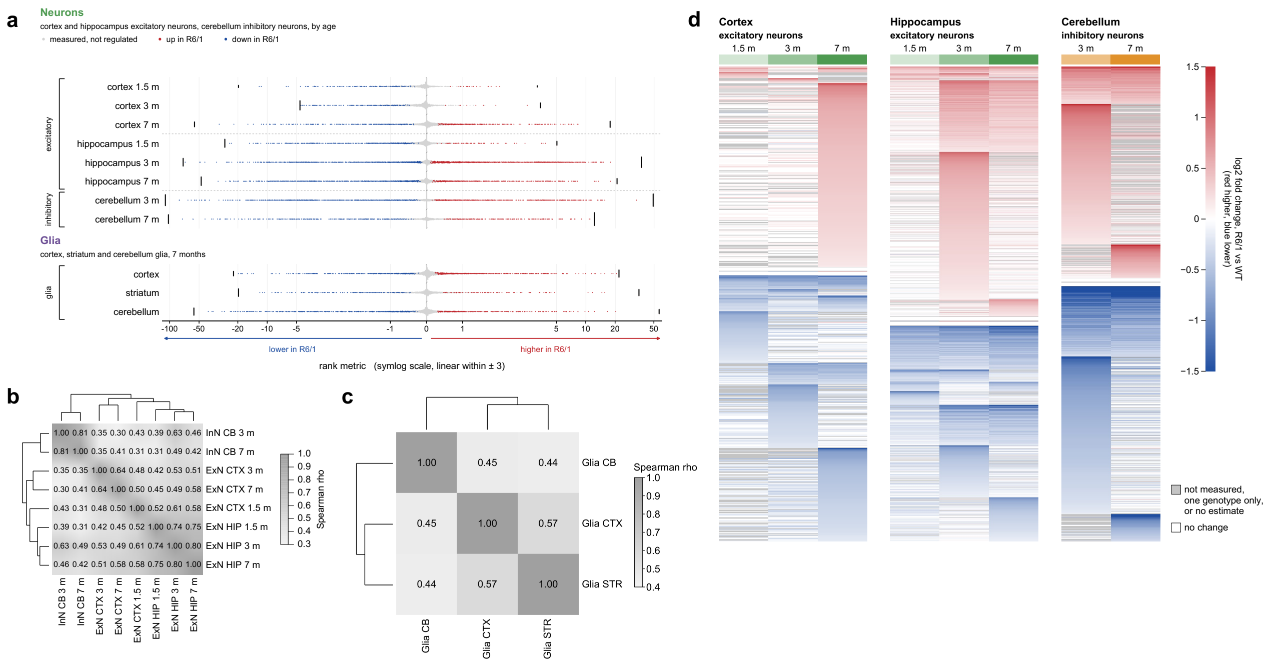


**Supplementary Fig. 3 Protein changes in R6/1 mice by population, age and region**. **(a)** Every measured protein with a p value in each comparison of R6/1 with WT littermates of the same age, placed by its rank metric, −log10 p multiplied by the log2 fold change (msqRob). The x axis is on a symmetric log scale, linear between −3 and 3. Red and blue, upregulated and downregulated in R6/1 (Benjamini-Hochberg $q < 0.10$, no fold-change cut-off). Grey, not regulated. The grey column at zero holds proteins with a fold change of zero or no estimate. Black bars at the row ends are the proteins detected in one genotype (qualitative counts). Having no fold change, they sit at the highest (R6/1 only) or lowest (WT only) rank metric of their comparison, and bar height represent their number. Red dots, blue dots and black bars are the proteins counted in Fig. 2. **(b)** Spearman correlation ($\rho$) of R6/1 log2 fold changes between the neuronal comparisons, over all proteins quantified in both, regulated or not. Rows and columns follow average-linkage clustering on $1 - \rho$. **(c)** The same for glia at 7 months. **(d)** Log2 fold change (R6/1/WT) at each age, of the proteins with a gene symbol and $q < 0.10$ in cortical, hippocampal excitatory neurons and in cerebellar inhibitory neurons. Rows are grouped by direction (higher, changing direction, lower) and ordered within each group by the first age with $q < 0.10$. Colours stop at ±1.5. White, no change. Grey, no fold change at that age (not quantified, no estimate, or detected in one genotype only). The values of panels (b) to (d) are in Supplementary Table 2. ExN, excitatory neurons. InN, inhibitory neurons. CTX, cortex. HIP, hippocampus. CB, cerebellum. STR, striatum. m, months. n = 4 mice per genotype, except for cortical excitatory neurons (6 R6/1 and 6 WT at 1.5 months, 5 and 5 at 3 months, 5 R6/1 and 6 WT at 7 months).

**Supplementary Figure 4**

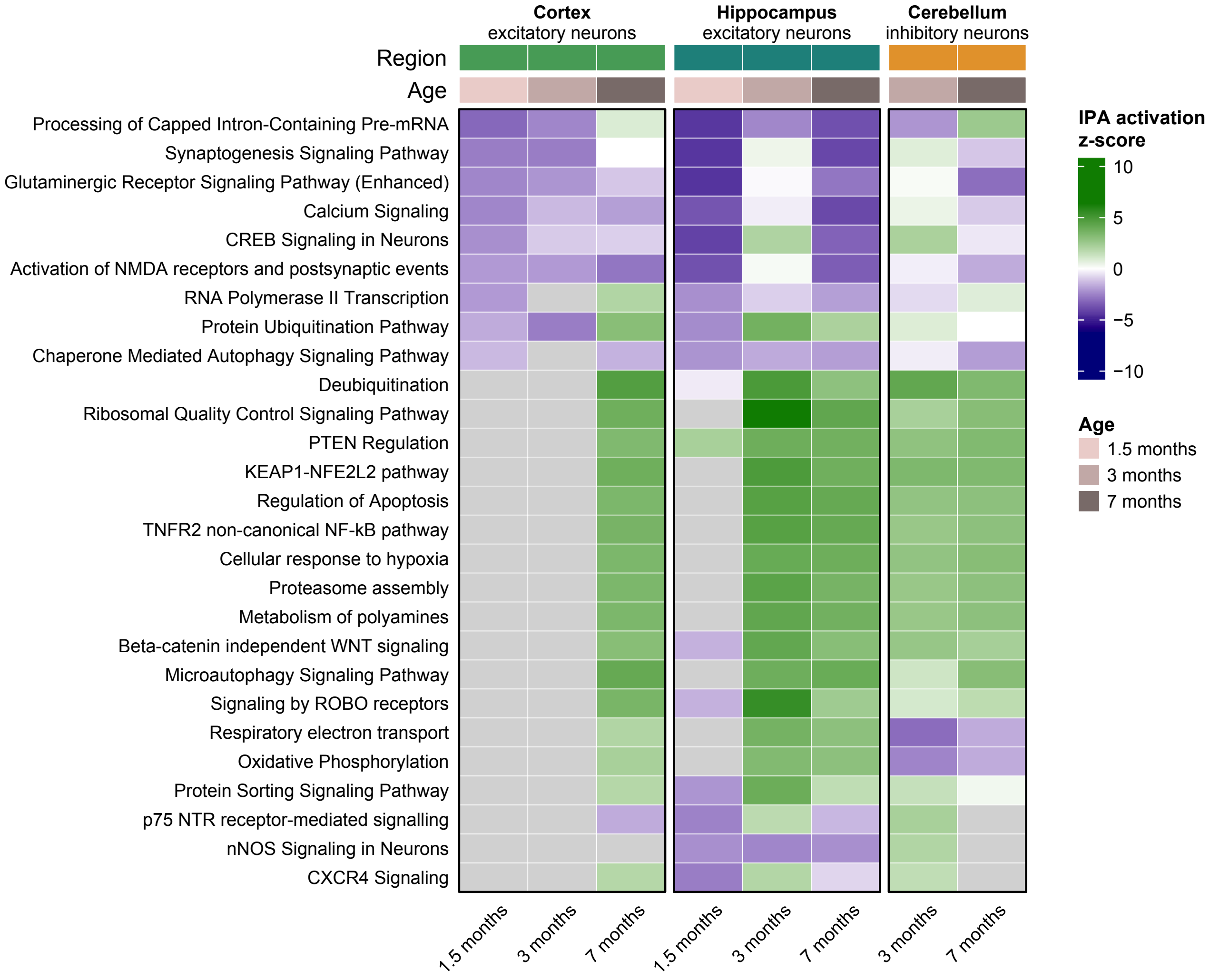


**Supplementary Fig. 4 IPA pathways in neurons of R6/1 mice across regions and ages**. Heat map of the IPA activation z-scores of 27 representative canonical pathways, R6/1 against WT, in excitatory neurons of cortex and hippocampus at 1.5, 3 and 7 months and in inhibitory neurons of cerebellum at 3 and 7 months. Each column is one comparison and each row one pathway, and the bars above the heat map give the region and the age. The z-scores come from an IPA comparison analysis of the eight core analyses shown in Fig. 3d, 4d and 6c, each made on the proteins regulated in that comparison ($q < 0.10$). Colour gives the activation z-score, as in the scale bar, green when positive and purple when negative. A positive z-score predicts that the pathway increases in R6/1 neurons, and a negative z-score that it decreases. The z-score is a prediction made from the direction of change of the regulated proteins in the pathway. Grey cells, no z-score computed. Cells are coloured whether or not the pathway was significantly enriched in that comparison, and the p value of every pathway is given in Supplementary Table 4. n = 4 mice per genotype, except for cortical excitatory neurons (6 R6/1 and 6 WT at 1.5 months, 5 and 5 at 3 months, 5 R6/1 and 6 WT at 7 months).

**Supplementary Figure 5**

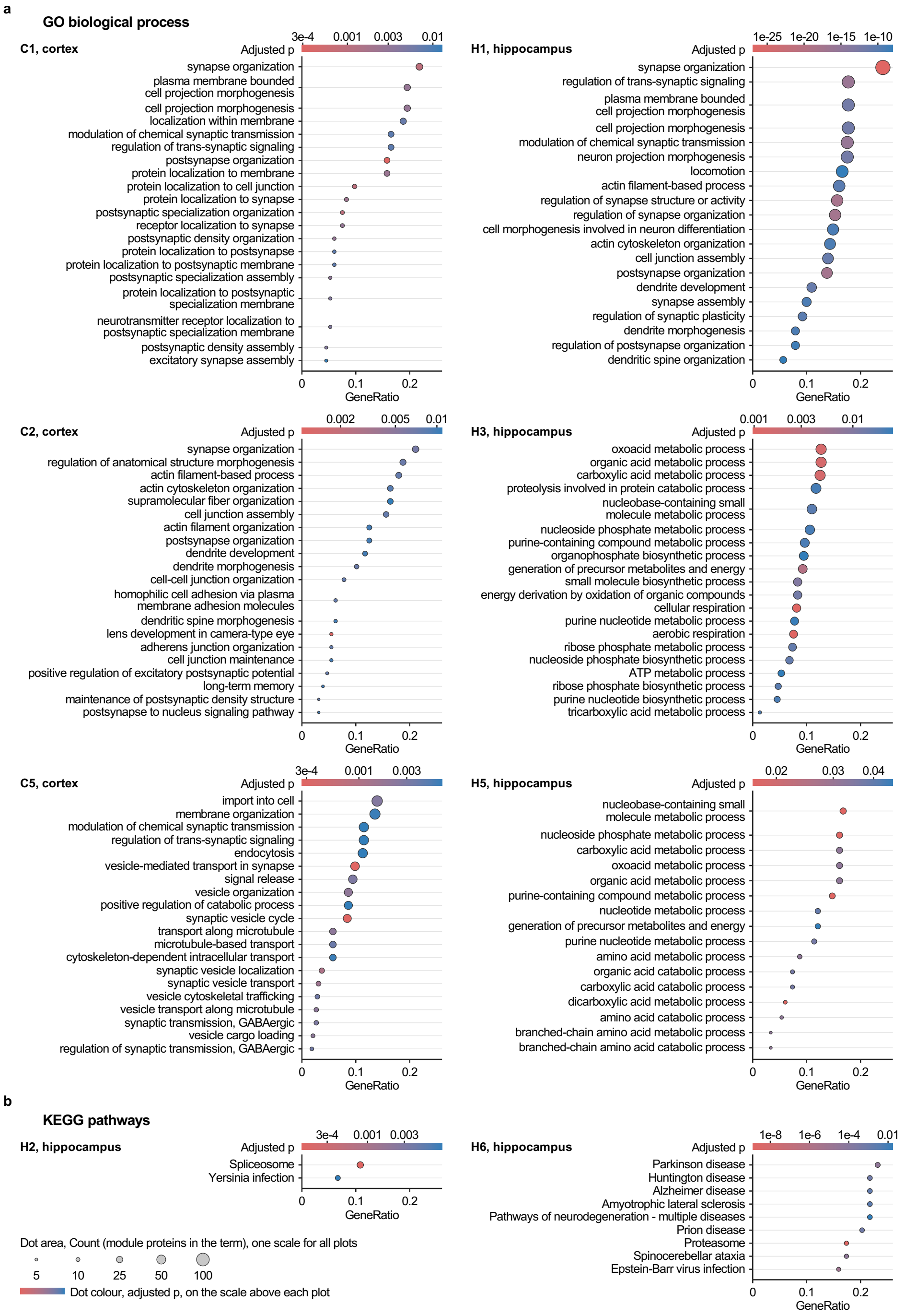


**Supplementary Fig. 5 GO terms and KEGG pathways over-represented in protein co-expression modules linked to R6/1 in cortical and hippocampal excitatory neurons found in the WGCNA analysis. (a)** GO biological process terms over-represented in modules C1, C2 and C5 of cortex and H1, H3 and H5 of hippocampus, the modules of Fig. 5 that returned GO terms (clusterProfiler, Benjamini-Hochberg adjusted $p < 0.05$ within each module). Each plot shows the 20 terms with the smallest p values (all 16 for H5). **(b)** All KEGG pathways over-represented in H2 (2 pathways) and H6 (9 pathways), tested in the same way. These are the two modules with a KEGG pathway in Fig. 5c. In every plot, rows are ordered by GeneRatio (x axis, largest at the top), the number of module proteins in the term divided by the module proteins with a GO biological process (a) or KEGG (b) annotation. Dot size (Count) gives the number of module proteins in the term or pathway and colour the adjusted p (p.adjust), red for the smallest values, as in the scale bar of each plot. All GO terms and KEGG pathways of all modules, with their proteins, are in Supplementary Table 5. n = 4 mice per genotype at each age, except in cortex (6 R6/1 and 6 WT at 1.5 months, 5 and 5 at 3 months, 5 R6/1 and 6 WT at 7 months).

#### Supplementary Figure 6

**a C4 is made of long proteins**

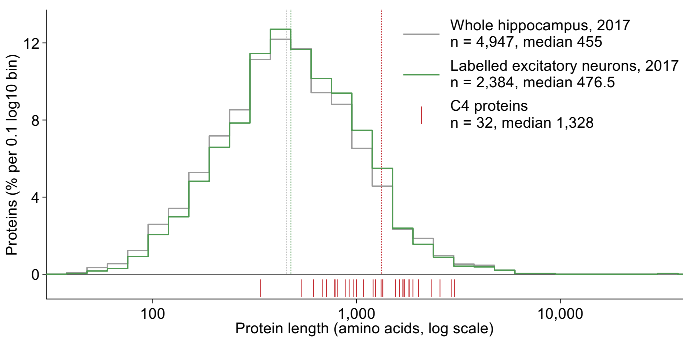


**b C4 proteins, length (amino acids)**

| | | | |
|---|---|---|---|
| FRY | 3,020 | SPAG9 | 1,320 |
| NBEA | 2,936 | SLC4A3 | 1,239 |
| TRIO | 2,569 | EHBP1 | 1,206 |
| CACNA1B | 2,325 | USP8 | 1,080 |
| CKAP5 | 2,011 | TAOK1 | 1,001 |
| UBR3 | 1,892 | KIF5B | 963 |
| AFDN | 1,827 | WDR47 | 920 |
| TSC2 | 1,808 | ANKLE2 | 886 |
| ITSN1 | 1,714 | EXOC6 | 804 |
| CDC42BPB | 1,713 | SMCR8 | 785 |
| CDC42BPA | 1,691 | VAC14 | 782 |
| PTPN23 | 1,690 | RUFY1 | 712 |
| KIF21A | 1,628 | PCDH10 | 683 |
| ARHGEF11 | 1,552 | LMNB2 | 615 |
| GPRASP1 | 1,347 | NRBP1 | 535 |
| MAPK8IP3 | 1,336 | GLRX3 | 337 |

**c C4 eigengene in WT and R6/1 at each age, hippocampus**

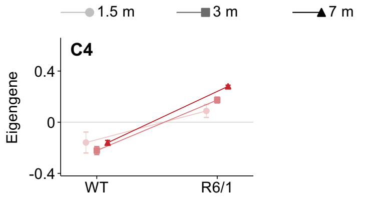


**Supplementary Fig. 6 Protein length of the cortical module C4 and its eigengene in hippocampus found in the WGCNA analysis.** C4 is the module of 38 proteins of cortical excitatory neurons that is higher in R6/1 (Fig. 5). **(a)** Distribution of protein length in two proteomes of our earlier study, the whole hippocampal proteome (grey, 4,947 protein groups) and the ANL-labelled proteome in excitatory neurons (green, 2,384 protein groups) [9]. The x axis is protein length in amino acids, on a log scale, and the y axis is the percentage of proteins of each proteome at each length. Red ticks under the histogram mark the 32 C4 proteins found in the whole hippocampal proteome. Dotted lines mark the medians, 455 (grey), 476.5 (green) and 1,328 amino acids (red). C4 proteins were longer than those of the whole hippocampal proteome (two-sided Mann-Whitney U test, Benjamini-Hochberg correction across the 11 modules linked to R6/1, $q = 2.0 \times 10^{-11}$), and 27 of the 32 were in its longest quarter (above 780 amino acids), where 8 were expected (hypergeometric test, upper tail, same correction, $q = 2.5 \times 10^{-11}$). **(b)** Length of the 32 C4 proteins found in the whole hippocampal proteome, longest first. The other 6 C4 proteins are not in that proteome. **(c)** Eigengene of C4 in hippocampal excitatory neurons of WT and R6/1 mice at 1.5, 3 and 7 months (mean and s.e.m.), from the 35 C4 proteins also present in the hippocampal network of Fig. 5. Red, as for the modules higher in R6/1 in Fig. 5c. Each line joins the WT and R6/1 means of one age, light with a circle for 1.5, medium with a square for 3 and dark with a triangle for 7 months. m, months. n = 4 R6/1 and 4 WT mice at each age.

#### Supplementary Figure 7

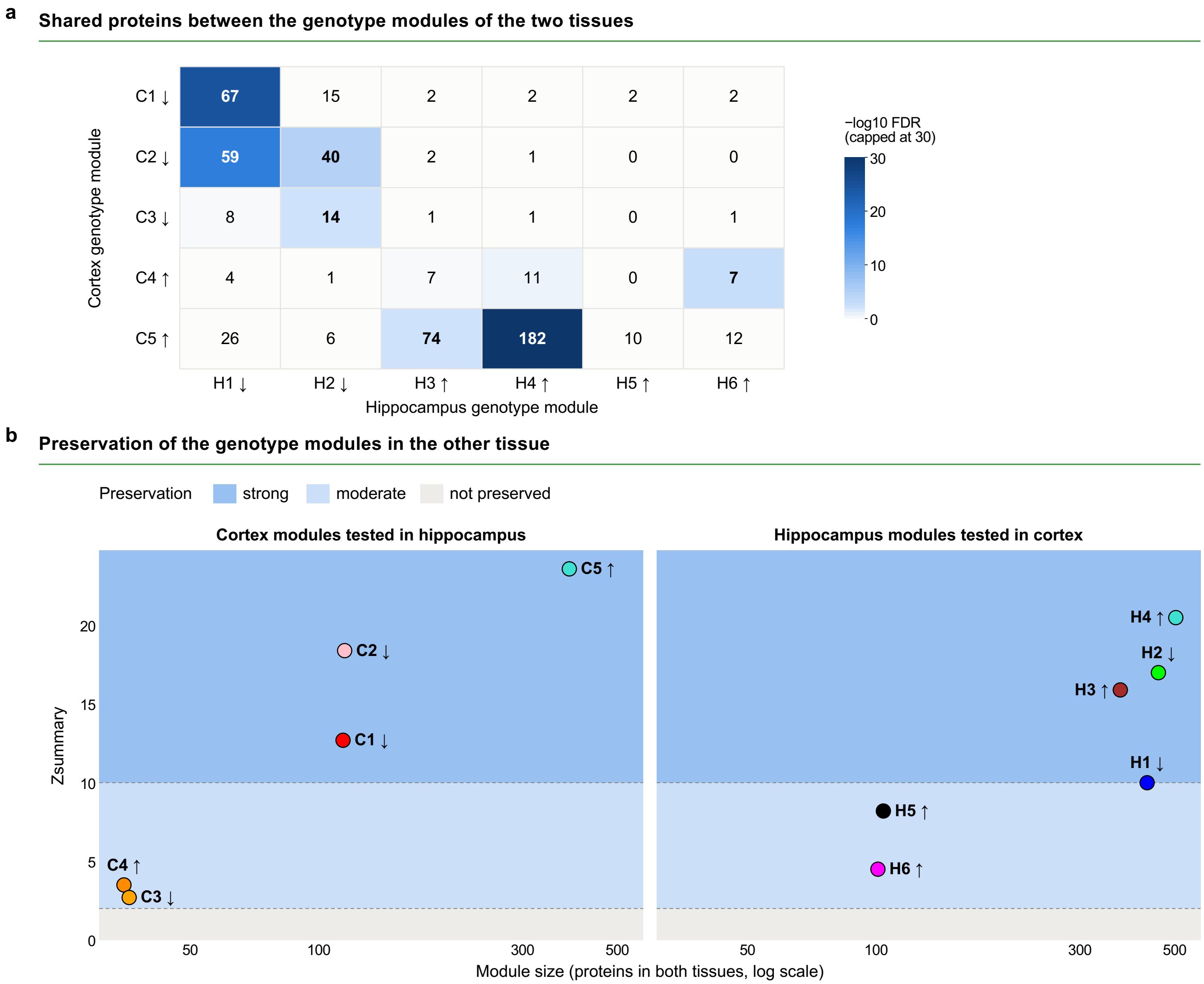


**Supplementary Fig. 7 Overlap and preservation of the modules linked to R6/1 between cortical and hippocampal excitatory neurons found in the WGCNA analysis.** The modules of Fig. 5 were compared on the 2,673 proteins present in both networks. Arrows mark modules lower (↓) or higher (↑) in R6/1. **(a)** Number of proteins shared by each cortical module (rows, C1 to C5) and each hippocampal module (columns, H1 to H6). Bold numbers mark the seven pairs that shared more proteins than expected by chance (one-sided hypergeometric test with Benjamini-Hochberg correction across the 30 pairs, FDR $< 0.05$). Cell colour gives −log10 FDR, capped at 30, as in the scale bar. **(b)** Preservation of each module in the network of the other region, cortical modules in the hippocampal network (left) and hippocampal modules in the cortical network (right). Each dot is one module, in its colour of Fig. 5a, placed by its number of proteins present in both networks (x axis, log scale) and its Zsummary preservation score (y axis, WGCNA modulePreservation). The shading marks the preservation classes, as in the key, strong (Zsummary above 10), moderate (2 to 10) and not preserved (below 2), with dashed lines at 2 and 10. H1 sits on the line at 10 and is classed moderate. All counts, FDR and Zsummary values are in Supplementary Table 5. FDR, false discovery rate. WGCNA, weighted gene co-expression network analysis. n = 4 mice per genotype at each age, except in cortex (6 R6/1 and 6 WT at 1.5 months, 5 and 5 at 3 months, 5 R6/1 and 6 WT at 7 months).

#### Supplementary Figure 8

a

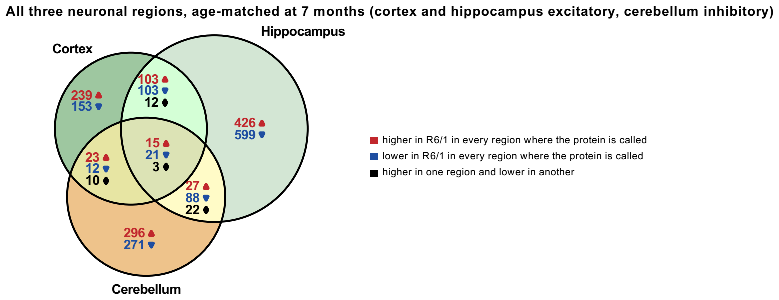


b

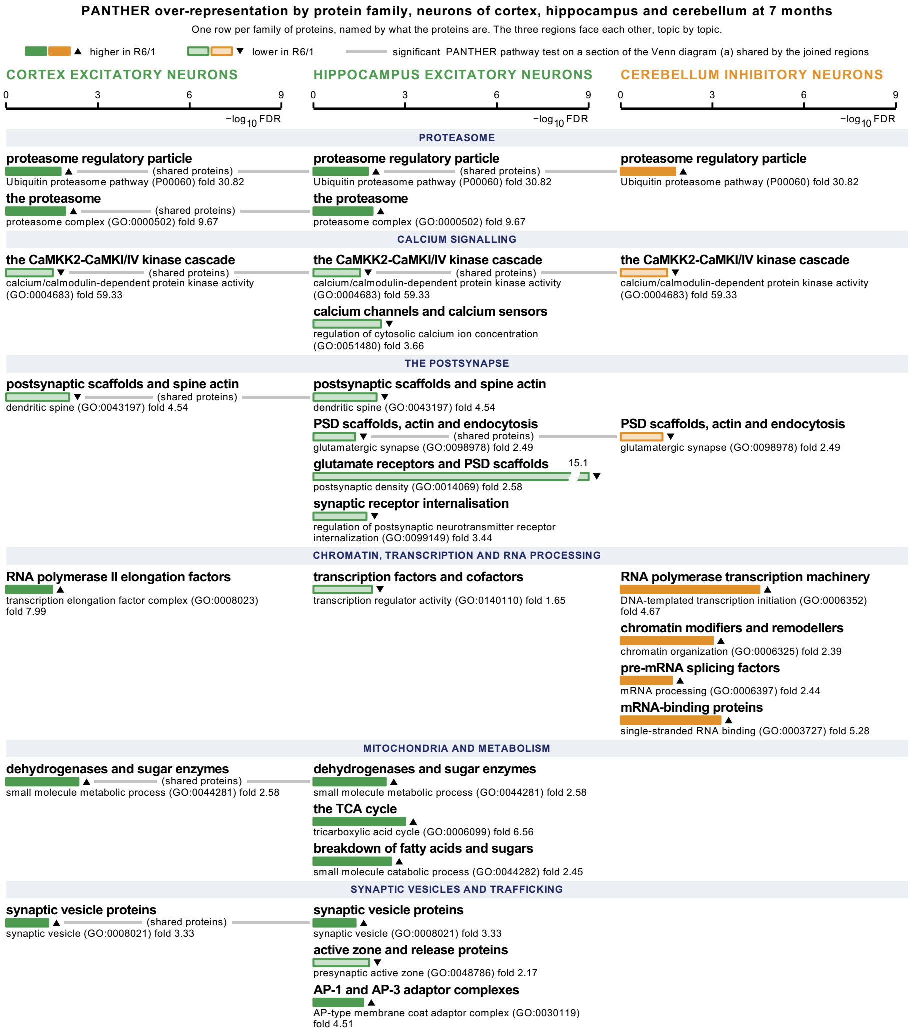


c

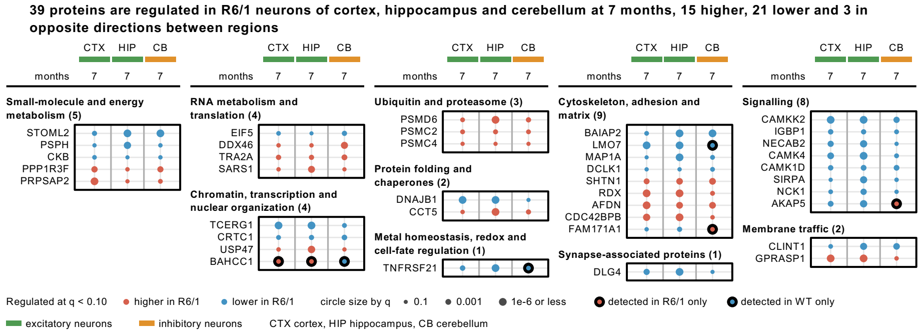

**Supplementary Fig. 8 Protein changes in cortical, hippocampal and cerebellar neurons of R6/1 mice at 7 months**. **(a)** Venn diagram of the protein groups regulated in excitatory neurons of cortex and hippocampus and in inhibitory neurons of cerebellum. In each section, the numbers give the protein groups that were upregulated (red, upward triangle) or downregulated (blue, downward triangle) in R6/1 in every region of the section, or upregulated in one region and downregulated in another (black, diamond). **(b)** Functions over-represented among the proteins upregulated or downregulated in R6/1 in each section of (a), in one column per region (PANTHER, Fisher's exact test with Benjamini-Hochberg correction, FDR < 0.05). Each row is one protein family, with a plain name above the bar and the PANTHER term, its identifier and its fold enrichment under it. Each row shows one over-represented function (a GO term or PANTHER pathway) found in at least five proteins of its Venn section, or at least three in the small central section regulated in all three regions (15 upregulated and 21 downregulated proteins). Functions found mostly in the same proteins are shown only once, except active zone and release proteins, most of which are also in the row glutamate receptors and PSD scaffolds. Solid bars with an upward triangle mark proteins upregulated in R6/1 and pale bars with a downward triangle proteins downregulated, in green for excitatory and in orange for inhibitory neurons. Rows are grouped by topic. A grey line marked (shared proteins) joins the columns of a row that is one test on a section shared by those regions, so its fold enrichment and FDR repeat in each column. An empty column does not mean that the proteins of the row were unchanged in that region. The proteins of every section and their PANTHER results are in Supplementary Table 3. **(c)** The 39 protein groups regulated in all three regions, the central section of (a), 15 upregulated, 21 downregulated and 3 changing direction between regions. Columns are excitatory neurons of cortex and hippocampus (green) and inhibitory neurons of cerebellum (orange) at 7 months. Proteins are grouped by UniProt function, with the number of proteins of each group in brackets. Circles are red when higher and blue when lower in R6/1, and their area increases with −log10 q, as in the key. Black ring, detected in one genotype only, drawn at the largest size. CTX, cortex. HIP, hippocampus. CB, cerebellum. PSD, postsynaptic density. FDR, false discovery rate. n = 4 mice per genotype, except for cortical excitatory neurons (5 R6/1 and 6 WT).

**Supplementary Figure 9**

#### Proteins regulated in R6/1 versus WT at 7 months, neurons and glia

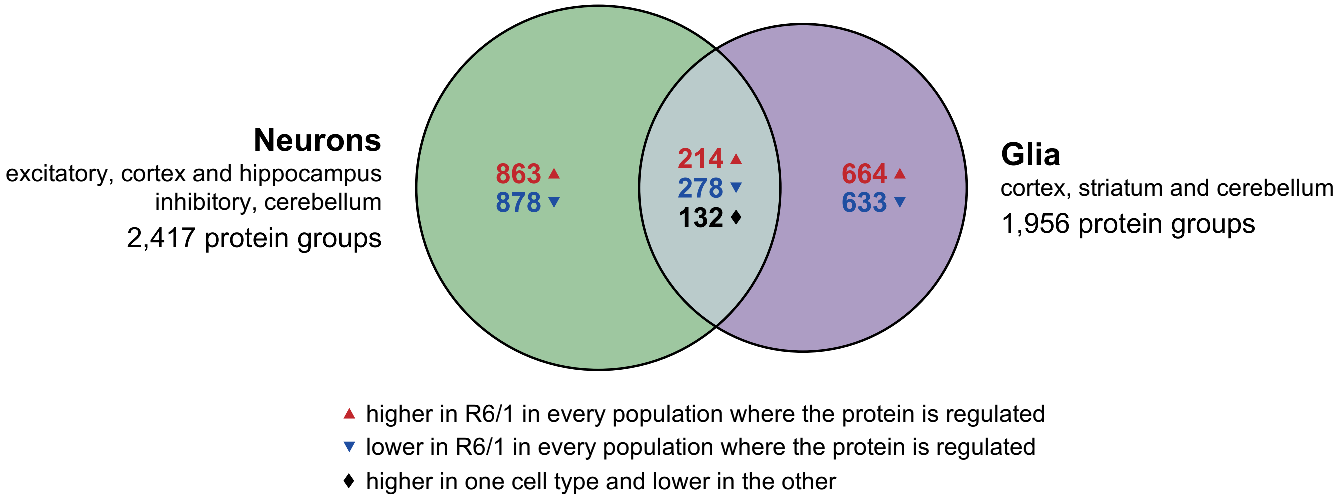


**Supplementary Fig. 9 Proteins regulated in neurons and glia of R6/1 mice at 7 months.** Venn diagram of the protein groups regulated in R6/1 compared with WT littermates in neurons, excitatory neurons of cortex and hippocampus and inhibitory neurons of cerebellum taken together (green), and in glia of cortex, striatum and cerebellum taken together (purple). In each section, the numbers give the protein groups that were upregulated (red, upward triangle) or downregulated (blue, downward triangle) in R6/1 in every population where they were regulated. In the overlap, black (diamond) gives the 132 protein groups upregulated in one cell type and downregulated in the other, 81 downregulated in neurons and upregulated in glia and 51 the reverse. The number under each cell type is the total of protein groups regulated in it, a protein group regulated in opposite directions in two populations of one cell type was counted in the total of each cell type in which it was regulated, but in none of the sections. Supplementary Table 3 lists them in three separate columns. The proteins of every section are listed in Supplementary Table 3. n = 4 mice per genotype, except for cortical excitatory neurons (5 R6/1 and 6 WT).

### Supplementary Figure 10

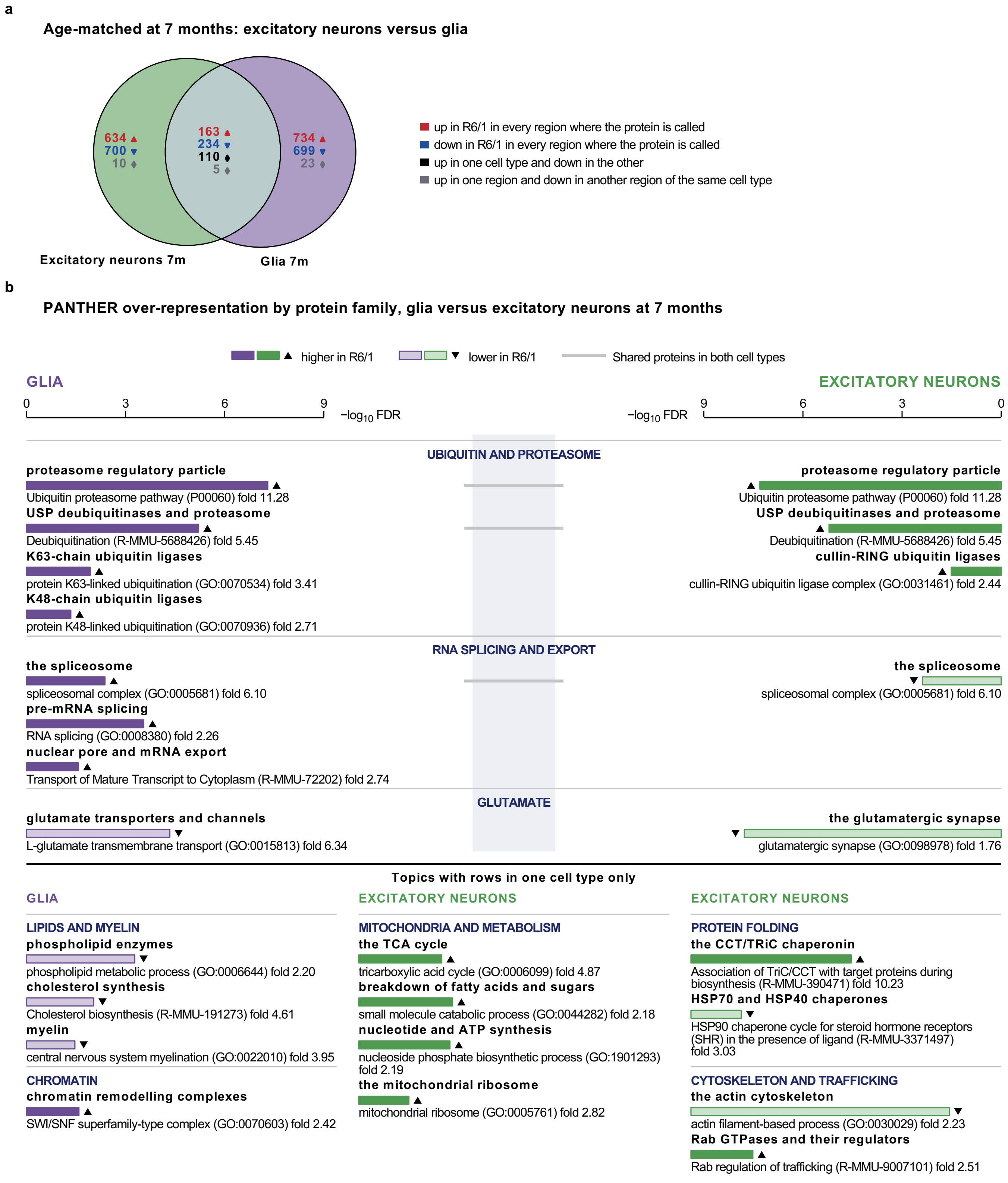

**Supplementary Fig. 10 Protein changes in glia compared with excitatory neurons of R6/1 mice at 7 months**. **(a)** Venn diagram of the protein groups regulated in R6/1 in excitatory neurons of cortex and hippocampus together (green) and in glia of cortex, striatum and cerebellum together (purple). In each section, the numbers give the protein groups that were upregulated (red, upward triangle) or downregulated (blue, downward triangle) in R6/1 in every region, upregulated in one cell type and downregulated in the other (black, diamond), or upregulated in one region and downregulated in another region of the same cell type (grey, diamond). **(b)** Functions over-represented among the proteins of each section of (a), glia on the left and excitatory neurons on the right (PANTHER, Fisher's exact test with Benjamini-Hochberg correction, FDR < 0.05). Each row is one protein family, with a plain name above the bar and the PANTHER term, its identifier and its fold enrichment under it. Bar length is −log10 FDR, on one scale for the whole panel. Solid bars with an upward triangle mark proteins upregulated and pale bars with a downward triangle proteins downregulated in R6/1, in purple for glia and in green for excitatory neurons. A grey line joins the two sides of a row that is one test on a section shared by both cell types, so its fold enrichment and FDR repeat on each side. The row HSP70 and HSP40 chaperones is named after 8 of its 14 proteins, which the Reactome pathway HSP90 chaperone cycle for steroid hormone receptors (R-MMU-3371497) groups together. The proteins of every section and their PANTHER results are in Supplementary Table 3. FDR, false discovery rate. n = 4 mice per genotype, except for cortical excitatory neurons (5 R6/1 and 6 WT).

**Supplementary File 1. Quality control of the cell-type-specific proteomes.** For each population, by region and age, principal component analysis of the samples with the no-ANL controls, protein intensity and number of protein groups in each sample, enrichment of each genotype over the no-ANL controls, the enriched and exclusive proteomes, and the mean log2 protein intensity of the cell-type-specific proteome in each genotype. Peptide intensity distributions are shown once per region and cell type. Brackets compare the per-sample median log2 intensities of the two groups they join (two-sided Welch t-test, unadjusted p). One, two, three and four asterisks mark $p \leq 0.05$, 0.01, 0.001 and 0.0001, and ns marks $p > 0.05$. In the peptide panels of neurons, each genotype at each age is compared with the no-ANL controls. Samples are labelled + for R6/1, − for WT and NC for the no-ANL controls. CTX, cortex. HIP, hippocampus. CB and CRB, cerebellum. STR, striatum. m, months. n = 4 mice per genotype, except for cortical excitatory neurons (6 R6/1 and 6 WT at 1.5 months, 5 and 5 at 3 months, 5 R6/1 and 6 WT at 7 months).

**Supplementary File 2. Proteins of the protein sets of Fig. 9. (a)** Antiviral response, viral restriction and interferon-responsive sets. **(b)** Proteasome, 20S core and 19S and accessory regulators. **(c)** Chaperones, DnaJ domain, protein refolding and CCT/TRiC. **(d)** Cytosolic ribosome. **(e)** Mitochondrial catabolism, TCA cycle and branched-chain amino acid catabolism on the first page and fatty-acid beta-oxidation in mitochondria on the second. Rows are the member proteins of each set and columns the comparisons of R6/1 with WT, as in Fig. 9, with glia at 7 months. Protein cells are coloured by the log2 fold change (R6/1/WT), as in the scale bar, and an asterisk marks the proteins regulated in R6/1. Dark red and dark blue cells with a triangle mark proteins upregulated in R6/1 (detected mainly in R6/1, in at most one WT mouse) or downregulated in R6/1 (detected mainly in WT, in at most one R6/1 mouse), all listed in Supplementary Table 1. These proteins have no fold change and did not enter the set test. Grey cells mark proteins not quantified or with no estimate in that comparison. The bottom rows give the cameraPR result of each set, as in Fig. 9, coloured by the signed −log10 FDR, red when the set is higher and blue when it is lower in R6/1, with an asterisk at FDR < 0.05. n.t., not tested. FDR, false discovery rate. CTX, cortex. HIP, hippocampus. CB, cerebellum. STR, striatum. m, months. n = 4 mice per genotype, except for cortical excitatory neurons (6 R6/1 and 6 WT at 1.5 months, 5 and 5 at 3 months, 5 R6/1 and 6 WT at 7 months).
