## Supplementary material for "Excitatory neurons downregulate synaptic proteins before motor onset and glia reduce neuronal support in Huntington’s disease mice": MS-DAP Files

### Excitatory neurons, cortex, 1.5 months

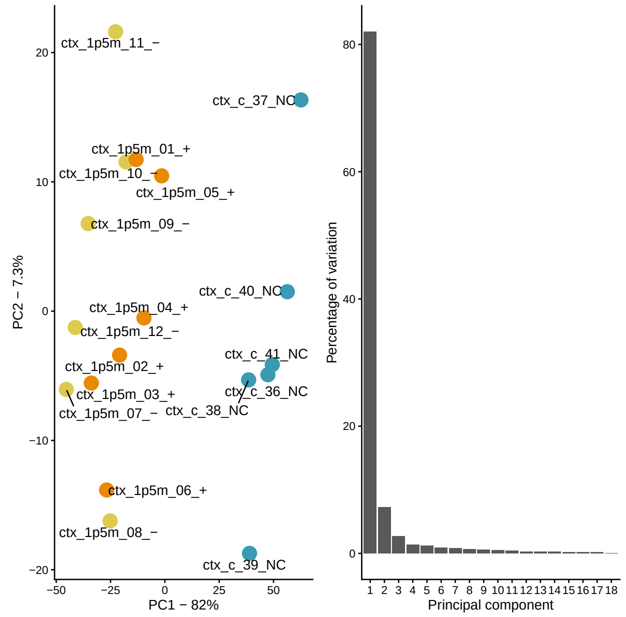


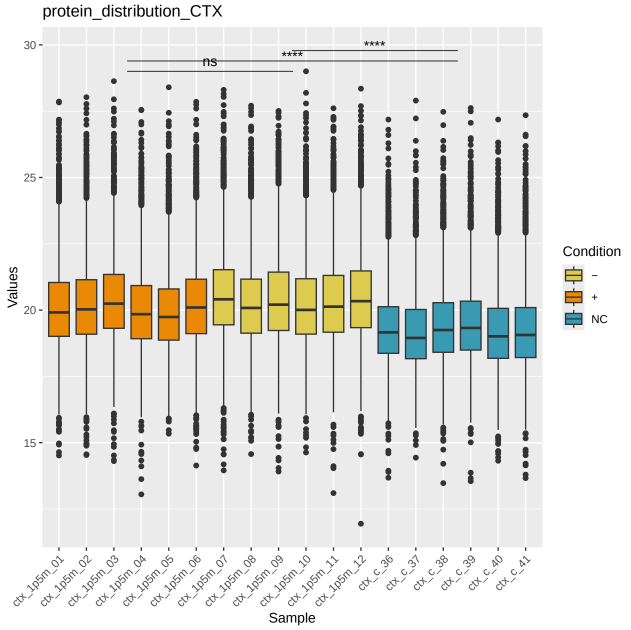


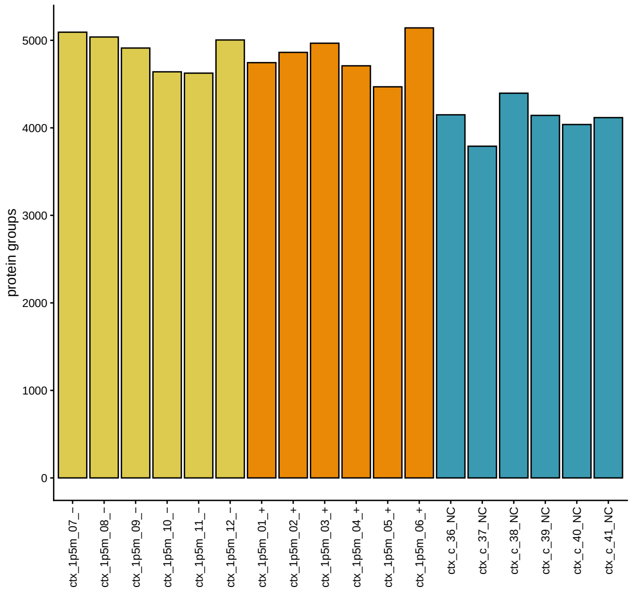


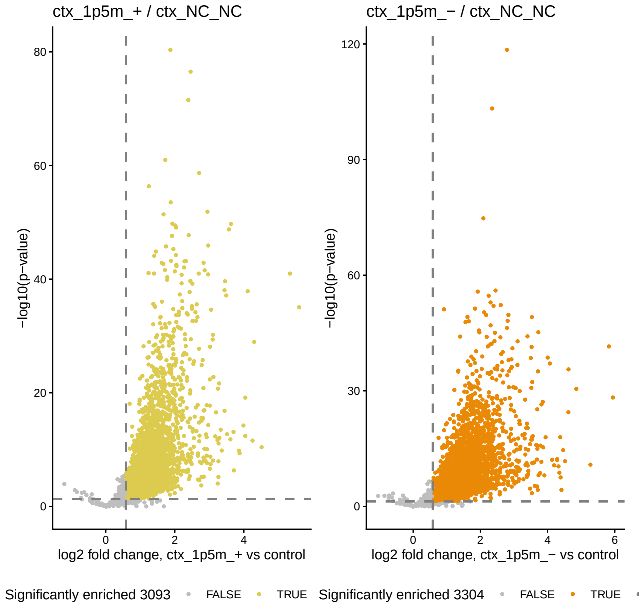


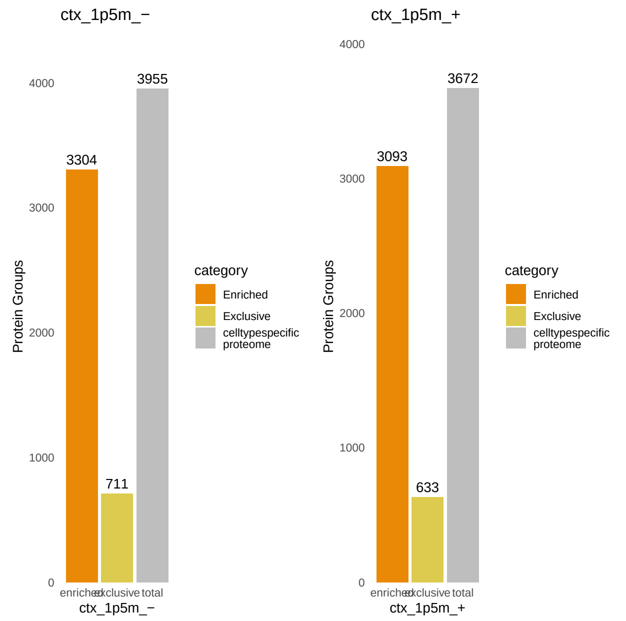


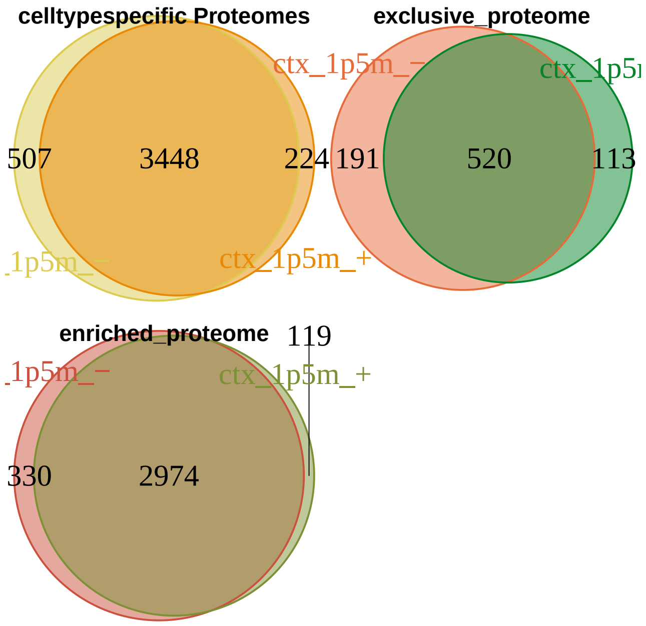

#### Excitatory neurons, cortex, 1.5 months, continued

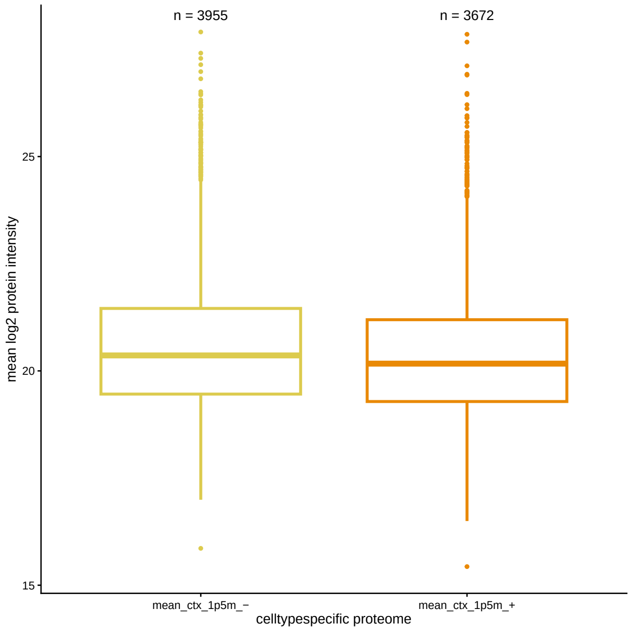


#### Excitatory neurons, cortex, 3 months

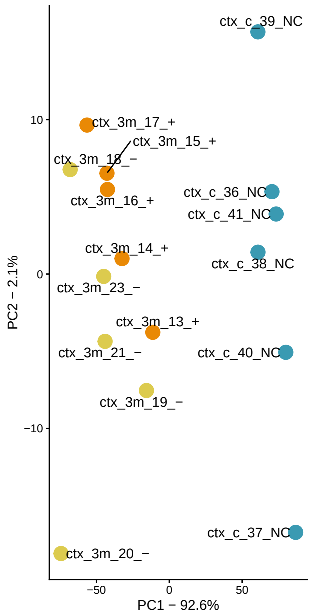


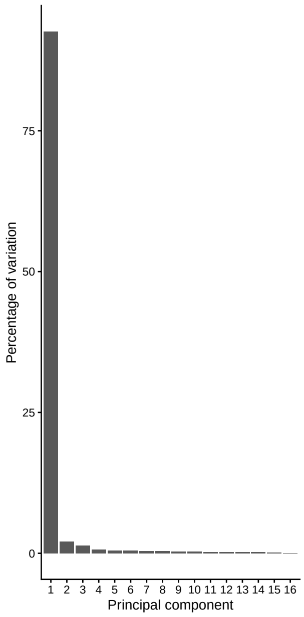


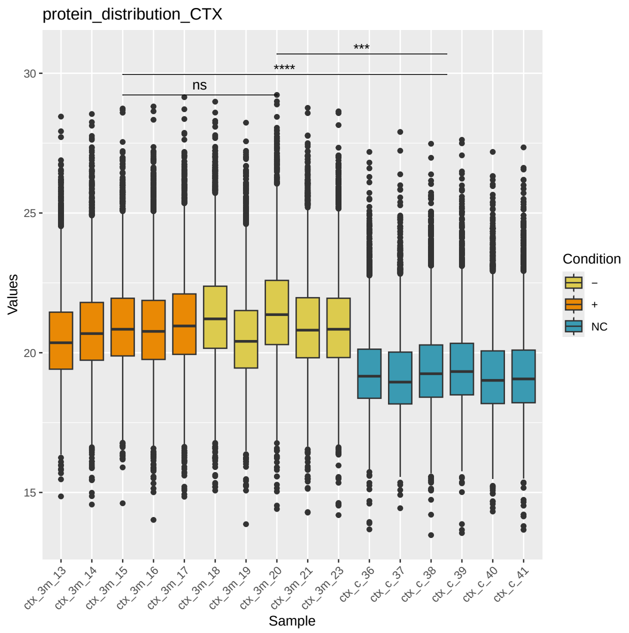


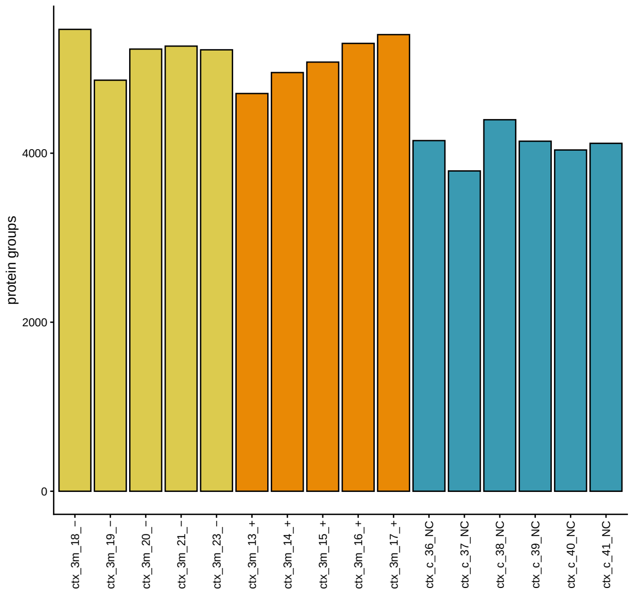


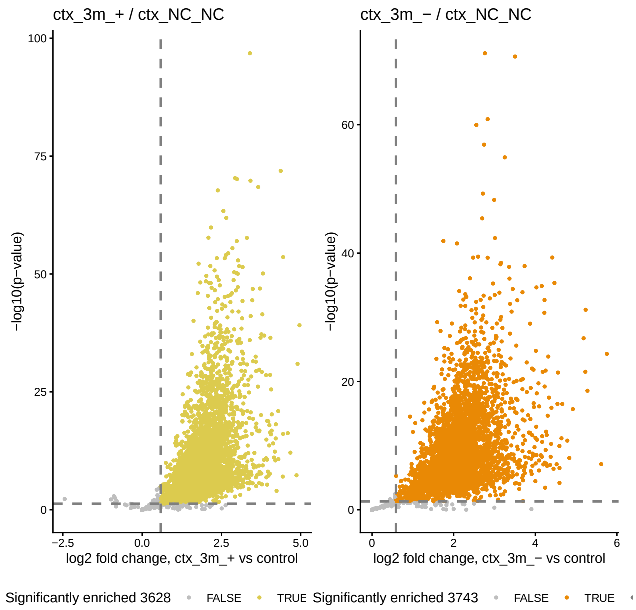

#### Excitatory neurons, cortex, 3 months, continued

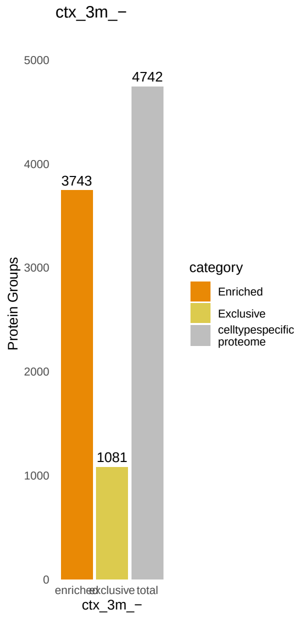


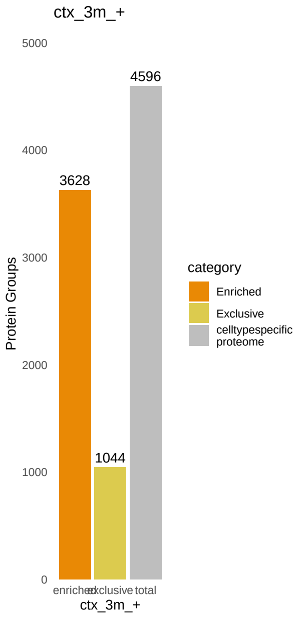


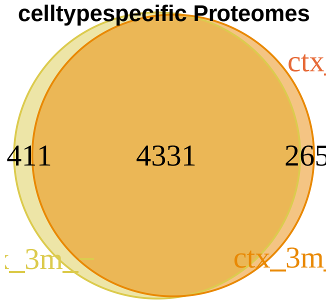


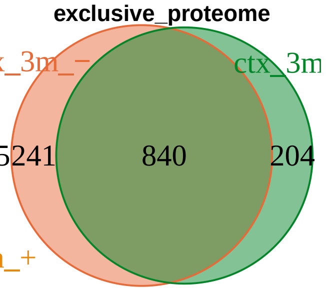


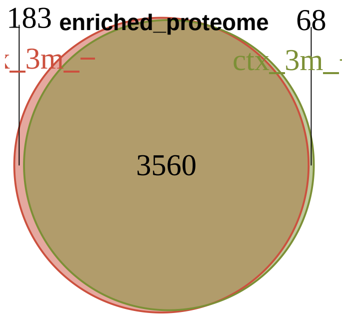


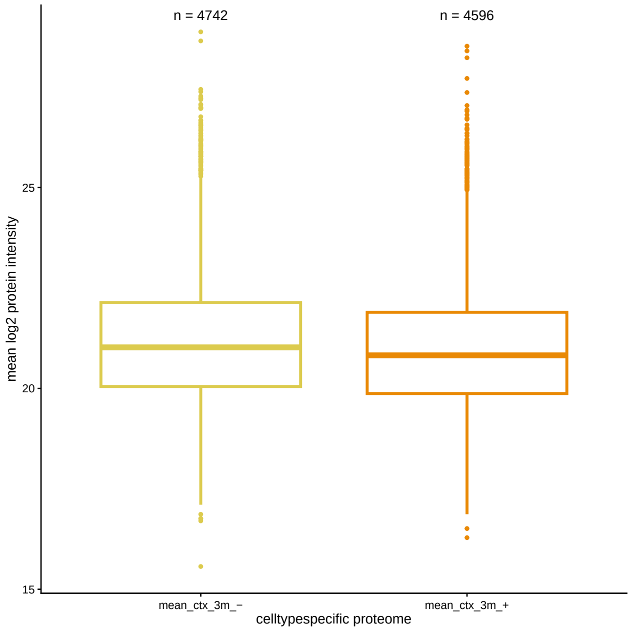


#### Excitatory neurons, cortex, 7 months

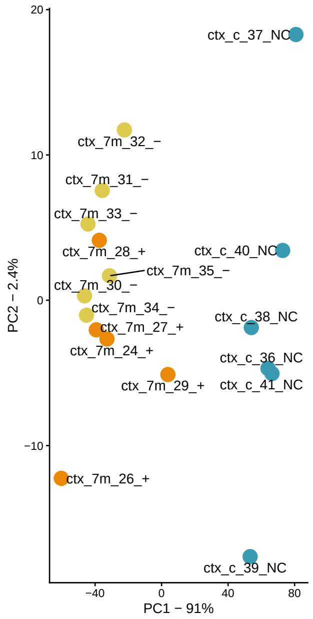


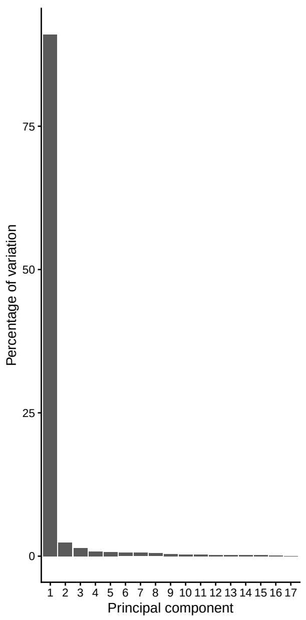


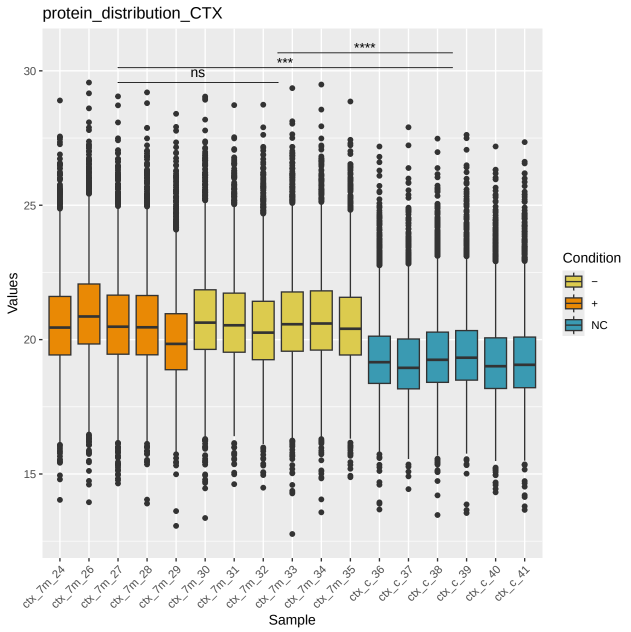

#### Excitatory neurons, cortex, 7 months, continued

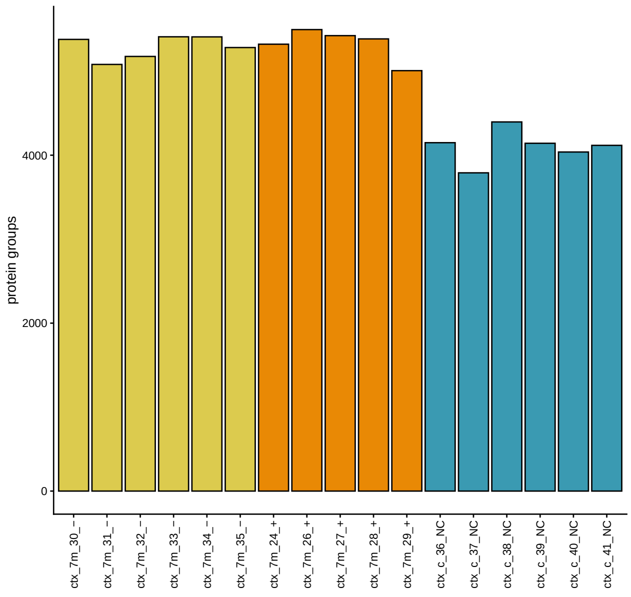


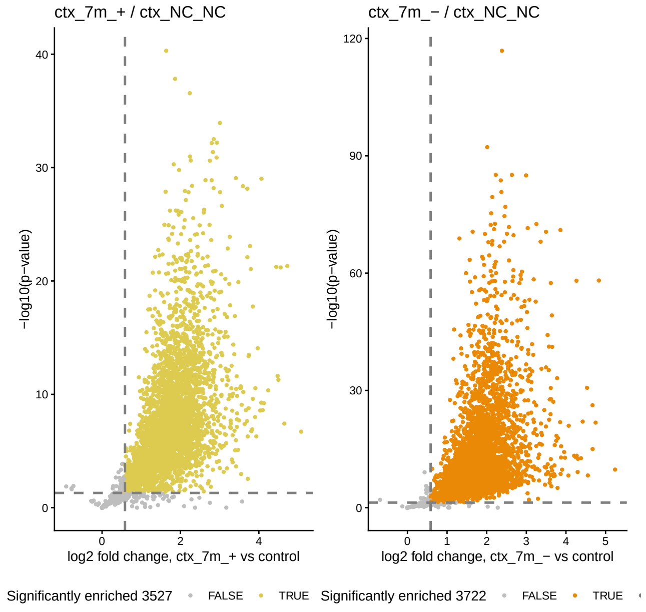


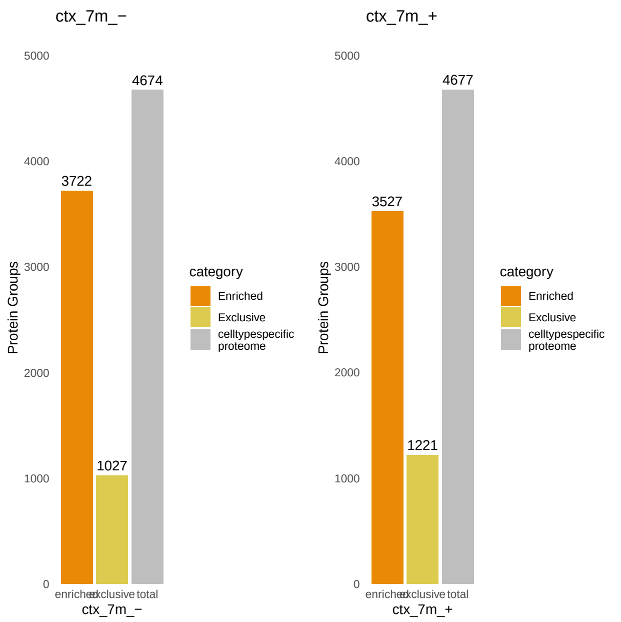


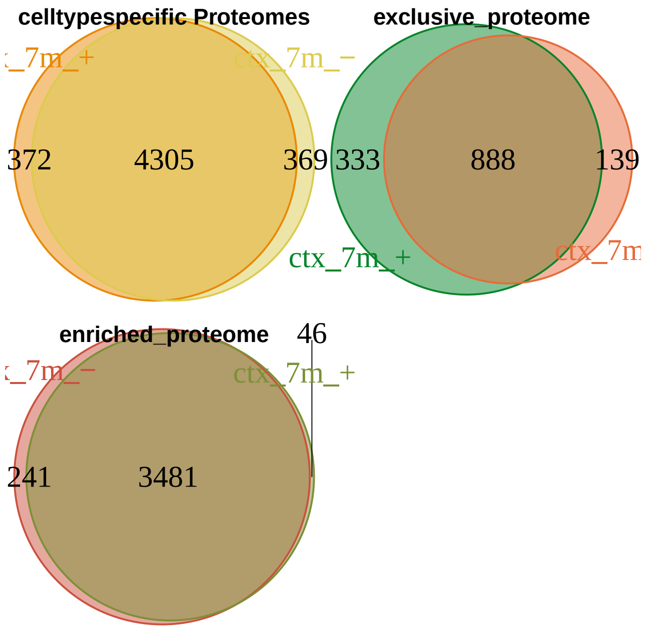


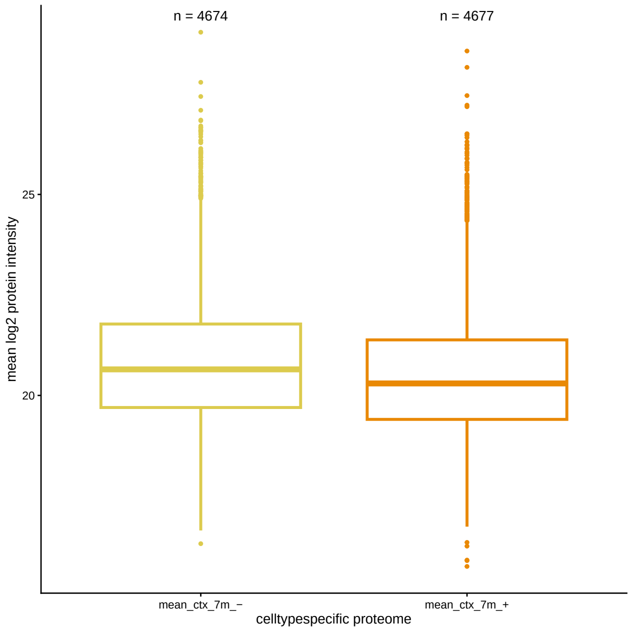

#### Excitatory neurons, cortex, 1.5, 3 and 7 months together, peptide intensities

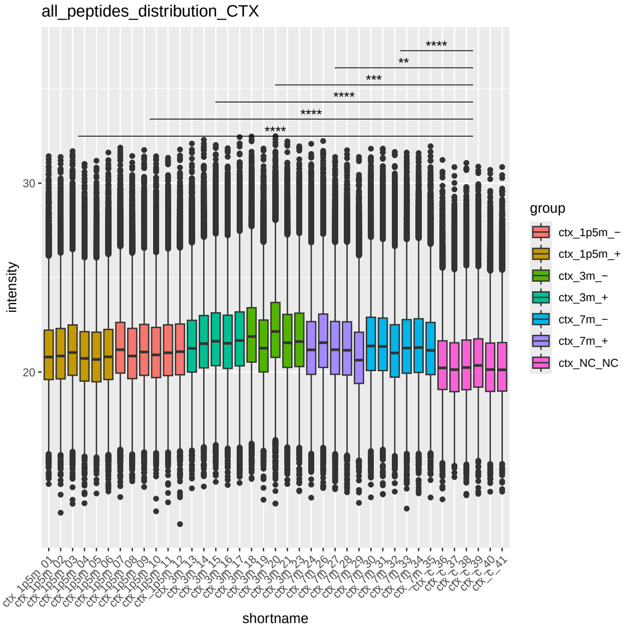


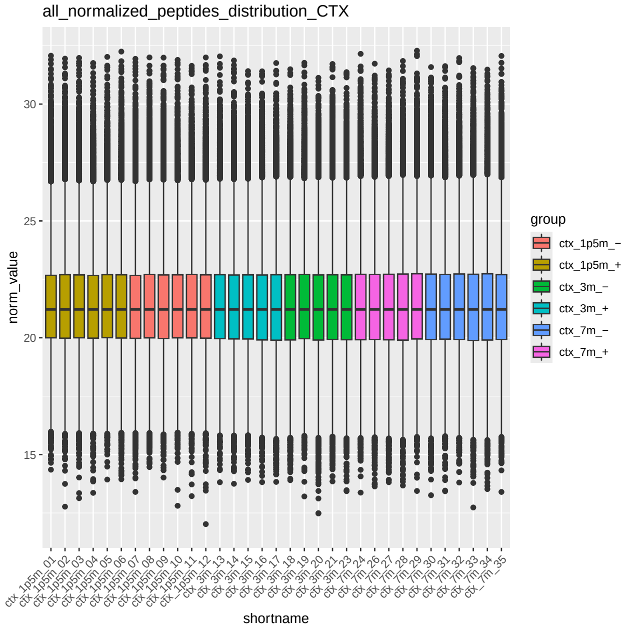


#### Excitatory neurons, hippocampus, 1.5 months

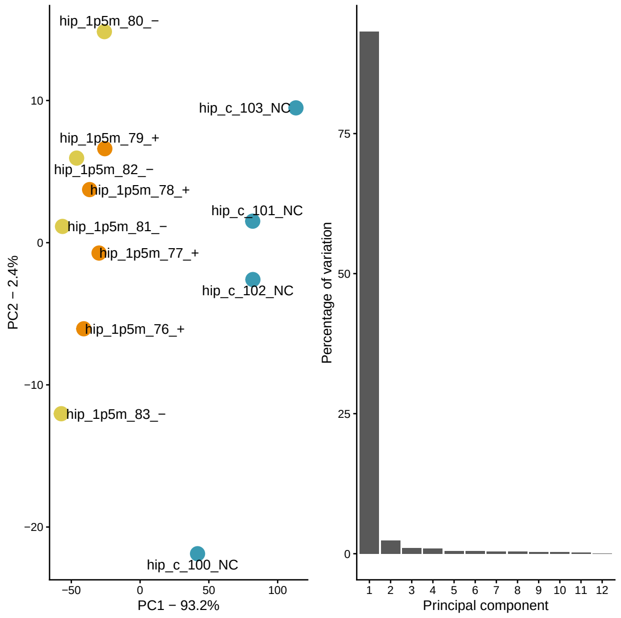


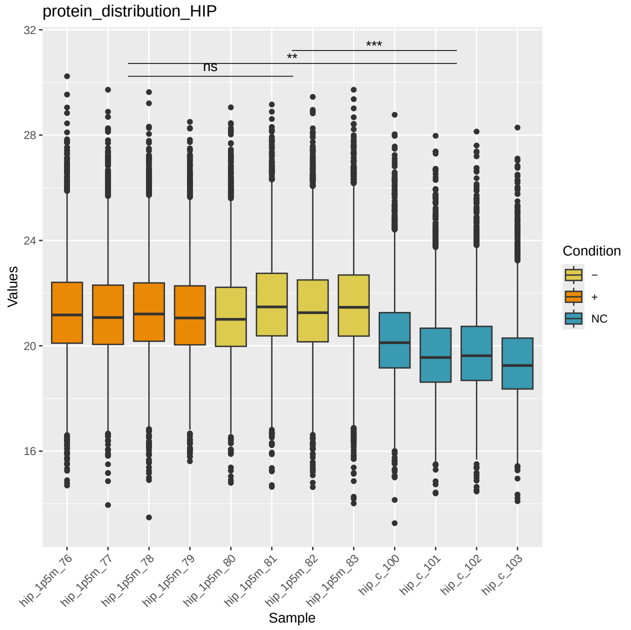


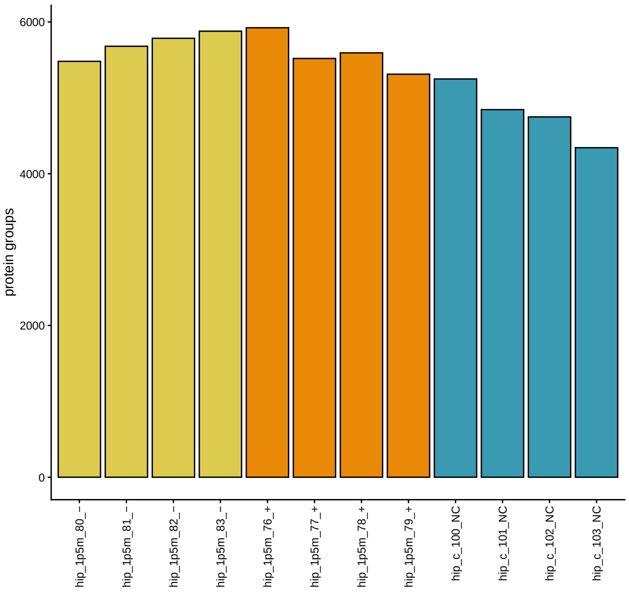


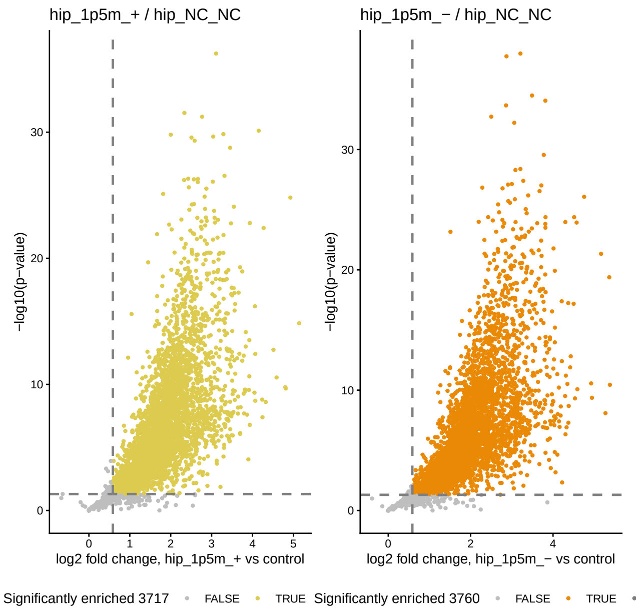

#### Excitatory neurons, hippocampus, 1.5 months, continued

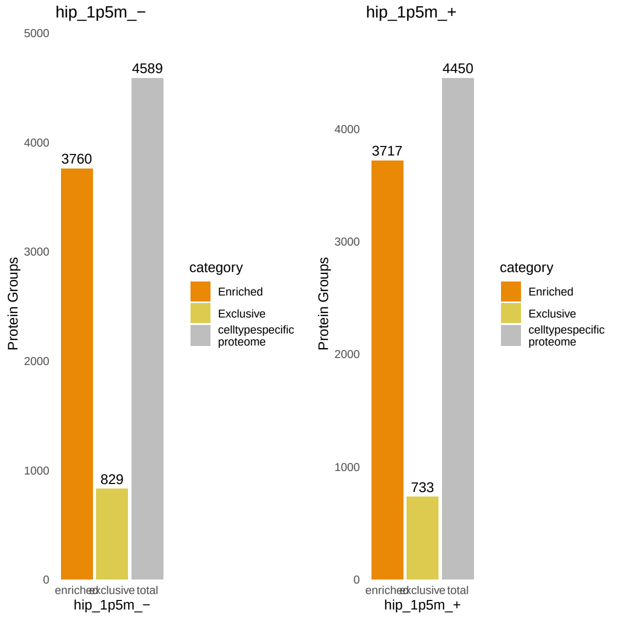


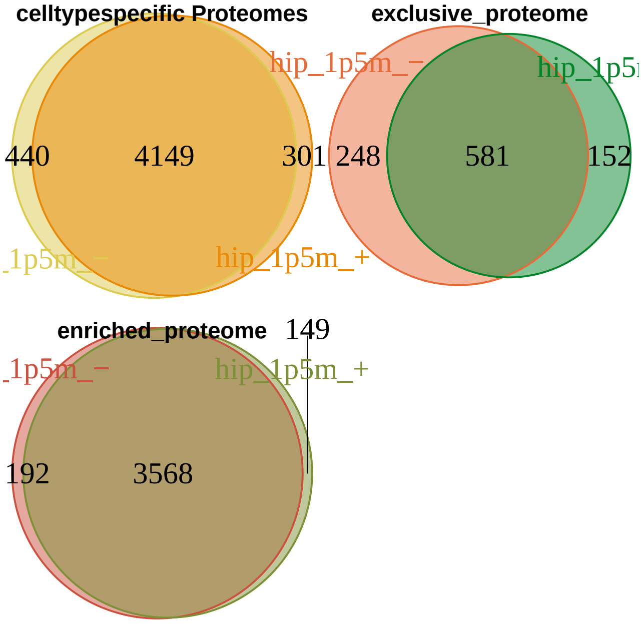


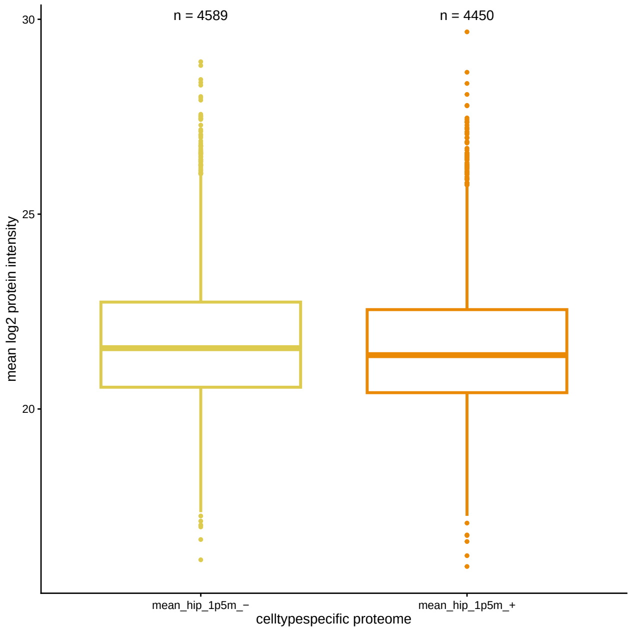


#### Excitatory neurons, hippocampus, 3 months

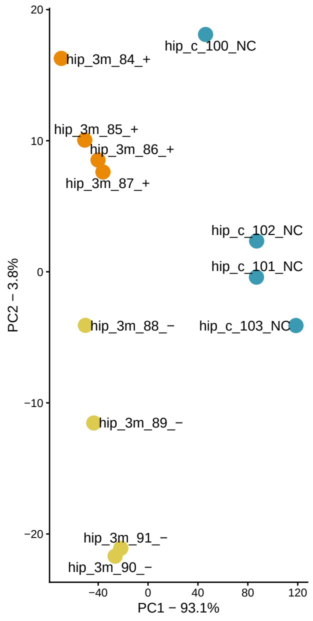


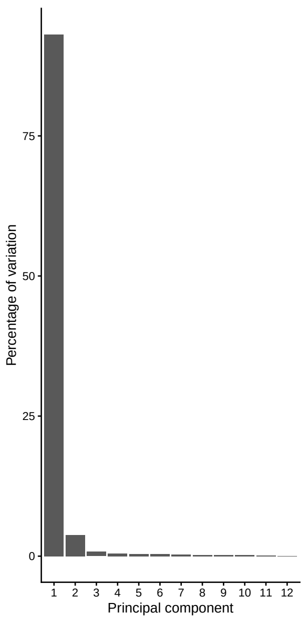


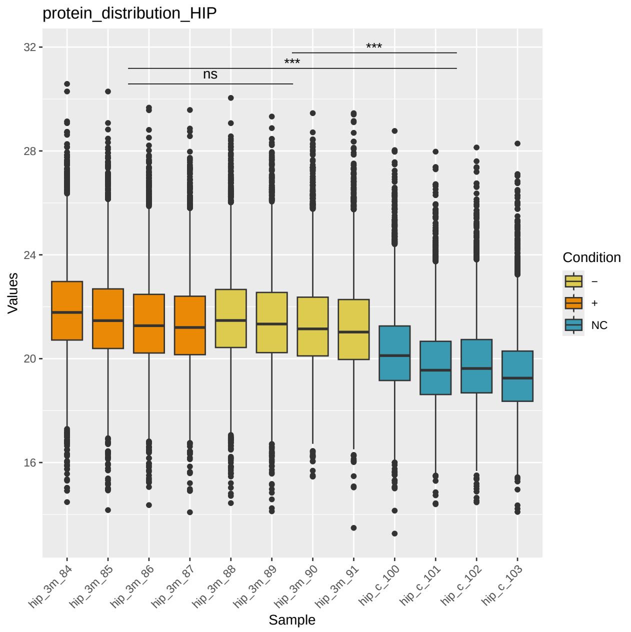

#### Excitatory neurons, hippocampus, 3 months, continued

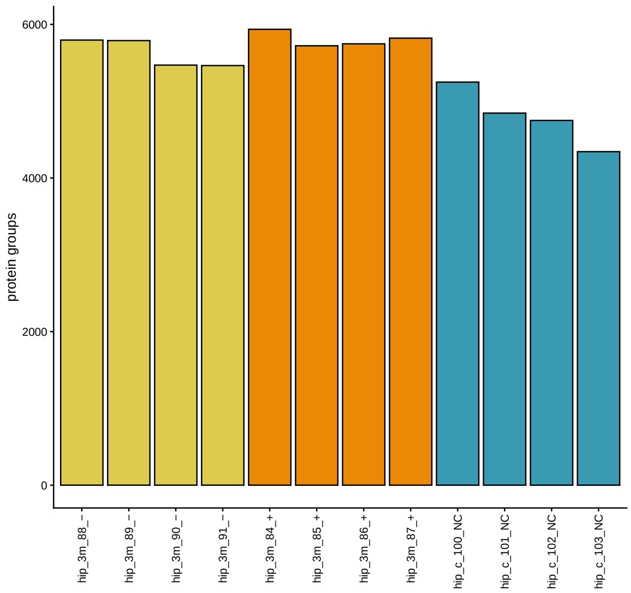


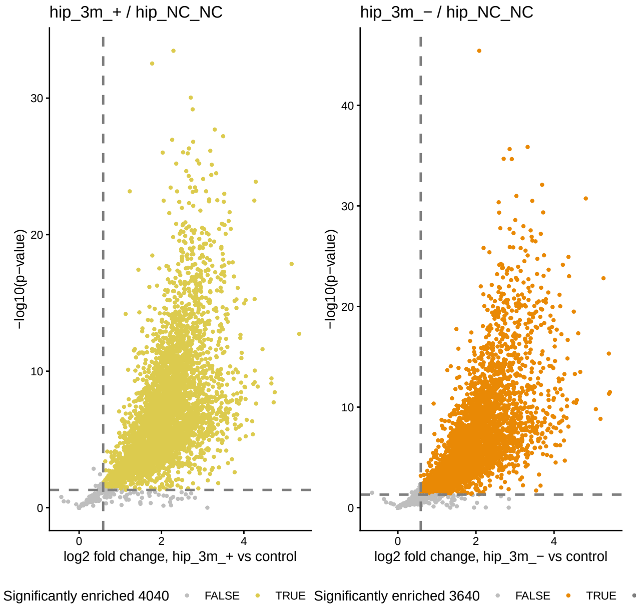


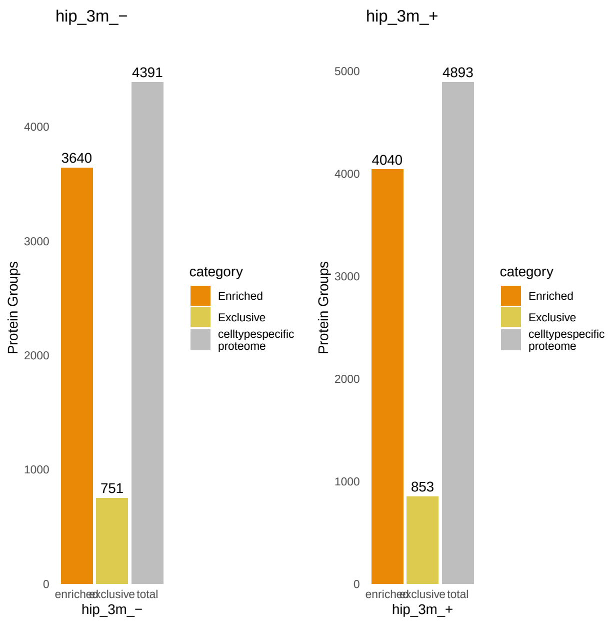


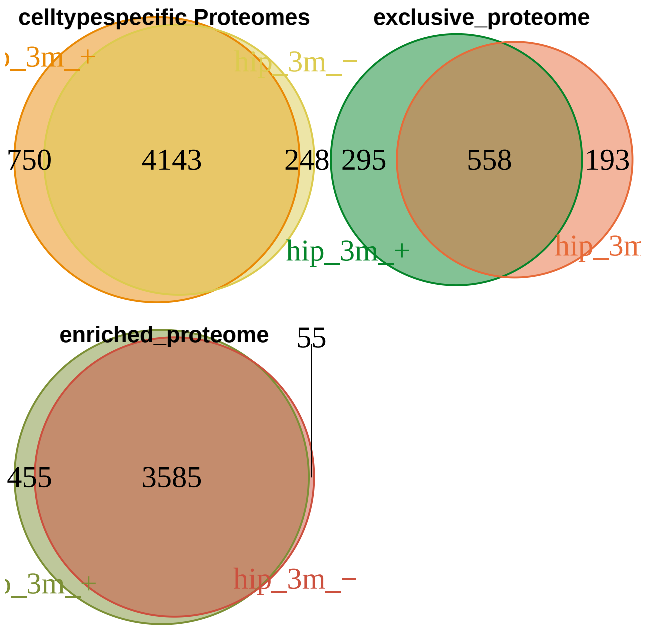


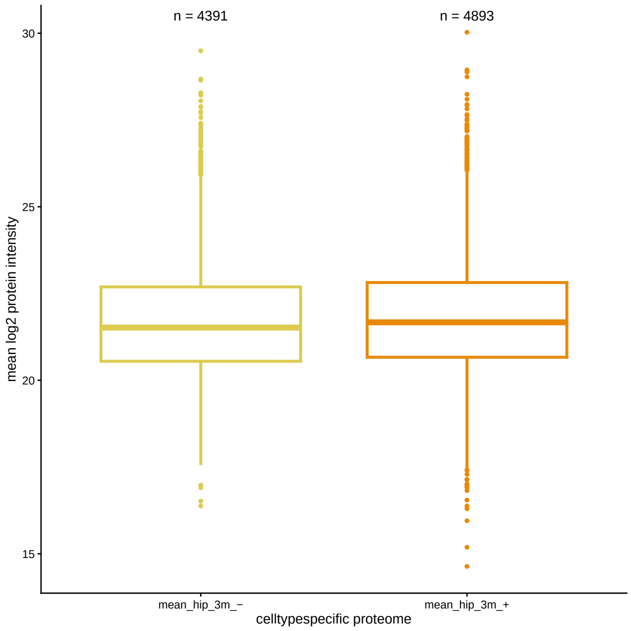

#### Excitatory neurons, hippocampus, 7 months

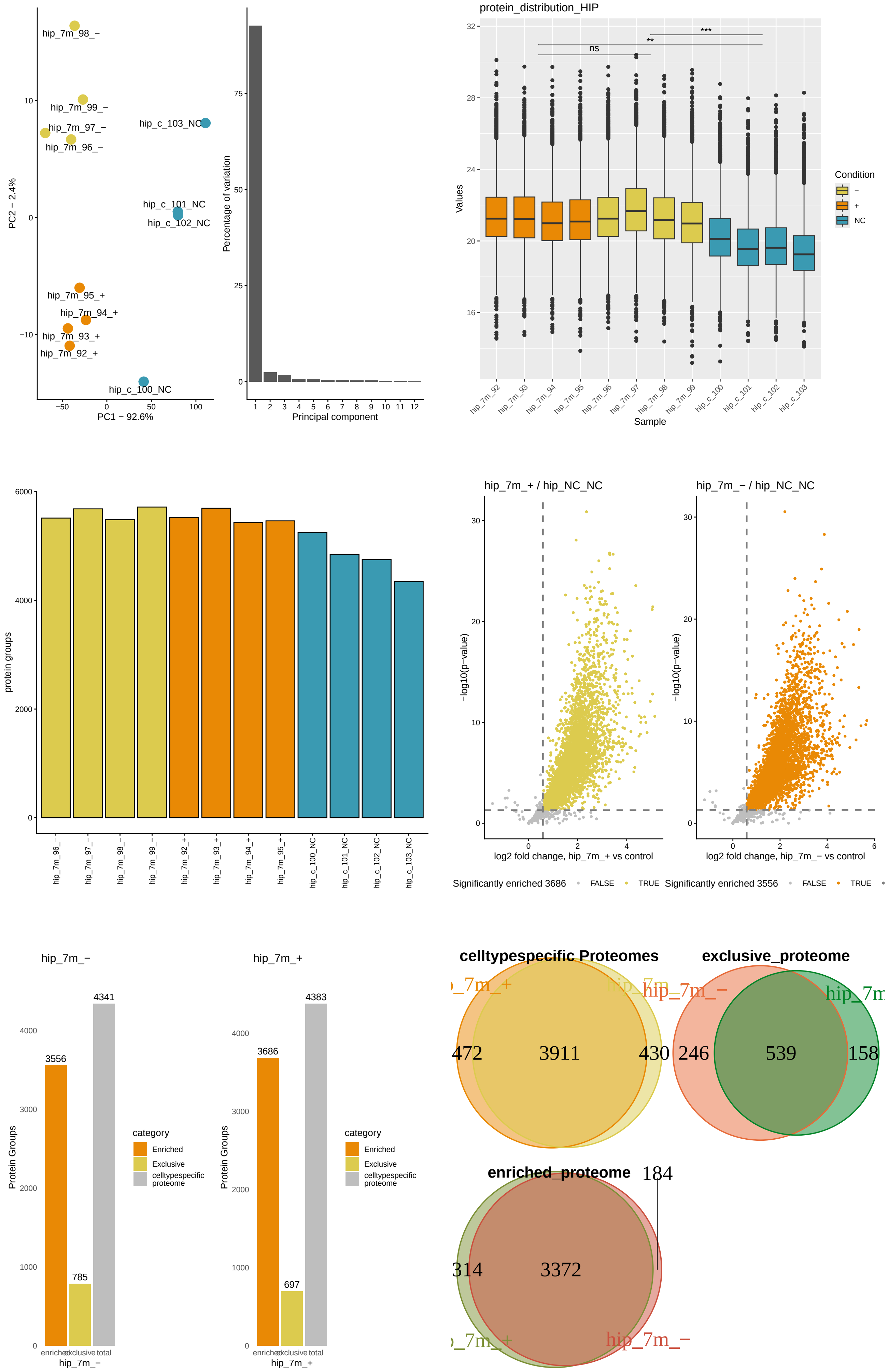

#### Excitatory neurons, hippocampus, 7 months, continued

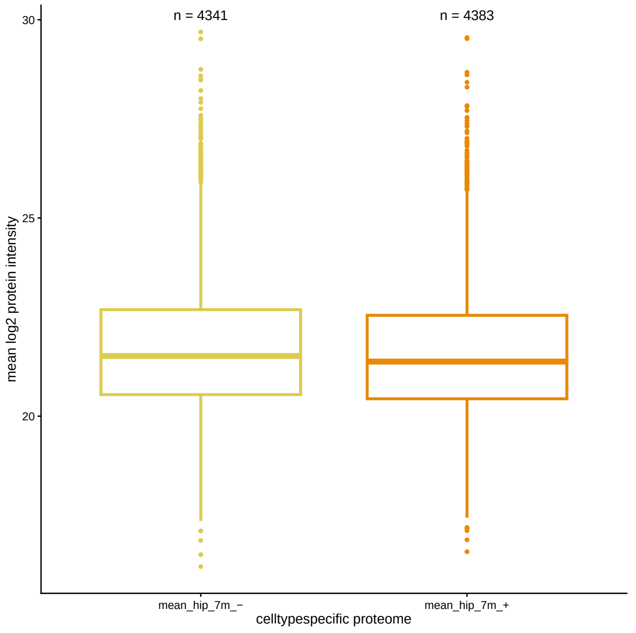


#### Excitatory neurons, hippocampus, 1.5, 3 and 7 months together, peptide intensities

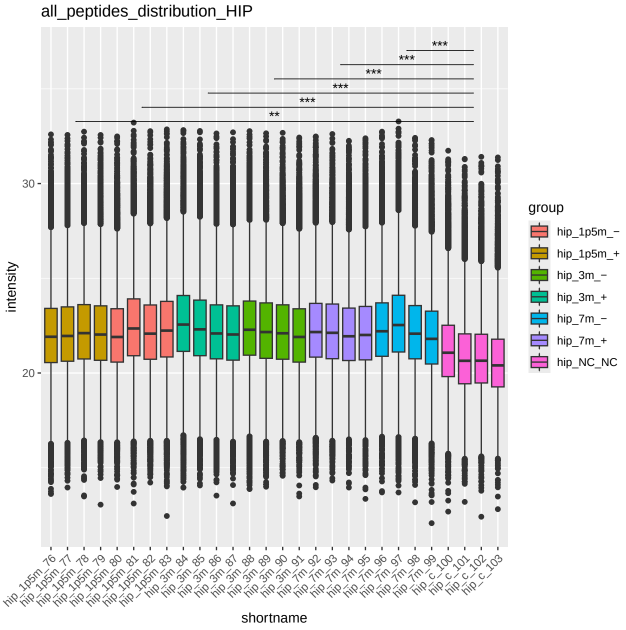


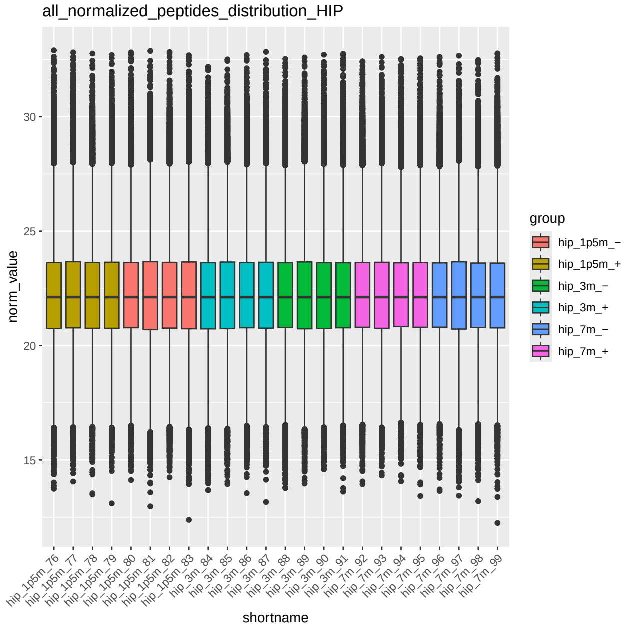


#### Inhibitory neurons, cerebellum, 3 months

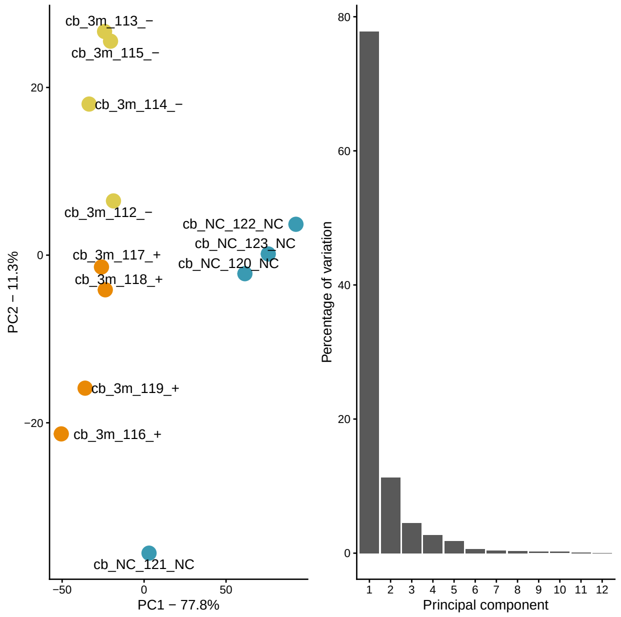


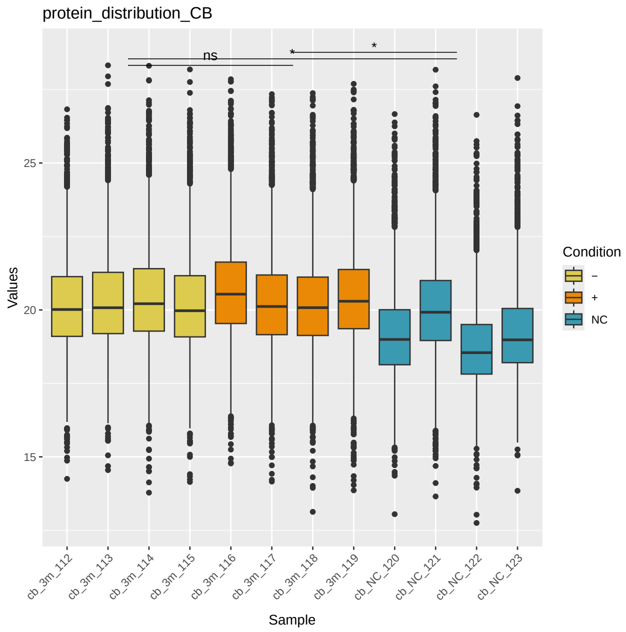

#### Inhibitory neurons, cerebellum, 3 months, continued

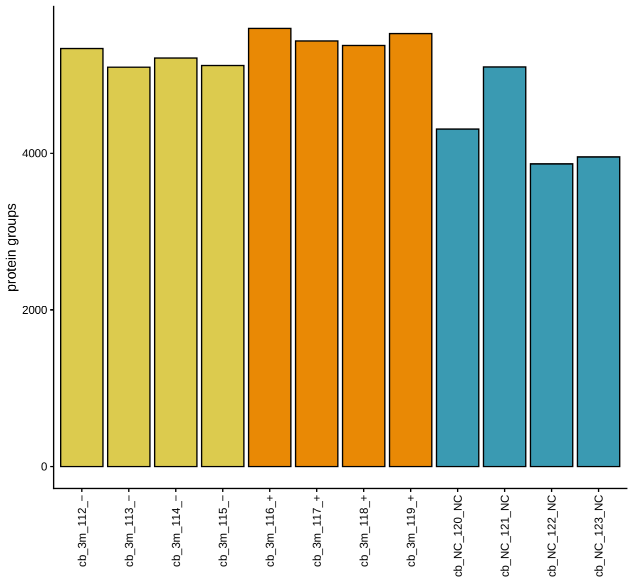


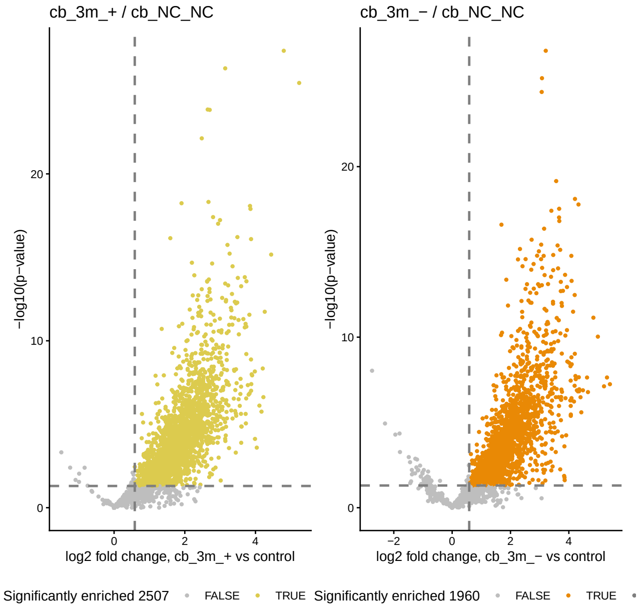


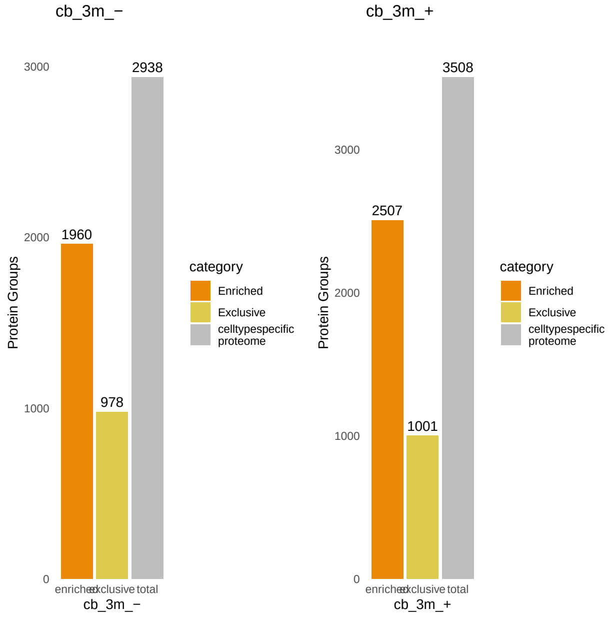


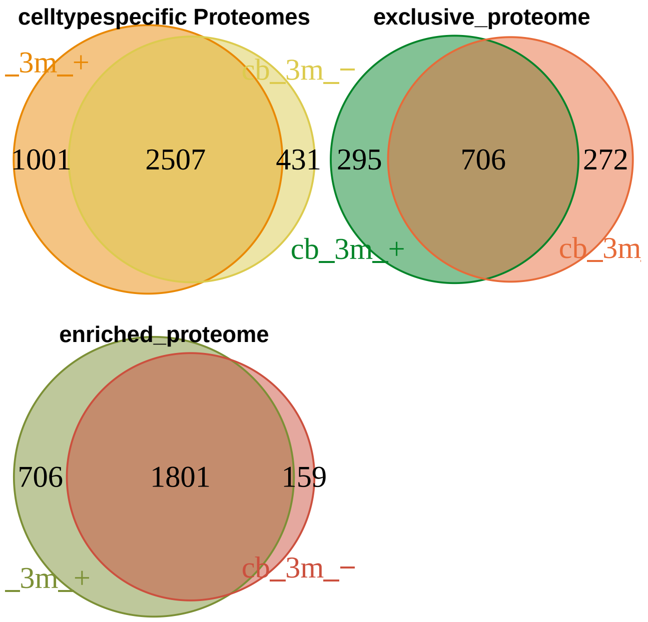


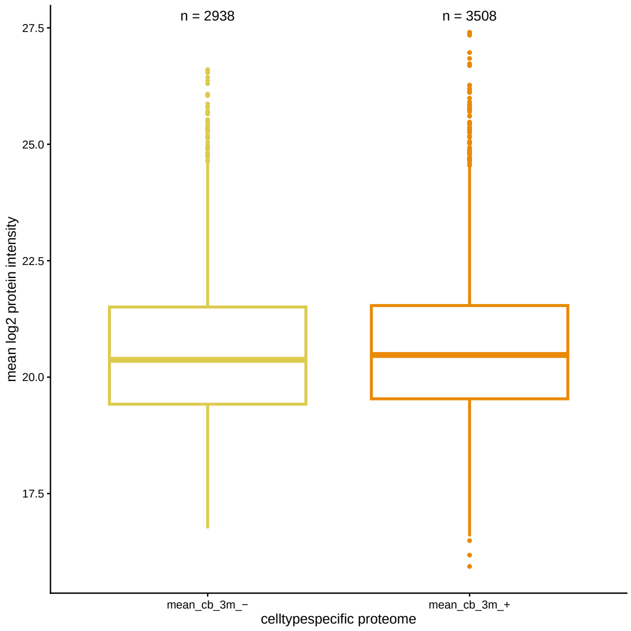

#### Inhibitory neurons, cerebellum, 7 months

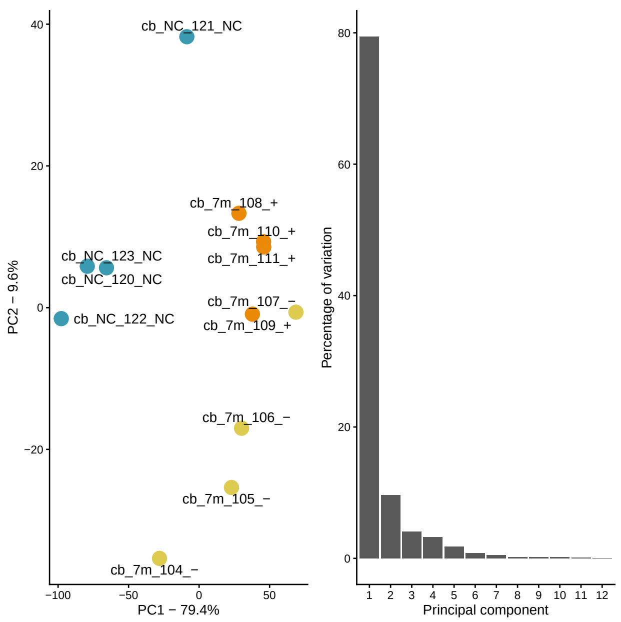


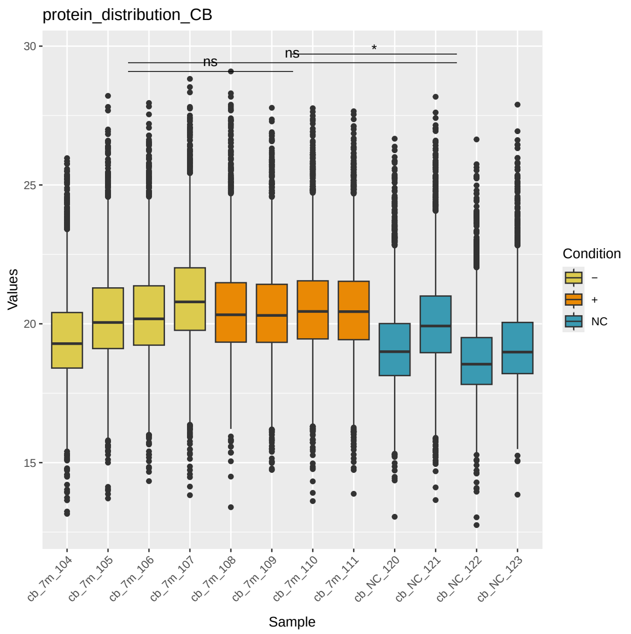


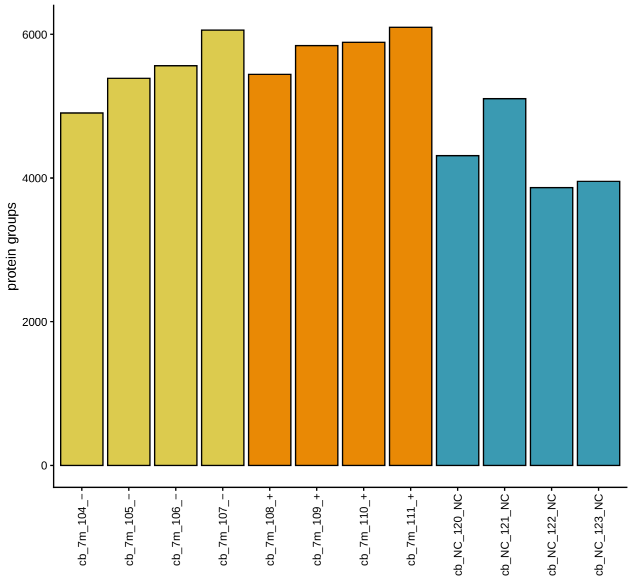


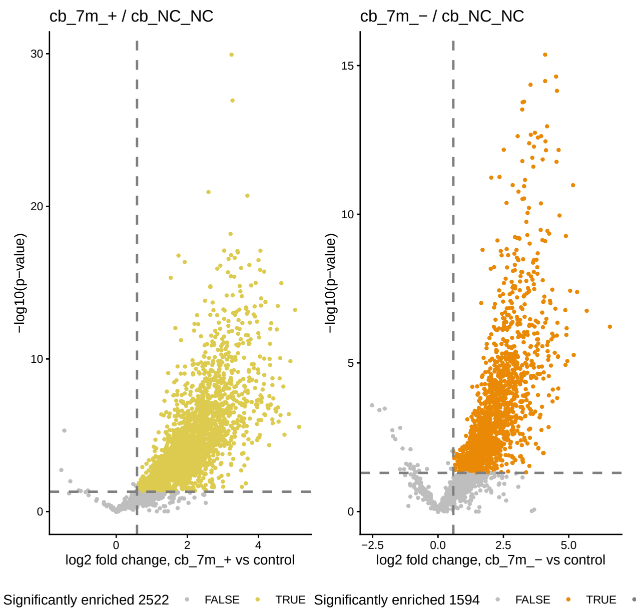


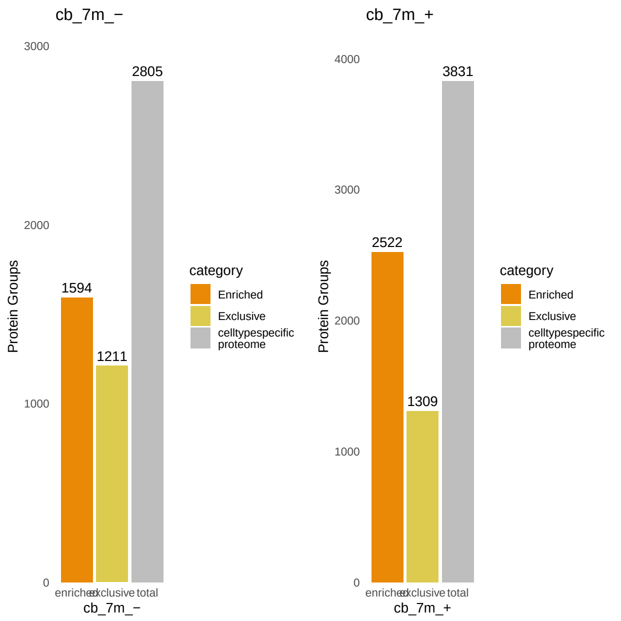


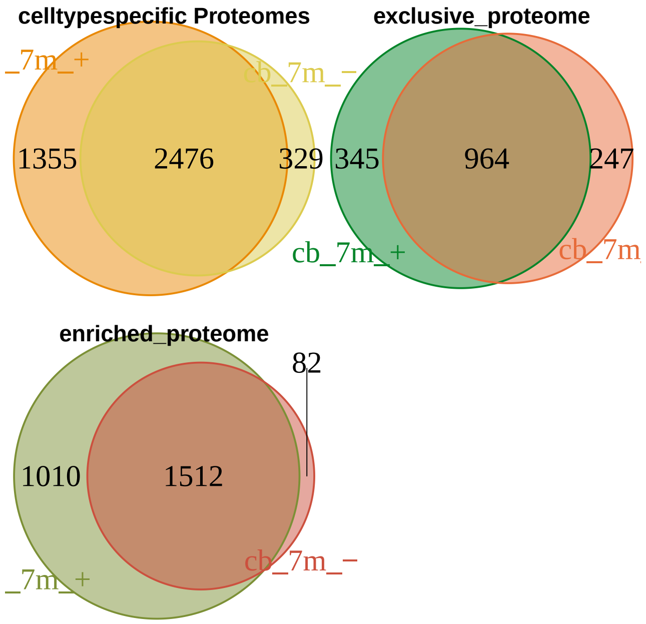

#### Inhibitory neurons, cerebellum, 7 months, continued

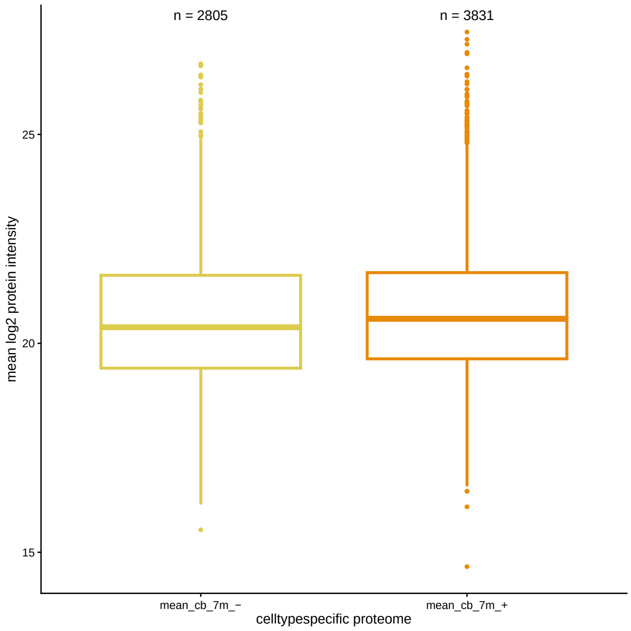


#### Inhibitory neurons, cerebellum, 3 and 7 months together, peptide intensities

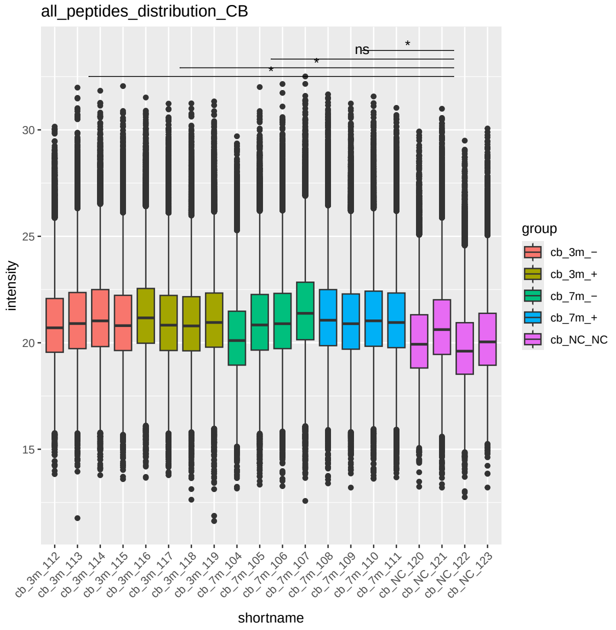


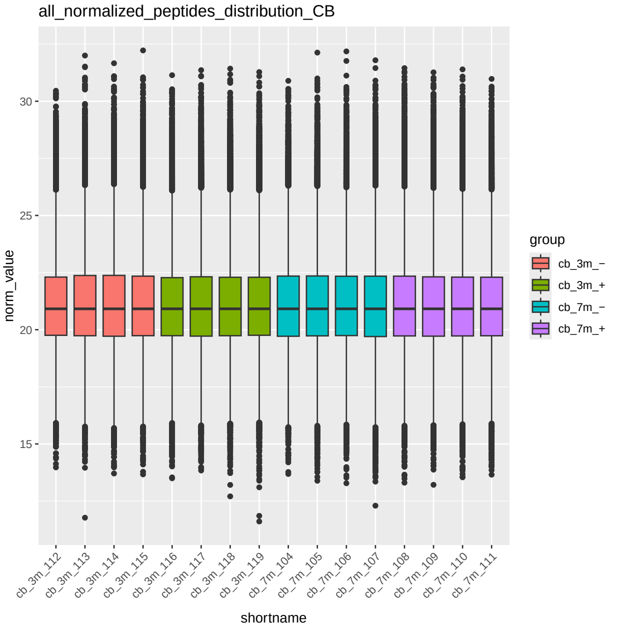


#### Glia, cortex, 7 months

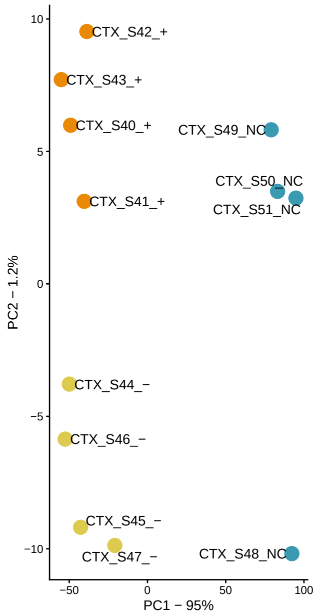


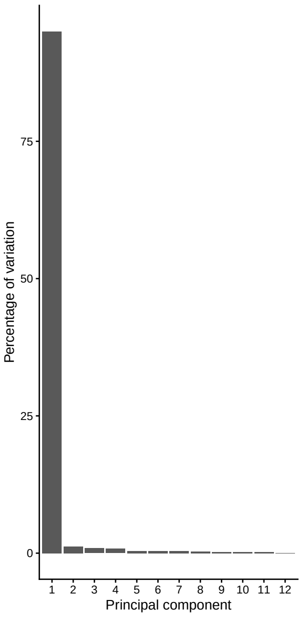


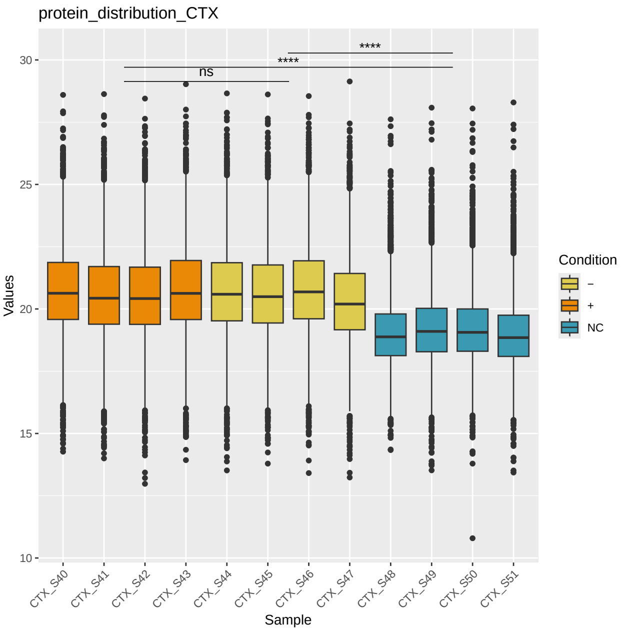

#### Glia, cortex, 7 months, continued

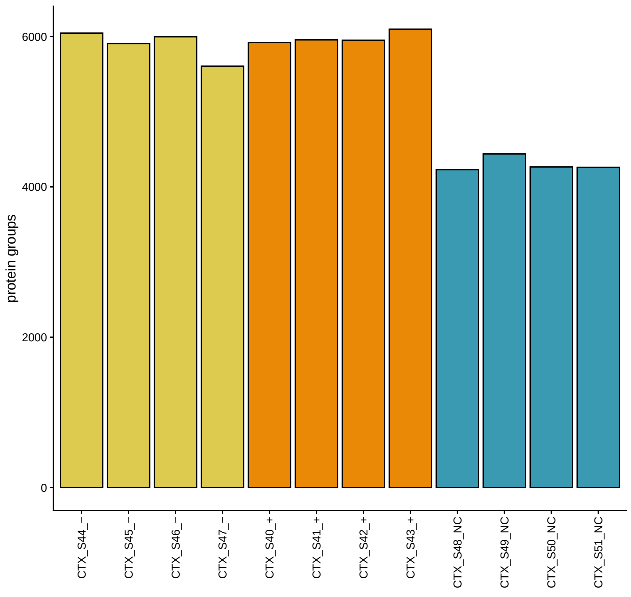


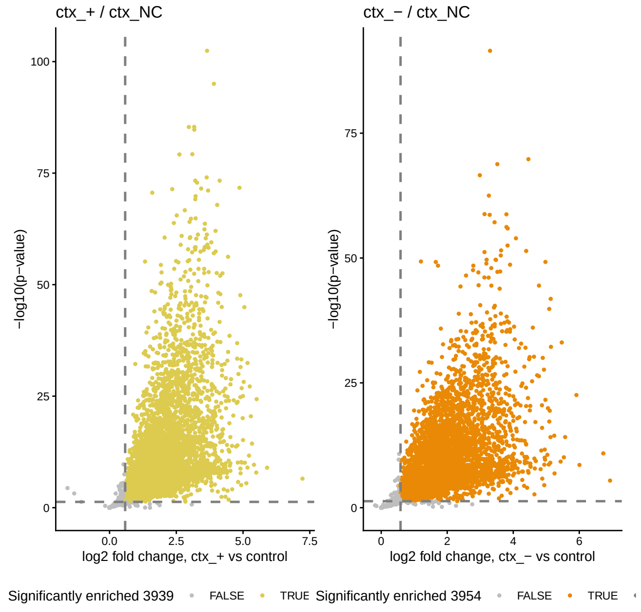


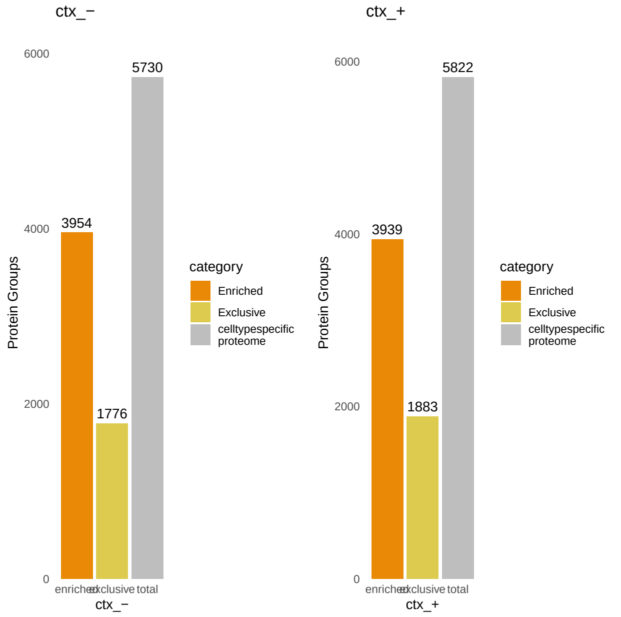


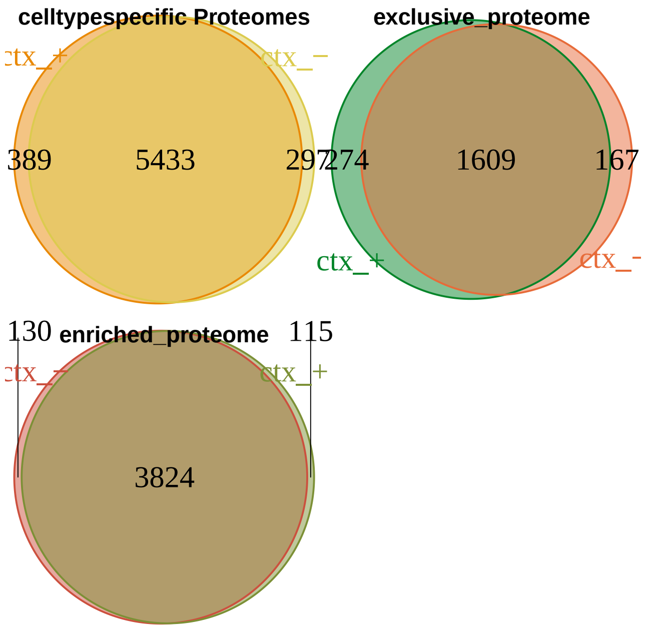


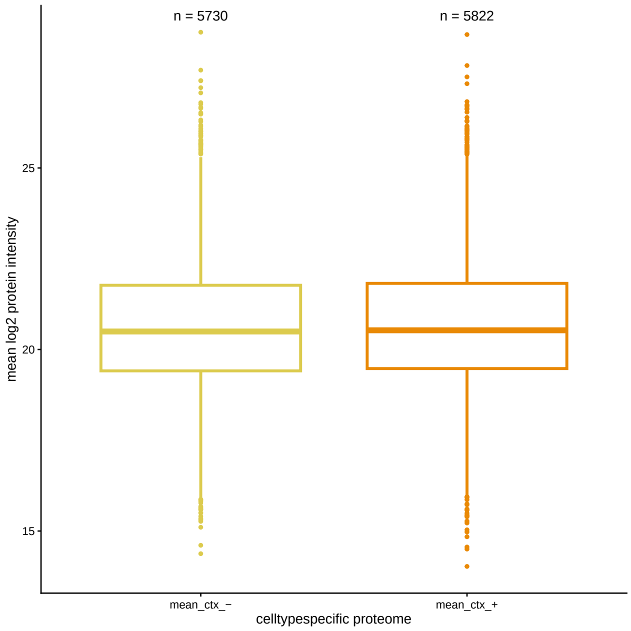


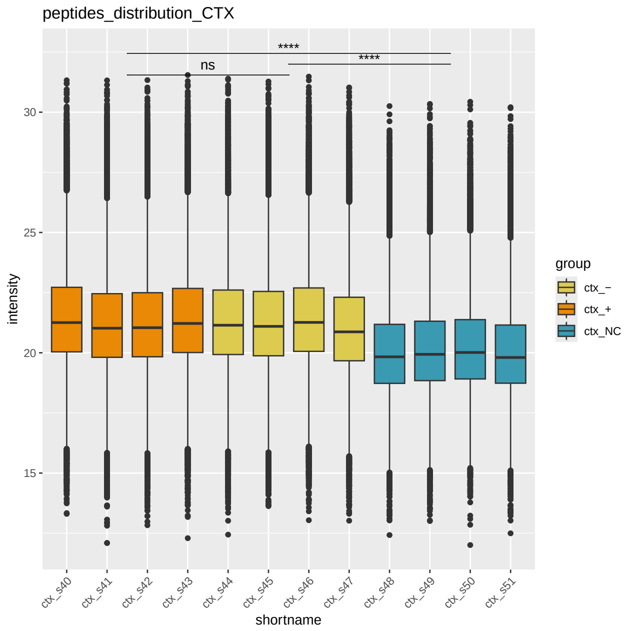

#### Glia, cortex, 7 months, continued

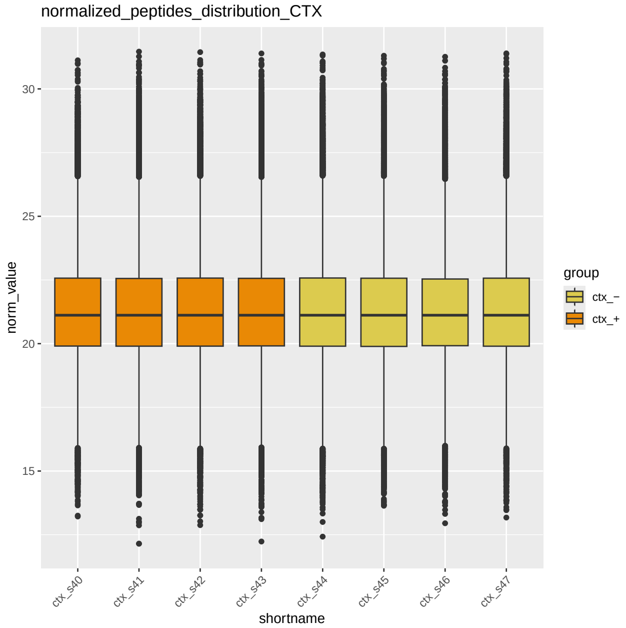


#### Glia, striatum, 7 months

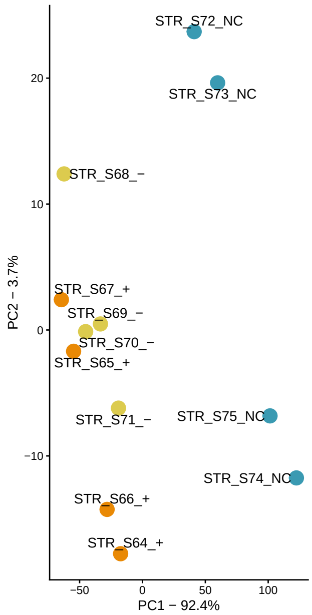


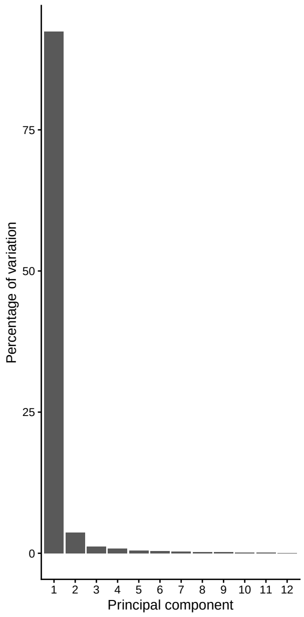


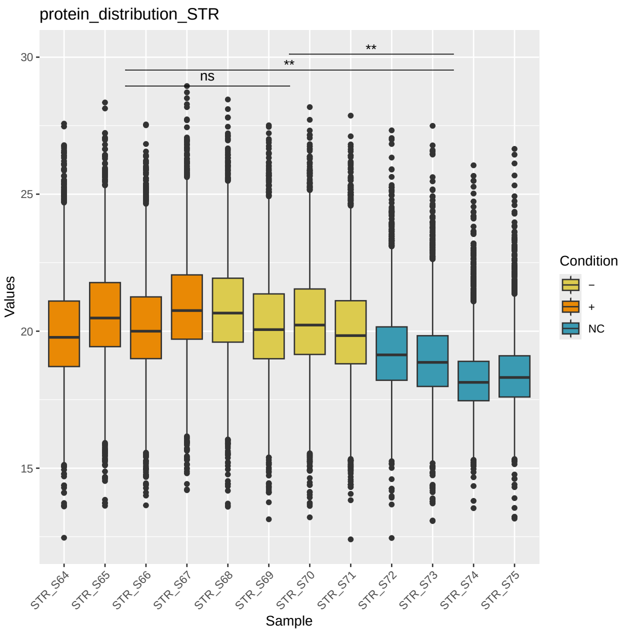


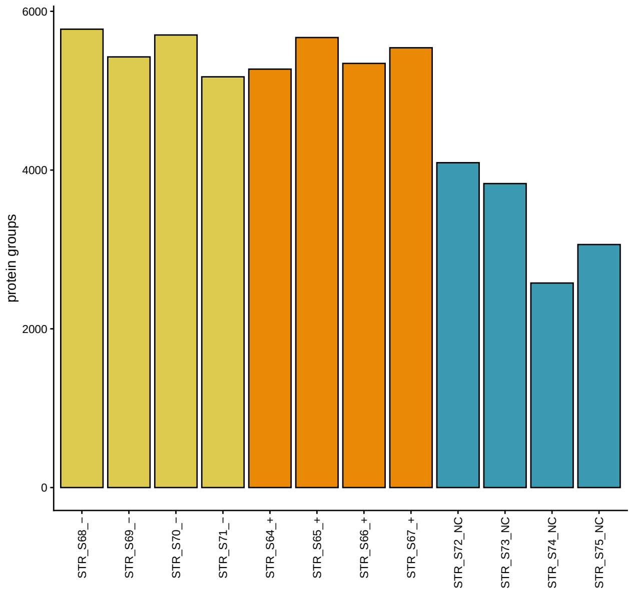


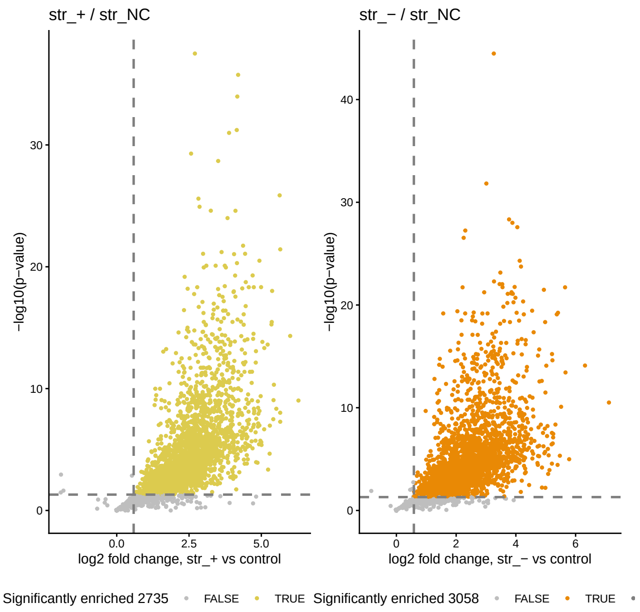

#### Glia, striatum, 7 months, continued

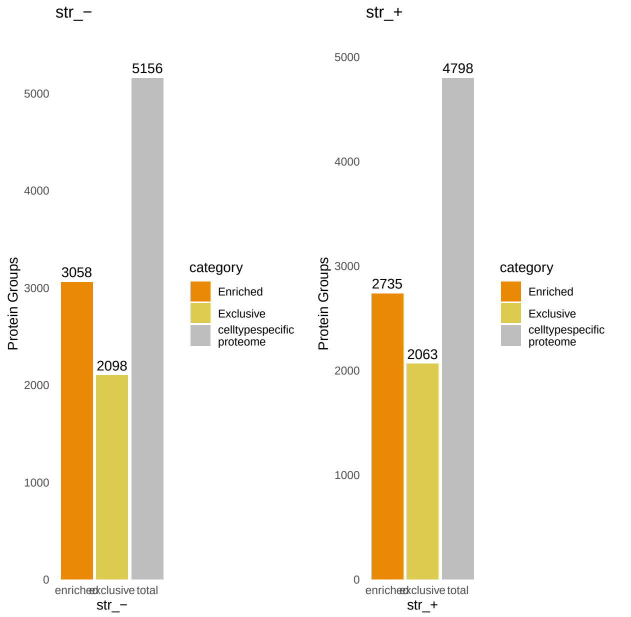


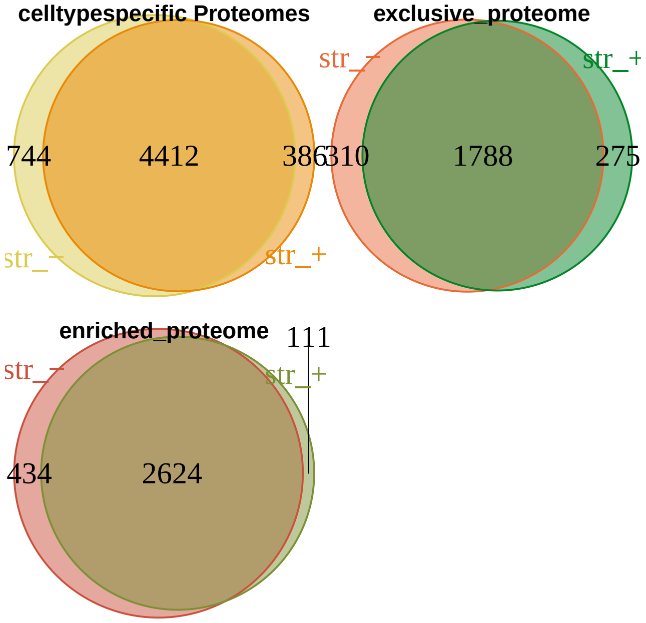


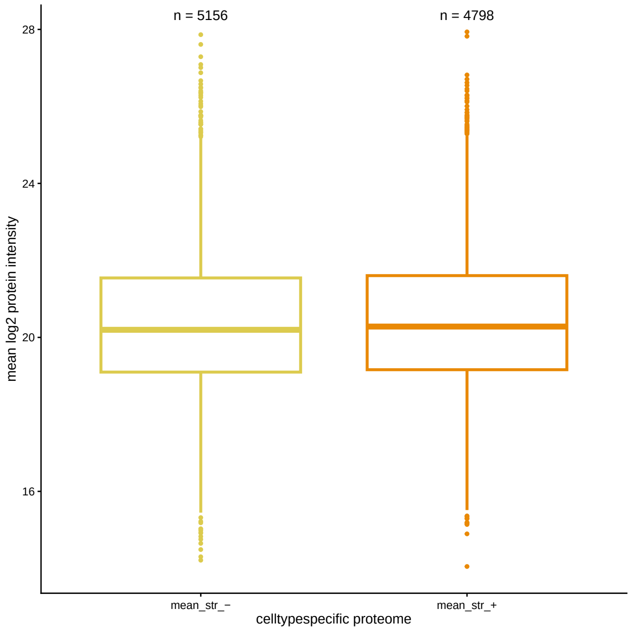


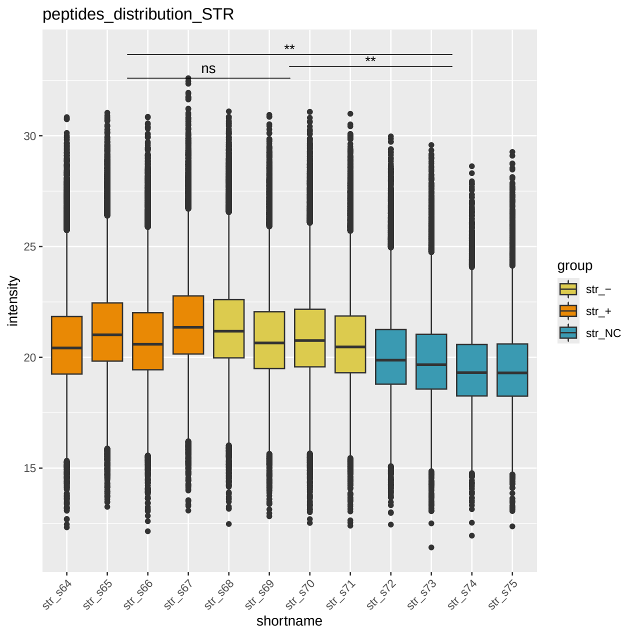


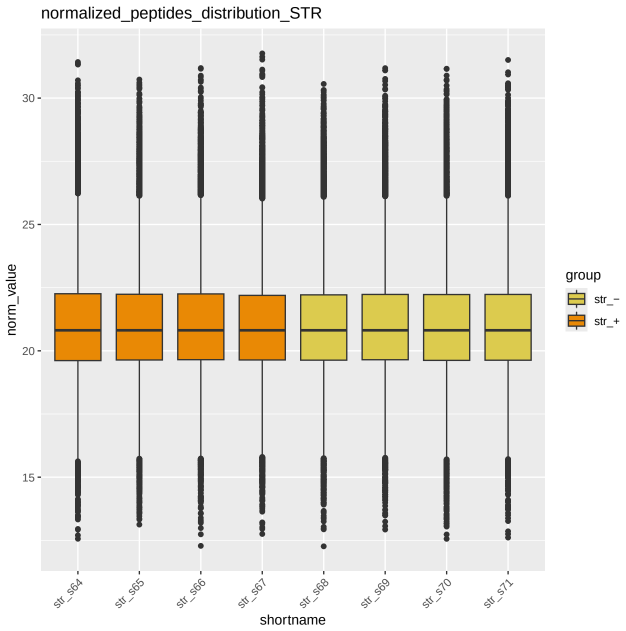

#### Glia, cerebellum, 7 months

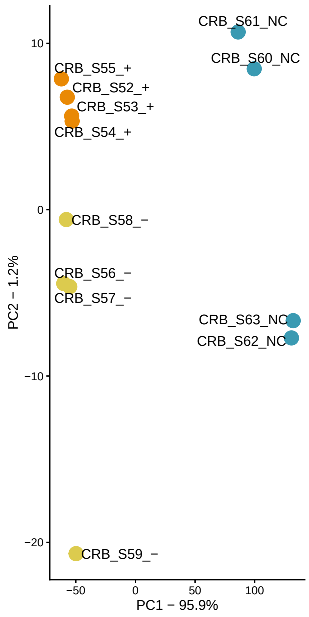


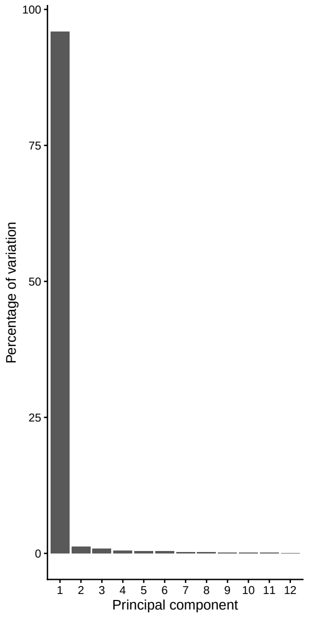


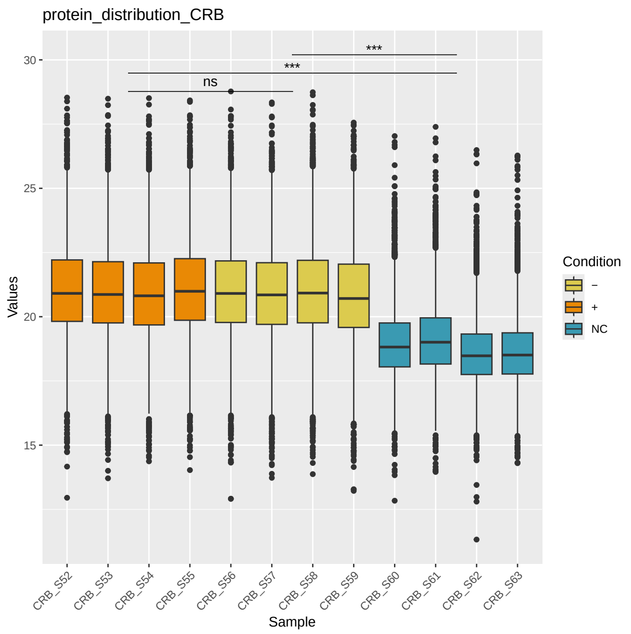


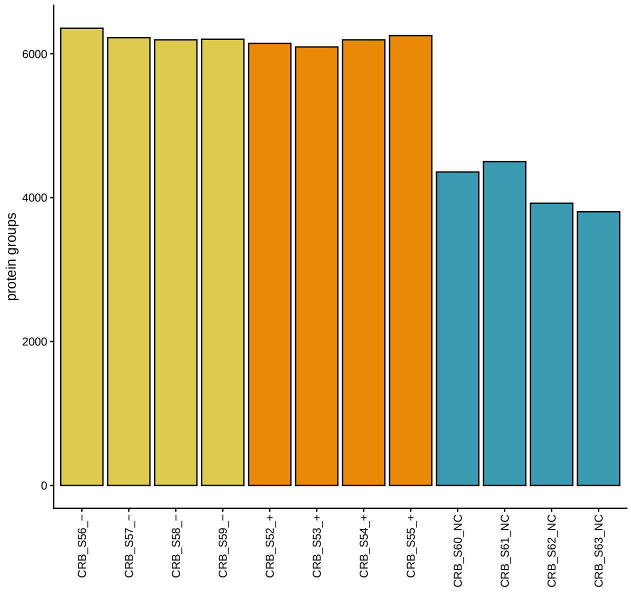


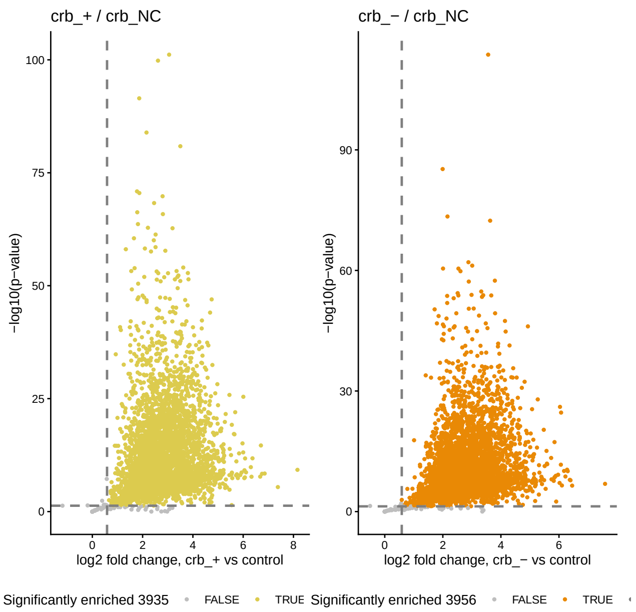


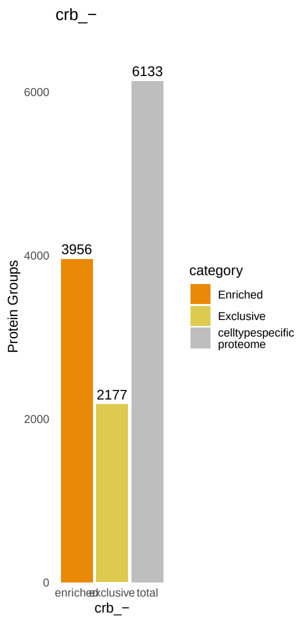


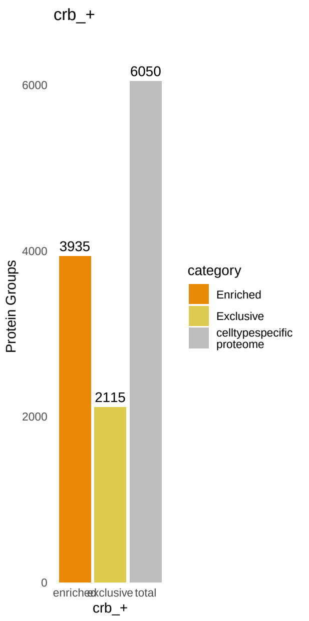


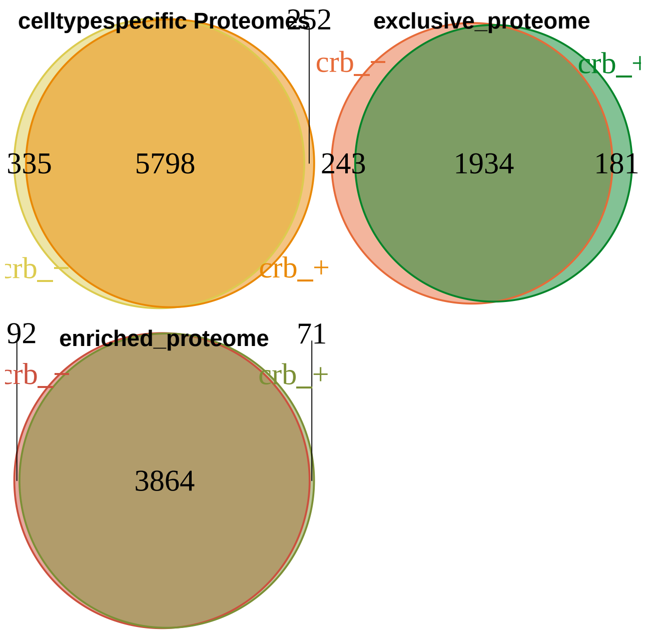

#### Glia, cerebellum, 7 months, continued

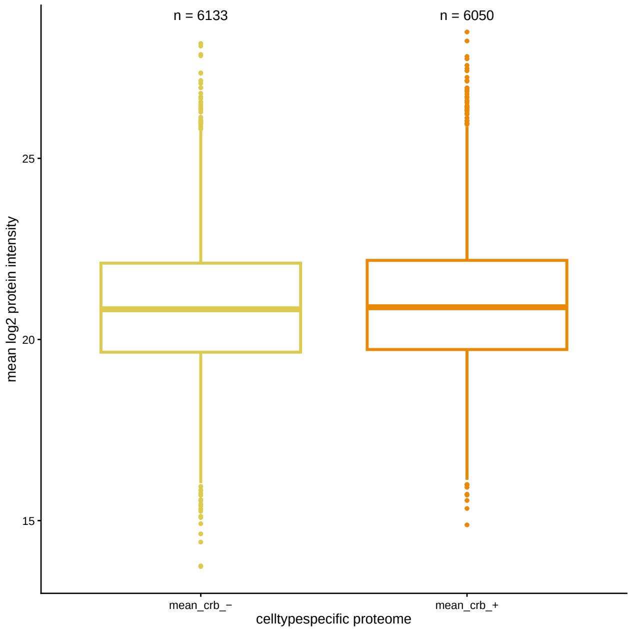


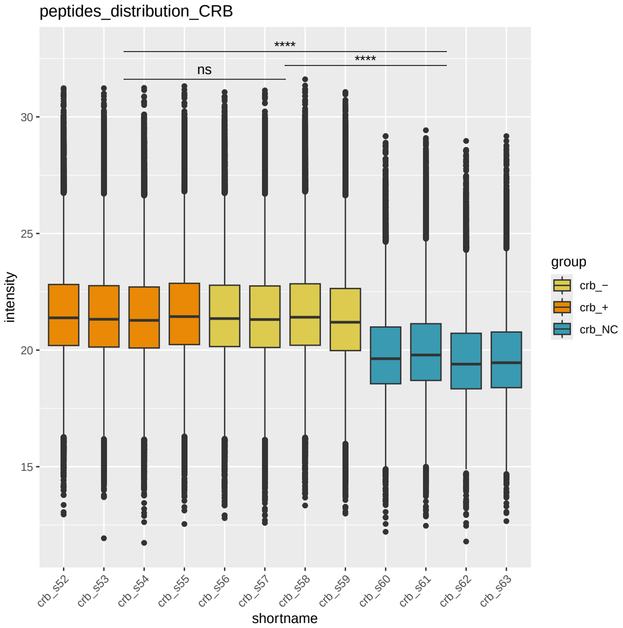


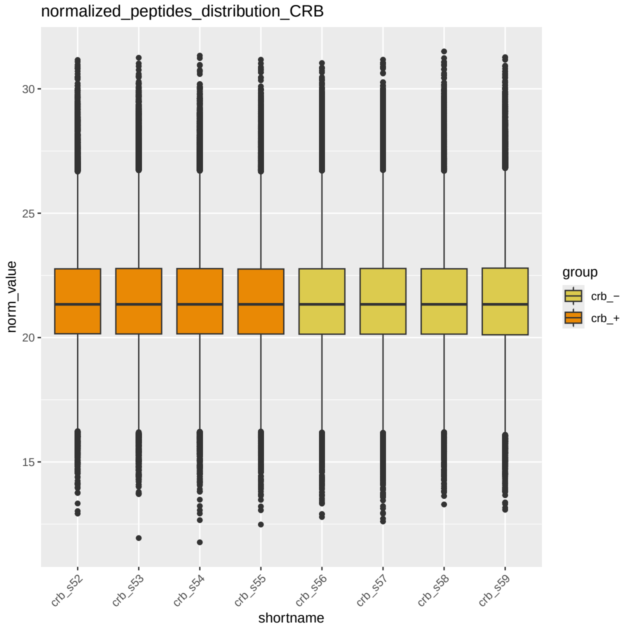
