## Supplementary material for "Excitatory neurons downregulate synaptic proteins before motor onset and glia reduce neuronal support in Huntington’s disease mice": ProteinHeatMAPS

**Supplementary File 2. Proteins of the protein sets of Fig. 9.**

(a) Antiviral response, viral restriction and interferon-responsive sets. (b) Proteasome, 20S core and 19S and accessory regulators. (c) Chaperones, DnaJ domain, protein refolding and CCT/TRiC. (d) Cytosolic ribosome. (e) Mitochondrial catabolism, TCA cycle and branched-chain amino acid catabolism on the first page and fatty-acid beta-oxidation in mitochondria on the second. Rows are the member proteins of each set and columns the comparisons of R6/1 with WT, as in Fig. 9, with glia at 7 months. Protein cells are coloured by the log2 fold change (R6/1/WT), as in the scale bar, and an asterisk marks the proteins regulated in R6/1. Dark red and dark blue cells with a triangle mark proteins upregulated in R6/1 (detected in R6/1 and not in WT) or downregulated in R6/1 (detected in WT and not in R6/1), all listed in Supplementary Table 1. These proteins have no fold change and did not enter the set test. Grey cells mark proteins not quantified or with no estimate in that comparison. The bottom rows give the cameraPR result of each set, as in Fig. 9, coloured by the signed −log10 FDR, red when the set is higher and blue when it is lower in R6/1, with an asterisk at FDR < 0.05. n.t., not tested. FDR, false discovery rate. CTX, cortex. HIP, hippocampus. CB, cerebellum. STR, striatum. m, months. n = 4 mice per genotype, except for cortical excitatory neurons (6 R6/1 and 6 WT at 1.5 months, 5 and 5 at 3 months, 5 R6/1 and 6 WT at 7 months).

**a**

### Antiviral and interferon-responsive astrocyte programmes increased in glia (the latter not in cortex), and in no neuronal contrast

Rows are proteins, ordered by mean fold change. In the interferon-responsive set, a name in brackets is the symbol Hasel et al. use.

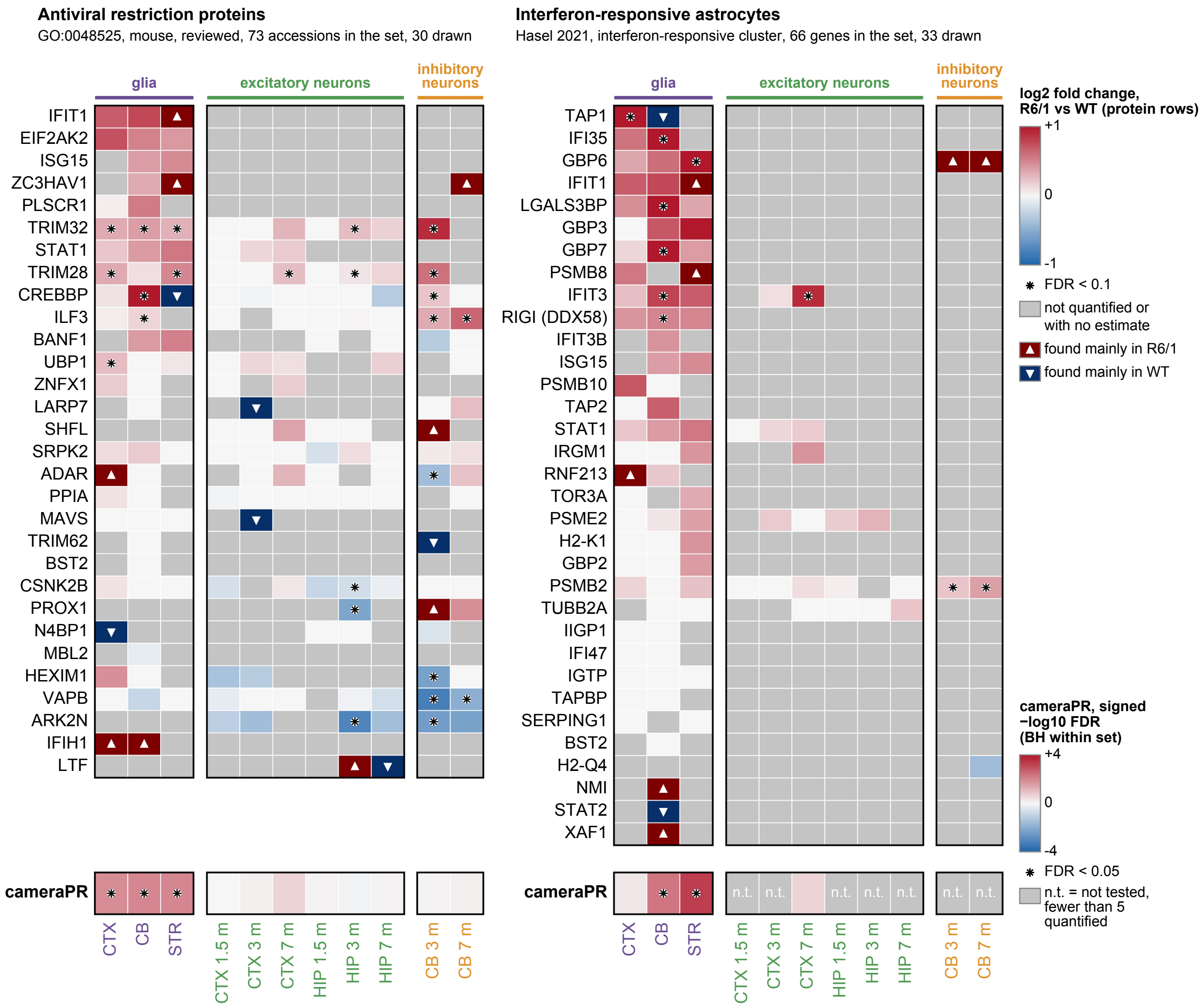

**b**

**Both proteasome sub-complexes increased in R6/1, earlier and more strongly in the 19S than the 20S**

Two lists - 20S core GO:0005839, 19 accessions in the set, 16 drawn, 19S accessory GO:0022624, 25 accessions in the set, 24 drawn

Rows are proteins, ordered by mean fold change, within each sub-complex.

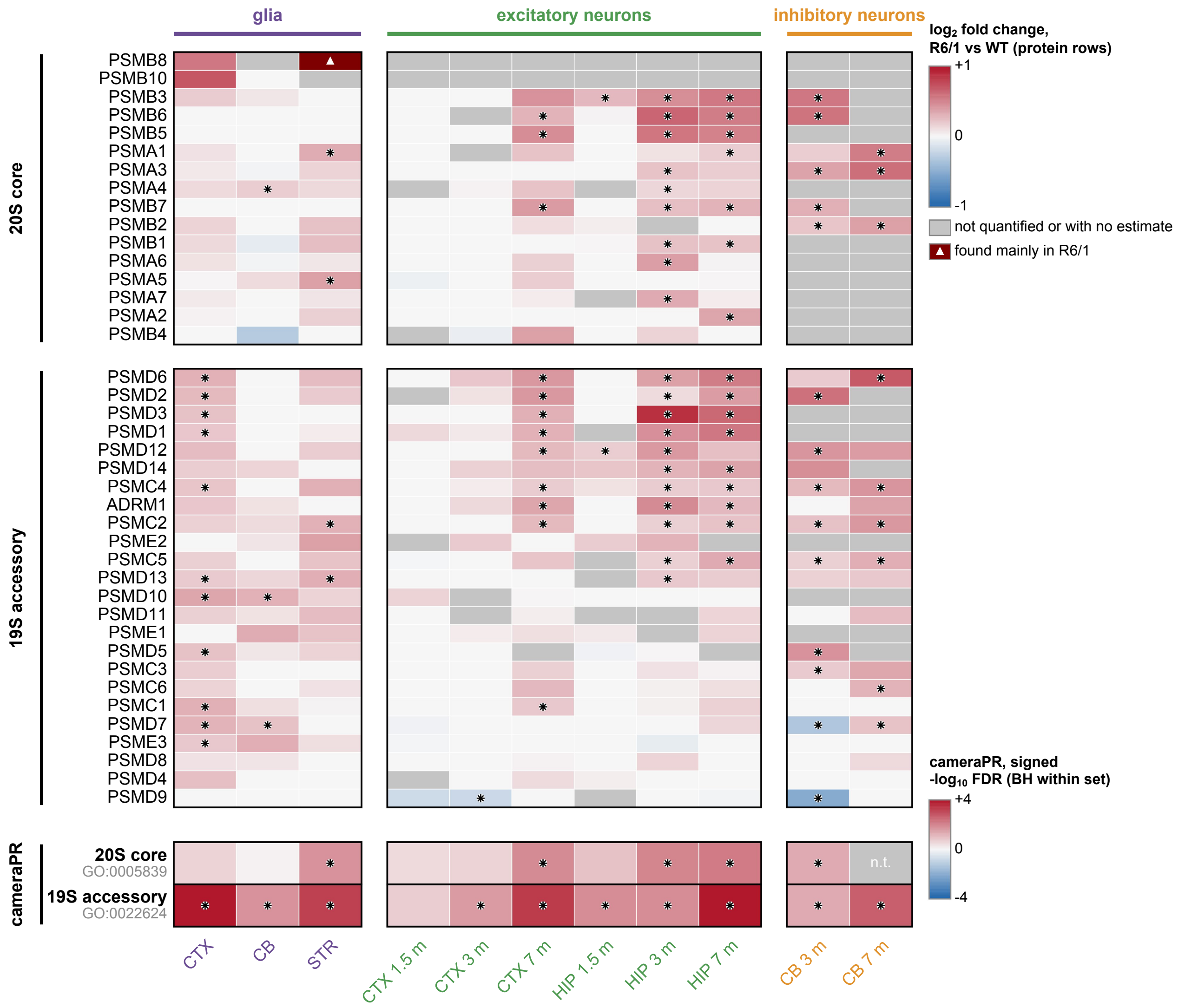

c

**The chaperone system splits in neurons, J-domain co-chaperones and refolding decreased, the chaperonin increased**

DnaJ, Pfam PF00226 42 accessions in the set, 28 drawn   refolding GO:0042026 22 accessions in the set, 17 drawn
chaperonin CCT/TRiC GO:0005832 9 accessions in the set, 9 drawn
Rows are proteins, ordered by mean fold change, within each arm.

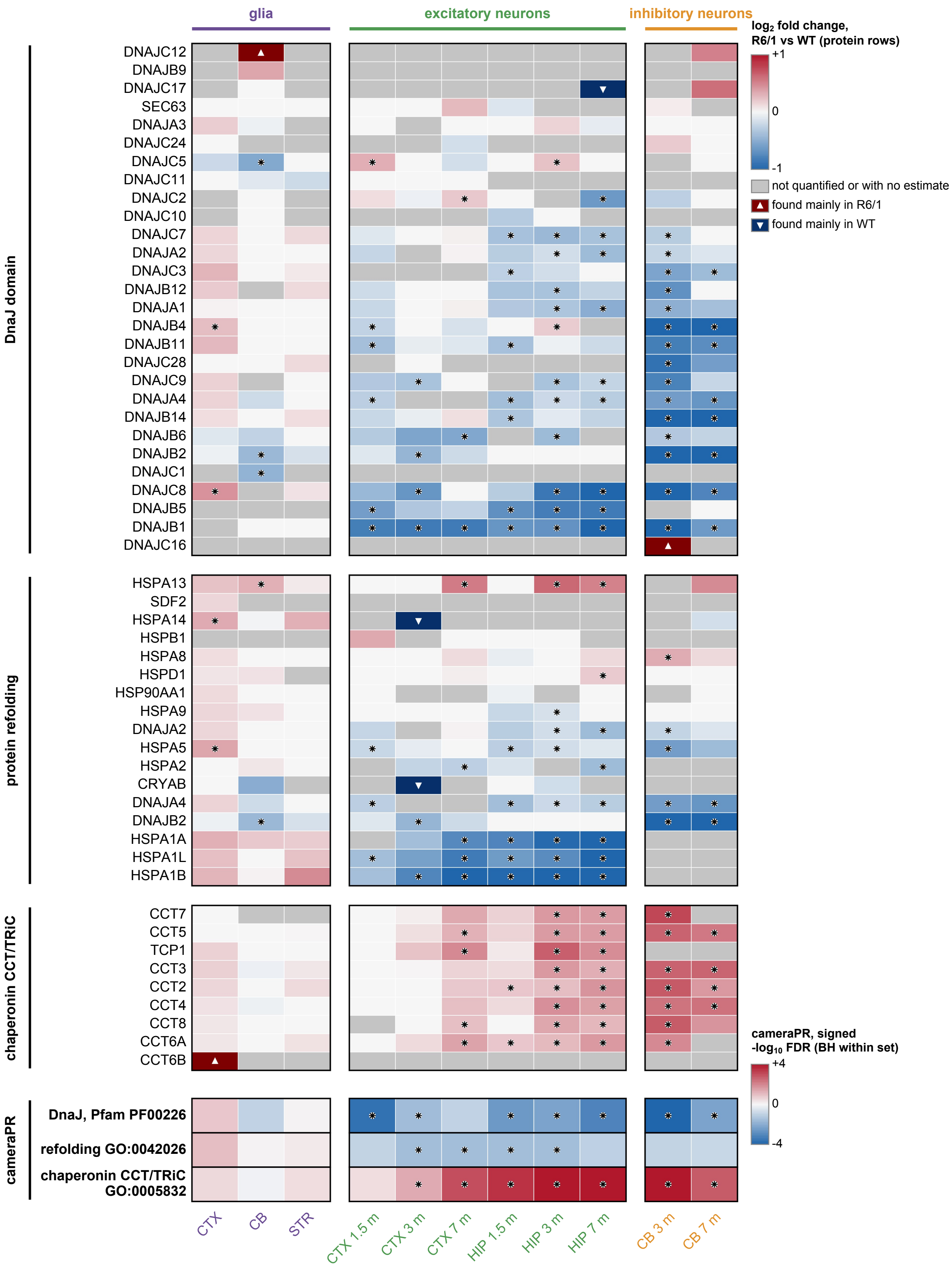

**d**

**Cytosolic ribosomal proteins increased at all three hippocampal ages and in cortex at 7 months**

Cytosolic ribosome, GO:0022626.

cytosolic ribosome 111 accessions in the set, 96 drawn

Rows are proteins, ordered by mean fold change.

Cerebellar glia show no significant set-level enrichment, CB glia FDR = 0.0840 (BH within set).

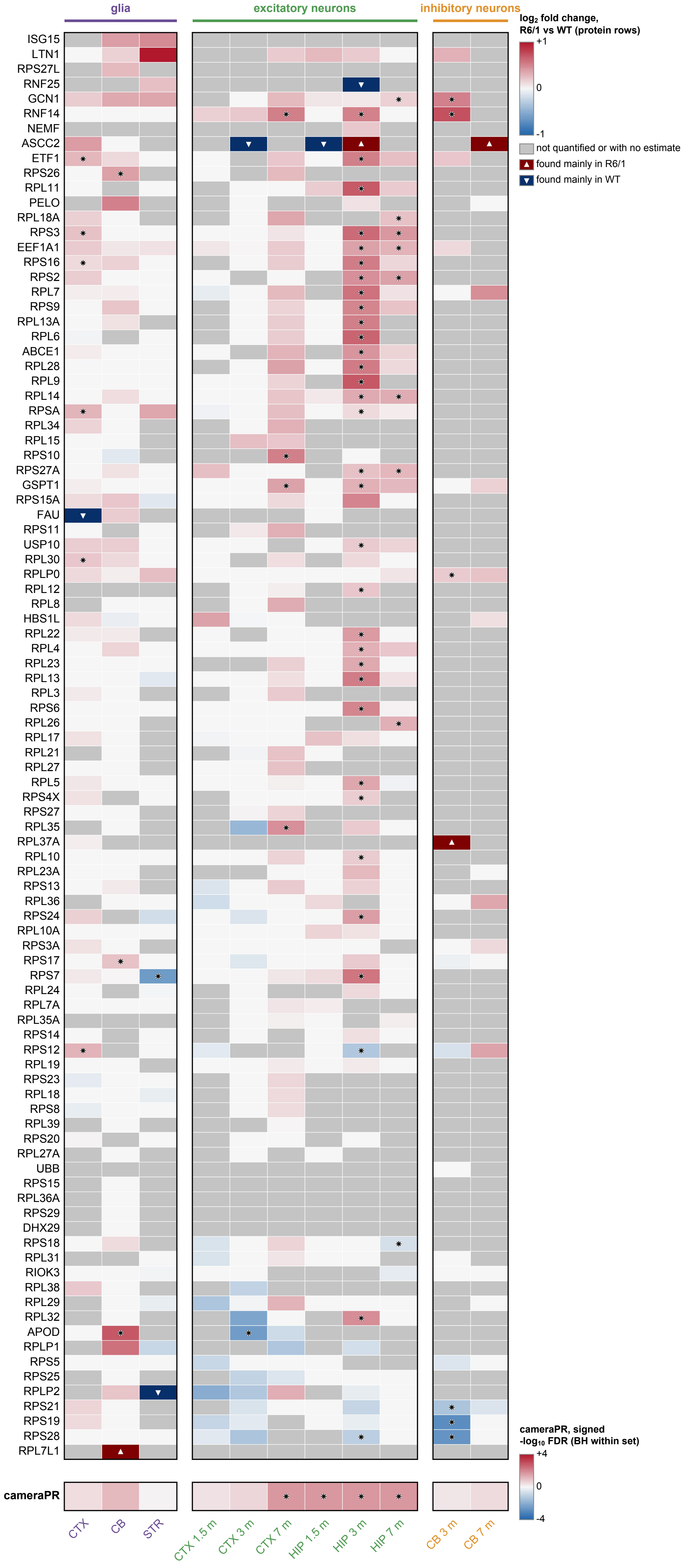

**e**

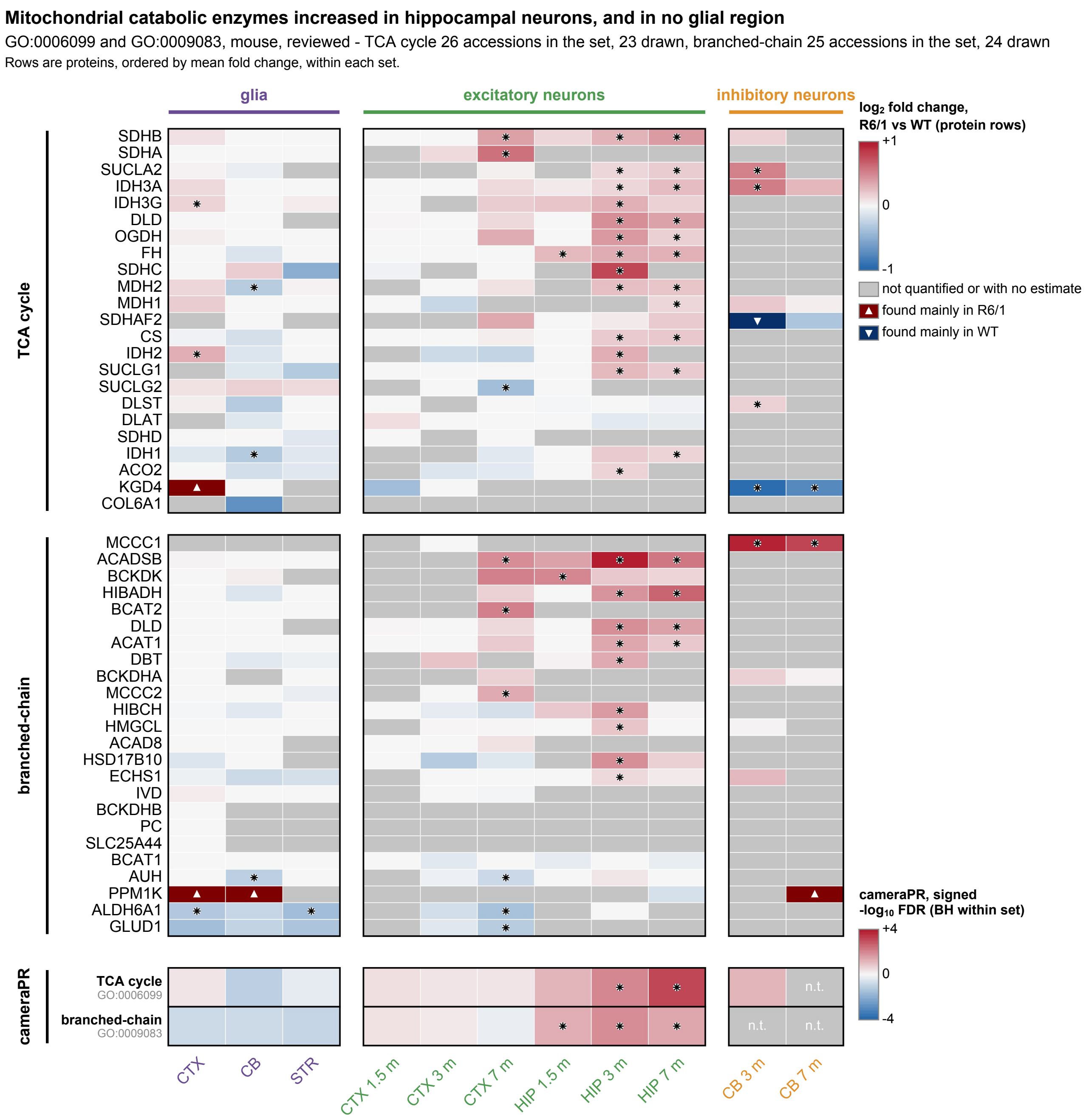

Mitochondrial catabolic enzymes increased in hippocampal neurons, and in no glial region
GO:0006099 and GO:0009083, mouse, reviewed - TCA cycle 26 accessions in the set, 23 drawn, branched-chain 25 accessions in the set, 24 drawn
Rows are proteins, ordered by mean fold change, within each set.
glia
excitatory neurons
inhibitory neurons
log2 fold change, R6/1 vs WT (protein rows)
+1
0
-1
not quantified or with no estimate
found mainly in R6/1
found mainly in WT
TCA cycle
SDHB
SDHA
SUCLA2
IDH3A
IDH3G
DLD
OGDH
FH
SDHC
MDH2
MDH1
SDHAF2
CS
IDH2
SUCLG1
SUCLG2
DLST
DLAT
SDHD
IDH1
ACO2
KGD4
COL6A1
branched-chain
MCCC1
ACADSB
BCKDK
HIBADH
BCAT2
DLD
ACAT1
DBT
BCKDHA
MCCC2
HIBCH
HMGCL
ACAD8
HSD17B10
ECHS1
IVD
BCKDHB
PC
SLC25A44
BCAT1
AUH
PPM1K
ALDH6A1
GLUD1
cameraPR, signed -log10 FDR (BH within set)
+4
0
-4
cameraPR
TCA cycle
GO:0006099
branched-chain
GO:0009083
n.t.
CTX
CB
STR
CTX 1.5 m
CTX 3 m
CTX 7 m
HIP 1.5 m
HIP 3 m
HIP 7 m
CB 3 m
CB 7 m

e

**Mitochondrial fatty-acid beta-oxidation increased in hippocampal neurons at every age, and in no glial region**

GO:0006635, mouse, reviewed, UniProt location mitochondrion SL-0173, downloaded 26 September 2026 - 33 accessions in the set, 29 drawn

Rows are proteins, ordered by mean fold change.

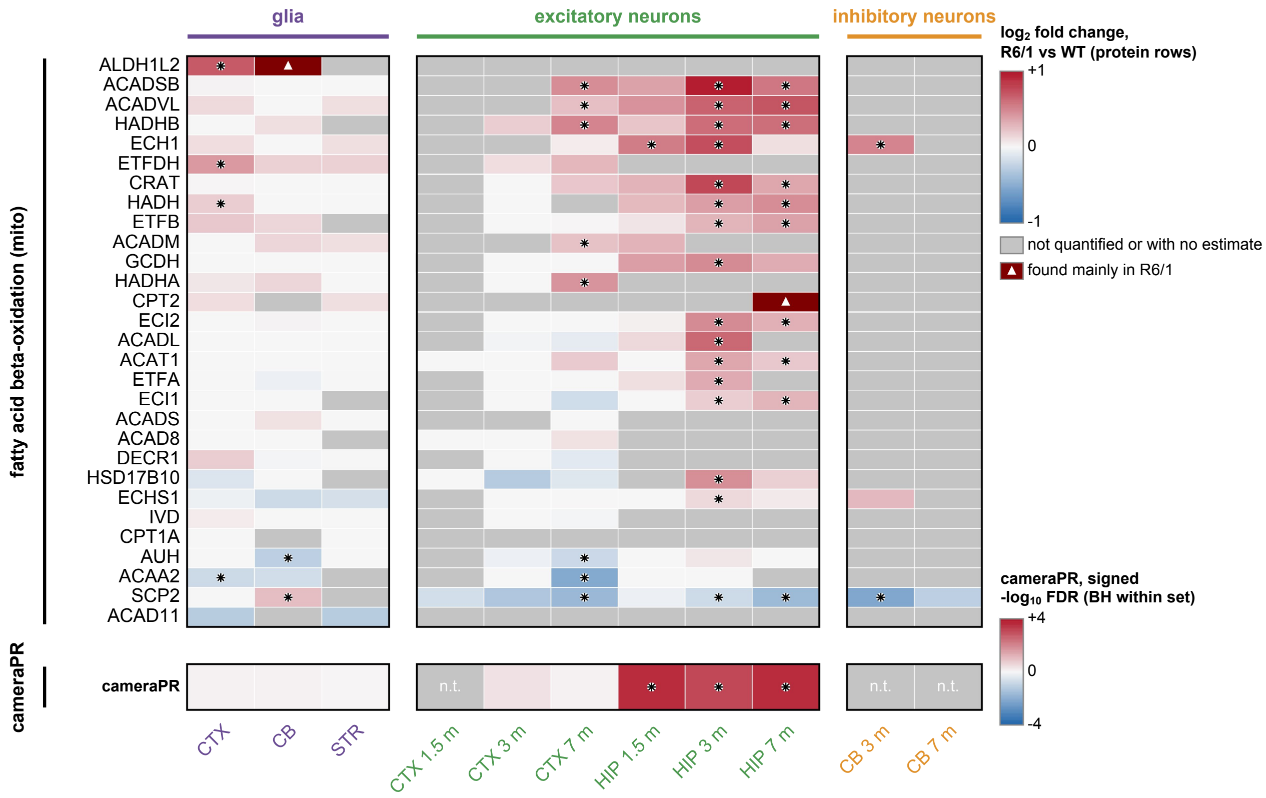
