## Supplementary material for "Excitatory neurons downregulate synaptic proteins before motor onset and glia reduce neuronal support in Huntington’s disease mice": Graphical_Abstract

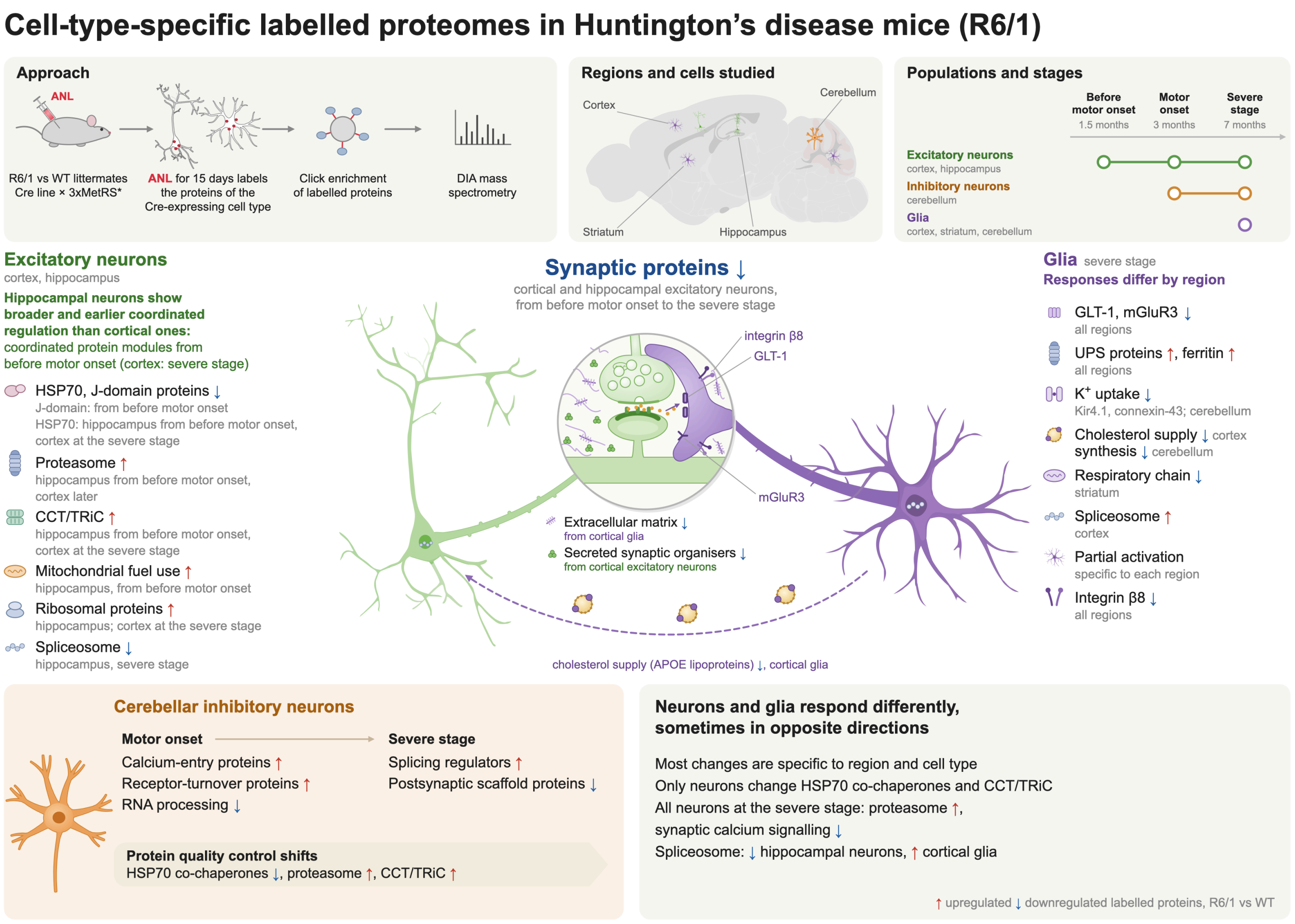


**Graphical abstract. Cell-type-specific labelled proteomes in R6/1 mice.** R6/1 mice and wild-type (WT) littermates expressing a mutant methionyl-tRNA synthetase (3xMetRS*) in one cell type (Cre-driven expression) received daily intraperitoneal injections of azidonorleucine (ANL) for 15 days. Only the cells expressing 3xMetRS* incorporate ANL into their proteins, and these labelled proteins were enriched by click chemistry and quantified by data-independent acquisition (DIA) mass spectrometry. Excitatory neurons were studied in cortex and hippocampus before motor onset (1.5 months), at motor onset (3 months) and at the severe stage (7 months), inhibitory neurons in cerebellum at motor onset and at the severe stage, and glia in cortex, striatum and cerebellum at the severe stage. The rest of the figure summarises the main changes in each population and how neurons and glia compare. Red and blue arrows mark labelled proteins upregulated or downregulated, respectively, in R6/1 compared with WT. At the magnified synapse, violet marks extracellular matrix proteins from glia and green marks secreted synaptic organisers from neurons. Drawings of the pyramidal neuron and mouse brain (Federico Claudi, DOI 10.5281/zenodo.3925905 and 10.5281/zenodo.3925911), CA1 pyramidal neuron (Stoyo Karamihalev, DOI 10.5281/zenodo.4312494) and astrocyte (Andrew Hardaway, DOI 10.5281/zenodo.5348394) were adapted from SciDraw (scidraw.io) under a CC BY 4.0 licence.
